# Self-organized mechanochemical instabilities drive the emergence of digit tissue morphogenesis

**DOI:** 10.1101/2025.08.31.673315

**Authors:** Rio Tsutsumi, Antoine Diez, Steffen Plunder, Ryuichi Kimura, Shinya Oki, Kaori Takizawa, Rei Nakano, Haruhiko Akiyama, Ritsuko Takada, Shinji Takada, Marco Musy, James Sharpe, Mototsugu Eiraku

## Abstract

How complex anatomical structures like digits emerge from an unstructured collection of cells remains a fundamental question in developmental biology. While Turing-type reaction-diffusion models explain the periodic pre-patterning of digits, the physical principles driving 3D morphogenesis remain incompletely understood. We develop a limb-mesenchymal organoid system that spontaneously forms elongated, digit-like protrusions, and use cellular-based computer simulations to identify sufficient microscopic mechanisms reproducing the tissue morphogenesis. We find that symmetry-breaking and elongation of digits arise from a combination of differential cell adhesion, morphogen-induced chemotaxis, and convergent extension. Coarse-graining the cellular-based model yields a continuum Partial Differential Equation description that couples a Cahn-Hilliard stabilization term with anisotropic diffusion, eventually leading to a “fingering instability” phenomenon mathematically parallel to that known in fluid physics. Our findings suggest that mechanochemical fingering instability may act in concert with chemical patterning to drive vertebrate digit morphogenesis.

## Main

The morphology of anatomical structures has evolved to execute specialized functions with remarkable efficiency. A prime example is the distal end of the vertebrate limb, which consists of a modular design of repetitive elements that form the digits, enabling complex tasks such as goal-directed manipulation. Although these structures vary in shape, size, and number across species, their core design principles —governed by molecular and cellular behaviors during development— are fundamentally conserved^1^.

Despite our detailed knowledge of the molecular and cellular behaviors involved, a fundamental question remains: how do microscopic interactions translate into the macroscopic shape of tissues^2^? This underscores the emergent nature of morphogenesis, where tissue-level structures cannot be predicted solely from a list of individual components^3^. In physics, similar problems have been addressed through statistical physics frameworks that bridge microscopic interaction rules and macroscopic laws. Whether tissue morphogenesis can be understood in an analogous way is an open and challenging question that requires resolving the complexity of biological systems into analytically tractable multiscale mathematical models.

In this context, a key concept is the idea of instability, which postulates that complex feedback loops can amplify small microscopic perturbations to produce macroscopic structures. One of the most influential instabilities linking molecular behaviors and tissue-scale pattern formation in biology is the Turing instability, driven by biochemical reaction–diffusion systems^4^. It effectively explains early patterning of digit and interdigit primordial tissues (the latter subsequently undergoes apoptosis) in the mouse limb bud (E10.5), where patterns emerge prior to significant cell movement^5,6^. However, while sufficient for initial establishment of molecular periodic patterning in the limb, this framework fails to account for the tissue-scale morphogenetic movements that physically shape digit protrusions along the distal–proximal axis by E12.5. The morphogenesis at later stages should depend on mechanochemical cell–cell interactions^7^, yet whether macroscopic morphogenesis can emerge from microscopic cellular behaviours through a mechanochemical instability remains an open question. To address this, we seek to identify the minimal biomechanical and biophysical “design principles” essential for digit formation.

Multicellular in vitro systems, particularly organoids, provide a powerful platform to study such questions, as they can self-organize and give rise to multicellular patterns and structures under defined controllable conditions, thereby bypassing much of the complexity of in vivo development^8–11^. Although *in vitro* derivation of the limb mesenchyme has been reported^12–17^, evidence of morphogenesis into identifiable limb anatomical structures has not been observed. Thus, we establish a minimal physical explanatory organoid model isolating the intrinsic morphogenetic behaviors of digit tissues using embryonic limb mesenchyme and combine it with agent-based and continuum mathematical models to identify the minimal design principles sufficient for digit formation. While Turing-type reaction–diffusion models explain molecular pre-patterning along the anterior–posterior axis within a given tissue geometry, our study addresses a distinct class of instability that underlies deformation of tissue geometry itself along the distal– proximal axis during digit morphogenesis. Here we show that, in a limb-mesenchymal organoid system and experimentally grounded modeling framework, this tissue deformation is driven by a mechanochemical fingering instability that emerges from cellular interaction rules and is mathematically parallel to classical fingering phenomena in fluid physics.

## Results

### Fgf8b and Wnt3a signaling are sufficient to induce digit-like structures in limb mesenchymal organoids

To identify the fundamental design principles required for digit morphogenesis of the hand, we first dissociated and cultured 9,000 limb mesenchyme cells derived from autopods of Sox9-GFP (to visualize cartilage differentiation) mouse embryos at E11.5, which formed spherical cellular aggregates in a three-dimensional (3D) culture conditions (Figure 1A). Aggregates cultured under serum-only conditions (negative control: NC) largely maintained their spherical shape, but displayed non-uniform Sox9 expression (Figure 1B: left panel, Movie 1). Fgf8b and Wnt3a are known to be secreted from the apical ectodermal ridge (AER) and the overlying ectoderm, respectively, and act synergistically to induce distal identity and growth of limb mesenchyme morphogenesis^18–20^. Indeed, as the supplementation of Fgf8b/Wnt3a (FW) induces the elongation of limb mesenchymal-organoid from E12.5 embryos^13^, we next investigated the effect of FW on cultured cellular aggregates from limb autopod mesenchyme from E11.5 embryos. We observed that FW cultured aggregates not only went beyond non-uniform Sox9 distribution, but also broke their *shape-symmetry* growing into stick-shaped cartilage elongations, despite the fact that we supplemented FW uniformly in the medium (Figure 1B: middle panel). Interestingly, when cultured with a larger cellular aggregate (27,000 cells), we observed the formation of multiple bifurcated cartilage rather than an isotropic scale up (Figure 1B: right panel, 1C). This indicates that FW is sufficient to initiate not only shape-symmetry breaking, but also the physical elongation of cartilage without any directional cues in the medium (Figure 1D).

**Figure 1.**
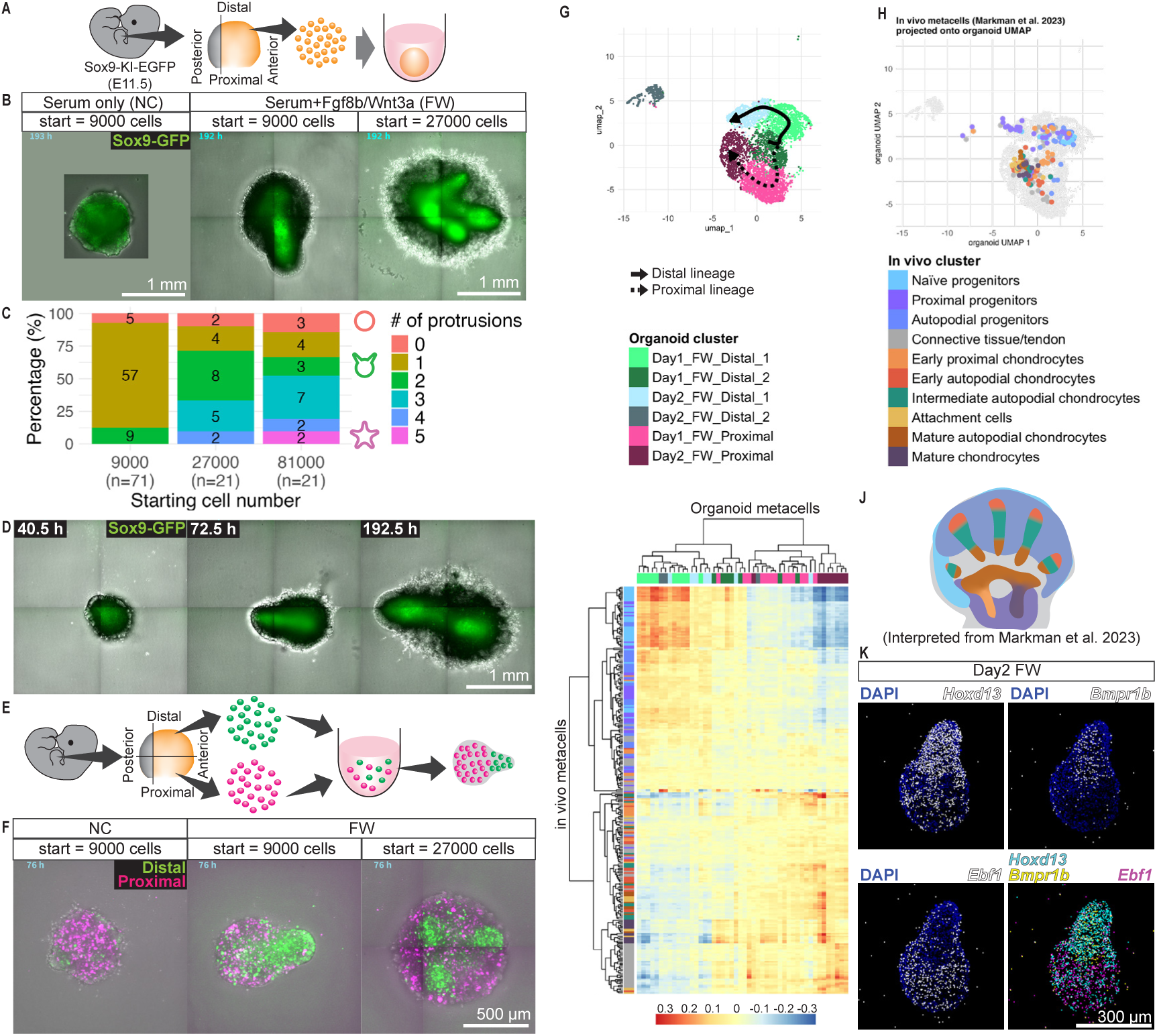
Formation of digit-like structures in limb-mesenchymal organoids. (A) Schematic of organoid generation from limb mesenchyme of Sox9-KI-EGFP E11.5 mouse embryos. (B) Endpoint morphology at 193 hours in serum-only (NC) or Fgf8b/Wnt3a-supplemented medium (FW), initiated with 9,000 or 27,000 cells. Sox9-GFP, green; Bright-field, gray. (C) Distribution of protrusion numbers by starting cell number. Stacked bars show percentages; labels indicate raw counts of organoids. Association was tested by batch-stratified Cochran–Mantel–Haenszel test using two eligible batches: M² = 28.07, df = 10, P = 0.00176. (D) Time-lapse imaging of FW organoids initiated with 9,000 cells. (E and F) The distal (H2B-Venus, green)–proximal (H2B-tdTomato, magenta) cell-labeling strategy and representative organoids. (G) UMAP of day-1 and day-2 FW organoid scRNA-seq profiles with lineage trajectories. (H) *In vivo* limb metacells from Markman et al. 2023 projected onto the organoid UMAP. (I) Correlation between expression profiles of organoid and *in vivo* metacells. (J) Spatial distribution interpreted from Markman et al. 2023. Color codings of organoid clusters in G and I, and *in vivo* clusters in H-J are common. (K) Expressions of *Hoxd13*, *Bmpr1b*, and *Ebf1* in FW organoids; HybISS spots are shown as filled circles (Original images: figure S1I).

To determine the mechanistic basis of symmetry-breaking in our aggregates, we first hypothesized that it could be based on the cell sorting between two cell populations —the cells originated from distal (digit) and proximal (palm) parts of the autopod. We dissected and labelled distal and proximal cells with H2BVenus and H2BtdTomato, respectively and cultured the aggregates of the two cells under NC and FW conditions (Figure 1E). Although distal and proximal cells formed an intermixed cellular aggregate under NC condition, they underwent sorting in FW conditions within 24 hrs, with the distal cell cluster subsequently forming an elongating tip region (Figure 1F: left and middle panels, Movie 2). This suggests that shape asymmetry is dependent on cell sorting. Interestingly, when more cells were used as starting material (27,000 cells), distal cells tended to cluster at multiple sites, indicating the bifurcation of cartilage (Figure 1F: right panel).

Next, we performed a high-depth transcriptome analysis for a locally defined area via photo-isolation chemistry (PIC)^21^, to identify the genes highly upregulated at the tip of the organoids. Combined with our scRNA-seq analysis the results showed not only that the tips of organoids corresponded well with the genes expressed in the distal digits, including *Hoxd13*, *Msx1*, *Lhx2*, and *Sox11*^22–25^ (Figure S1A), but also suggested that FW maintains distal and proximal identities and naïve developmental state, while NC drives both distal and proximal cells towards proximal identity and into a differentiated state, consistent with the role of AER-derived signals in sustaining distal limb mesenchyme (Supplementary Text: scRNA-seq, Figure S1 and S2).

The pseudo-time analysis inferred the progression of expression profiles in distal cells and proximal cells (Figure 1G). Subsequently, we compared the transcriptomic profiles of the organoids and *in vivo* limb mesenchyme by projection of metacells from *in vivo* limb buds^26^ onto the same UMAP space (Figure 1H) as well as the hierarchical clustering heatmap showing the correlation of gene expression profiles between the organoids and *in vivo* metacells (Figure 1I). The results suggested that the distal cells in organoids undergo the consistent developmental progression of the limb mesenchyme of *in vivo* digits, from naïve to autopodial, and proximal progenitors. In contrast, the proximal cells showed more advanced progressions from early, intermediate and mature chondrocytes, implying the correspondence of the distal and proximal cells in organoids to the digit and more proximal elements, such as metacarpals and carpals, respectively (Figure 1J)^26^. Furthermore, the hybridization-based in situ sequencing (HybISS)^27^ showed that the distal part of the organoids expressed the distal autopod marker *Hoxd13,* as well as the digit marker gene *Bmpr1b*, while the proximal part expressed the proximal marker *Ebf1* (Figure 1K)^22,28,29^. Therefore, based on our cell lineage and expression profile data, these suggest that FW is sufficient to trigger the emergence of a digit-like morphological structure in our 3D limb-mesenchymal organoid system, thereby capturing core morphogenetic behaviors underlying digit formation *in vivo*.

To analyze the contribution of cell sorting to the patterning of Sox9 expression in quantitative manners, we also cultured H2BtdTomato-labelled distal and H2BtagBFP-labelled proximal cells, from the Sox9-GFP mouse embryos on two-dimensional (2D) micromass culture (Figure S3A). Similar to our findings in limb-mesenchymal organoids, we observed that FW conditions promoted cell sorting between the distal and proximal cells in comparison to NC conditions (Figure S3B, Movie 3). Interestingly, when spots of Sox9 expression emerged, they typically appeared to be associated with either sorted clusters of distal cells or proximal cells (Figure S3B). This is consistent with limb development *in vivo*, as the molecular pre-patterning of Sox9 corresponds not only to distal digital elements, but also the more proximal metacarpal and carpal skeletal elements. Strikingly, although Sox9 spots could form in distal clusters or proximal clusters, they appear less likely to emerge in regions with strong intermixing of proximal and distal (or on the border between the cluster), indicating that the reaction-diffusion mechanism of molecular patterning operates strongest between cells of the same proximo-distal identity.

To quantify the level of cell sorting, we computed the nearest neighbor score (NNS) by counting the number of the same type of cells (distal or proximal) and normalizing it such that the score would be one under the null hypothesis (random distribution), while the score would be higher when cell sorting occurs (See methods) (Figure S3C). This confirmed that the proportion of distal cells with at least nine distal neighbours (out of 10 neighbouring cells) increased over time, and those cells exhibited higher Sox9-GFP intensities. This effect was weaker for proximal cells, but nevertheless consistent: over time the more clustered proximal cells increase their expression of Sox9 (Figure S3D, E). These results suggest that under FW conditions, the cell sorting between distal and proximal cells has a strong influence on the spatial patterning of cartilage (Figure S3D,E, S4A). In contrast, when only distal cells were cultured in NC condition, labyrinth-like patterns were created, and the intervals between these nodules were further extended under FW conditions, which has been attributed to Fgf wavelength modification (Figure S4B, C)^5,6,30^.

### Cell adhesion and chemotaxis drive symmetry-breaking but not digit-like structure elongation

Considering that dynamic cell movements are required for symmetry-breaking and elongation observed in our 3D limb-mesenchymal organoids, we next built an Agent (cellular)-Based Model (ABM). Cell sorting is typically achieved through differential cell adhesion. Indeed, limb-mesenchymal cells at different positions and ages have different cell adhesiveness^31^. For example, EphA7 is known to regulate the cell sorting between distal and proximal autopod mesenchyme at E11.5 mouse embryos^32^. Cell adhesiveness can be parametrized as the contact angle between cell doublets, whereby cells with stronger adhesion have larger angles and vice versa ^33^ (Figure 2A, left schematic). Thus, we cultured aggregates of distal and proximal cells separately under NC and FW conditions, dissociated them on day 1 and day 2 to make doublets, and measured the angles between cells (Figure S5A). We found that cell adhesions between distal-distal cells were stronger than distal-proximal and proximal-proximal on day 1, and that the distal-proximal adhesion became equally strong as distal-distal on day 2 when cultured in FW, while no differences in cell adhesions were observed when cultured in NC (Figure 2A, S5B).

**Figure 2.**
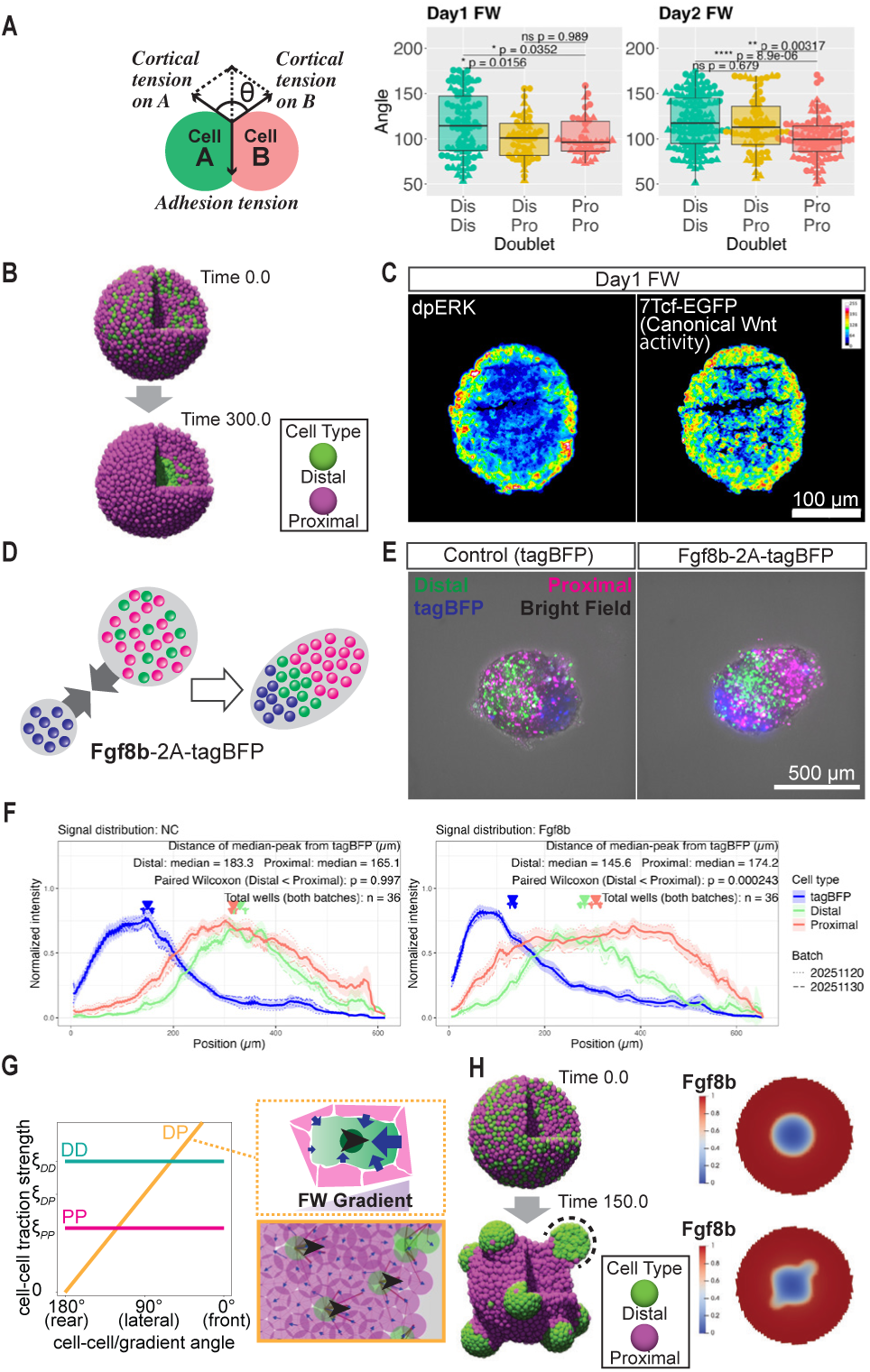
Differential adhesion plus Fgf8b-directed traction bias explains symmetry breaking. (A) Beeswarm+box plots of Angle by doublet class (DD, DP, PP) for Day1/Day2 FW. Dots = doublets; shape = batch. Stats: one-way ANOVA per experiment; if significant (α=0.05), Tukey HSD shown. (B) Agent-Based Model (ABM) implementing differential adhesion only (DD>DP>PP) (C) Day-1 FW immunostaining: dpERK and 7Tcf-EGFP (red→blue). (D) Assembly assay: a proximal signaling aggregate (tagBFP or Fgf8b-P2A-tagBFP) assembled with a Distal+Proximal aggregate. (E) Live imaging at 96 h. (F) Signal distribution at 96 h. Intensities for tagBFP, H2B-Venus (distal), and H2B-tdTomato (proximal) were integrated along a line from the tagBFP region; intensity-weighted medians defined peaks. Distances from the tagBFP peak to Distal vs Proximal peaks were compared by paired one-sided Wilcoxon (hypothesis: Distal closer). Plots show mean normalized profiles ±95% CI with median markers. (G) Rule map: angle to the Fgf8b gradient (x-axis) vs traction strength (y-axis). DD (green, strong) and PP (magenta, weak) are constant; DP (yellow) is angle-dependent, biasing toward Fgf8b gradient. Dotted box: DP vectors (blue arrows) drive Distal motion (black arrowhead); Solid box: predicted Distal motion toward Fgf8b. (H) ABM implementing differential adhesion + traction bias toward the Fgf8b gradient. Right: Fgf8b field.

Based on these results, we constructed a minimal representative ABM to test whether differential adhesion is sufficient to explain the initial symmetry-breaking of distal and proximal cells. By incorporating differential cell adhesion, whereby distal-distal adhesion is stronger than the distal-proximal or proximal-proximal, we found that distal cells segregate from proximal cells, but cluster at the centre of the organoid rather than at the surface (Figure 2B, S5C: Dotted box, Movie 4). This indicates mechanistically that differential cell adhesion alone is sufficient to drive cell sorting, but insufficient for distal cells to cluster on the surface of our organoids and that an additional mechanism is likely to be involved.

As exposure to FW in the culture medium may lead to an inherent bias in signalling cues received between the surface and centre of the organoid, we investigated the activation of the downstream signalling pathways in response to FW within the organoids. We performed immunostaining against phosphorylated-ERK and electroporated 7Tcf-GFP, to visualize the activation of Fgf signalling in response to Fgf8b and canonical Wnt signalling in response to Wnt3a, respectively. Indeed, in both cases, we observed that cells at the surface of the organoid had stronger signal activation than at the centre (Figure 2C), indicating FW gradient is inherently established within the organoid towards its surface. Given this spatial bias, we next investigated the chemotactic behaviour of distal cells towards FW. We generated limb-mesenchymal organoids that included small aggregates of proximal cells which overexpressed either Fgf8b-T2A-tagBFP or tagBFP, as a control (Figure 2D). As expected, after culturing under NC conditions, distal cells moved to the close proximity of the Fgf8b-T2A-tagBFP overexpressing cells, but not to tagBFP overexpressing cells (Figure 2E, F, Movie 6). Importantly, chemotactic activity was not observed when known morphogens expressed in limb mesenchyme, such as Wnt5a or Inhba (encoding activin), were overexpressed in these small aggregates (Figure S5D, E). These results showed that distal cells undergo chemotaxis along the Fgf8b gradient.

Based on the experimental evidence for the inner-outer gradient of FW signaling within the organoid and Fgf8b-directed chemotaxis of distal cells, we next tested whether adding chemotactic bias to the ABM could explain surface localization of distal clusters and subsequent digit elongation. Here, we modelled cell motion solely with reciprocal pulling/pushing mechanisms between pairs of cells that preserve the symmetry of Newton’s third law^34^. For directed cell migration towards the gradient in the ABM, in addition to the stronger adhesion strength of distal cells, we also implemented that distal cells have a directional traction bias. This was defined such that when distal cells are adjacent to proximal cells, they pull the proximal cells and propel themselves strongly towards the chemotactic gradient (Fgf8b, in this case) (Figure 2G). Using such parameters, the ABM was able to recapitulate the morphological symmetry-breaking in limb-mesenchymal organoids, but not elongation beyond spherical clusters of distal cells located at the surface (Figure 2H, dotted line, S5C: Solid box, Movie 5). Thus, differential adhesion and Fgf8b-directed chemotactic bias are sufficient for symmetry-breaking, but not for sustained digit-like tissue elongation.

### Wnt5a regulates sustained digit elongation via convergent extension

Although our ABM successfully reconstructed both the shape symmetry-breaking events, it was unable to recapitulate the digit elongation process beyond spherical distal clusters. Since both Fgf8 and Wnt3 have been shown to promote cell proliferation in a context-dependent manner^19^, we investigated the role of FW in inducing cell proliferation in our organoid system and whether this contributes to symmetry-breaking and digit elongation. We generated limb-mesenchymal organoids and performed EdU-pulse labelling to determine how NC and FW conditions affect distal and proximal cell proliferation, after two hours of chasing on Day 1 and 2. Although FW conditions promoted a higher proliferation rate compared to NC conditions, irrespective of the stage, the ratio of distal to proximal cells remained relatively unchanged (Figure 3A). Furthermore, when cell proliferation was inhibited using Trichostatin A (TSA), tissue elongation was still observed, albeit in smaller aggregates (Figure S6A). These results indicate that an alternative cellular mechanism is required for the sustained elongation of the distal cell clusters.

**Figure 3.**
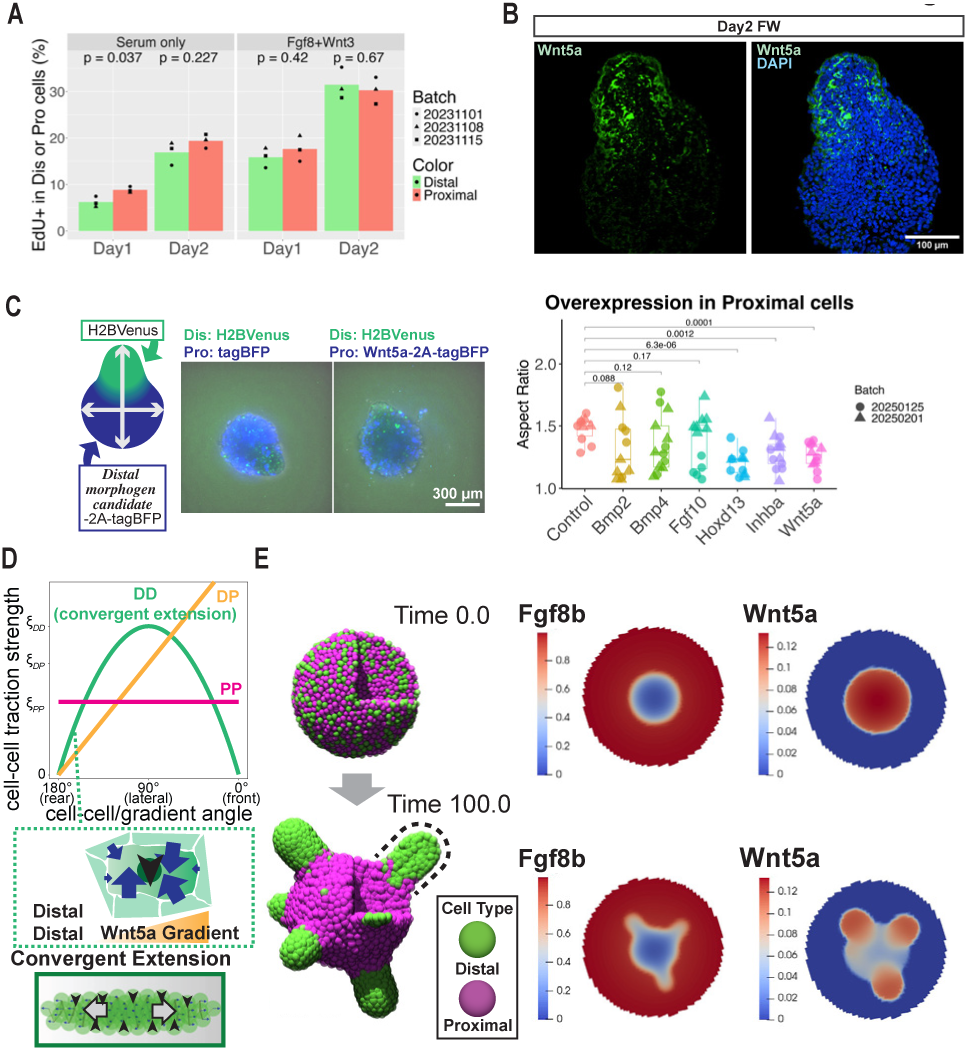
Wnt5a expression in distal cells and convergent extension give rise to digit-tissue elongation. (A) EdU incorporation (% EdU+) in Distal (green) vs Proximal (magenta) cells on Day 1/Day 2 under NC and FW. Bars show means; dots are individual samples (shape = Batch). Two-sided unpaired t-tests (Distal vs Proximal); unadjusted p values shown. (B) Day-2 FW organoid immunostaining of Wnt5a (green) with DAPI (blue). (C) Overexpression screen in Proximal cells. (Left) Schematics and example of the organoids. (Right) Box–jitter plots show organoid aspect ratio by gene (x); each dot is one aggregate (shape = Batch). Control = tagBFP-only; constructs express Bmp2, Bmp4, Fgf10, Hoxd13, Inhba, Wnt5a. Statistics: planned two-sided t-tests vs Control; unadjusted p values are printed above brackets. (D) Rule map: angle to the gradient (x-axis) vs traction strength (y-axis). DD traction is strongest orthogonal to Wnt5a; DP is strongest along Fgf8b; PP is weak/constant. Dotted boxes show traction biases in DD and DP cases. Bottom box shows Tissue-scale CE: Distal motion orthogonal to the Wnt5a gradient drives elongation. (E) ABM combining differential adhesion (DD>DP>PP), DP bias toward Fgf8b, Distal Wnt5a expression, and DD bias orthogonal to Wnt5a. Dotted line marks protrusion shape.

Tissue elongation can occur without exerting force on an external substrate via push-pull mechanisms while maintaining symmetrical cell-cell interactions, by undergoing the process of convergent extension. This can be achieved through the establishment of a morphogen gradient that allows cells to pull each other orthogonally along the gradient or axis of elongation^34^. Previous studies have shown that Wnt5a triggers planar cell polarity, which is known to be necessary for convergent extension in the digit^35–37^. Indeed, we observed that distal cells expressed higher levels of Wnt5a in 2D micromass culture and at the distal tips in 3D limb mesenchymal organoids (Figure S6B, Figure 3B). This suggests that the accumulation of morphogens such as Wnt5a in distal clusters could potentially trigger convergent extension in our organoid system.

As gradient formation is required for convergent extension, to perturb this distal-proximal gradient, we electroporated proximal cells with the distal morphogen gene construct CAG-tagBFP-2A-Wnt5a as well as other potential distal morphogen genes, including Bmp2, Bmp4, Fgf10, and Inhba (encoding Activin). Additionally, since Hoxd13 acts upstream of the distal identities, we used it as a positive control^38,39^. We found that organoids expressing Hoxd13, Inhba, and Wnt5a showed a significant reduction in the aspect ratios (the longest axis vs orthogonal axis; Figure 3C, Movie 7). *In vivo*, the distal-proximal gradient of Wnt5a controls convergent extension and promotes the formation of the expression domain of activin^37,40^. Together, these findings suggest that a distal-proximal gradient (e.g. Wnt5a) is necessary to establish convergent extension for sustained digit elongation.

Based on our data, we added another traction bias in our ABM defined such that two adjacent distal cells pull each other more strongly in the direction orthogonal to the distal-proximal elongation gradient (Wnt5a in this case). In combination with a standard (symmetrical) repulsion force between pairs of cells, this mechanism then allows for a tissue-level convergent-extension motion (Figure 3D). The ABM incorporating differential adhesions, distal expression of Wnt5a, chemotaxis, and convergent extension successfully recapitulated the symmetry-breaking and elongation, showing that these mechanisms are sufficient to explain digit morphogenesis in limb-mesenchymal organoids (Figure 3E see dotted line, Movie 8, Figure S6C: Solid box).

### Quantitative validation and predictive power of the ABM

To validate the contribution of convergent extension for digit elongation in our limb-mesenchymal organoids and the ABM, we developed a novel imaging data analysis pipeline to detect and quantify local tissue deformations. First, we performed two-photon live-imaging microscopy^11^ on the limb-mesenchymal organoids to generate four-dimensional (4D) images and tracked the cell trajectories using TrackMate^41^ and ELEPHANT^42^. Next, we interpolated the velocity vector fields obtained from the tracks using a Gaussian kernel smoothing method. From the smoothed vector fields, we computed a deformation map of the organoid based on a notion originating from continuum mechanics, called Cauchy-Green strain tensor. This tensor measures how a reference sphere in the initial frame of the experiment would be deformed by the vector field. This deformation is represented by ellipses, whereby the longer axes indicate the directions of tissue elongation. We then plotted the primary axes of deformation of these ellipses and projected them onto the elongation axis of the organoid, which essentially measures the extent of convergent extension (Figure 4A-C, see Supplementary Text: Cauchy-Green Tensor Estimation). Our analysis indeed showed that convergent extension is maximal where the Wnt5a gradient is the steepest, at the distal-proximal interface in the limb-mesenchymal organoids (Figure 4D: top panel). Importantly, the same analysis could be applied to the ABM, thereby enabling direct quantitative comparisons between experimental observations and simulations. Consistent with experimental observations, the maximal convergent extension was detected at the distal-proximal cell interface in the ABM implementing differential adhesions, chemotaxis, distal expression of Wnt5a, and convergent extension (Figure 3E and 4D: middle panel). In contrast, when only differential adhesion and Fgf8b-directed traction bias were implemented (Figure 2H), the convergent extension maximum was weaker and co-localized with the distal cluster (Figure 4D: bottom panel).

**Figure 4.**
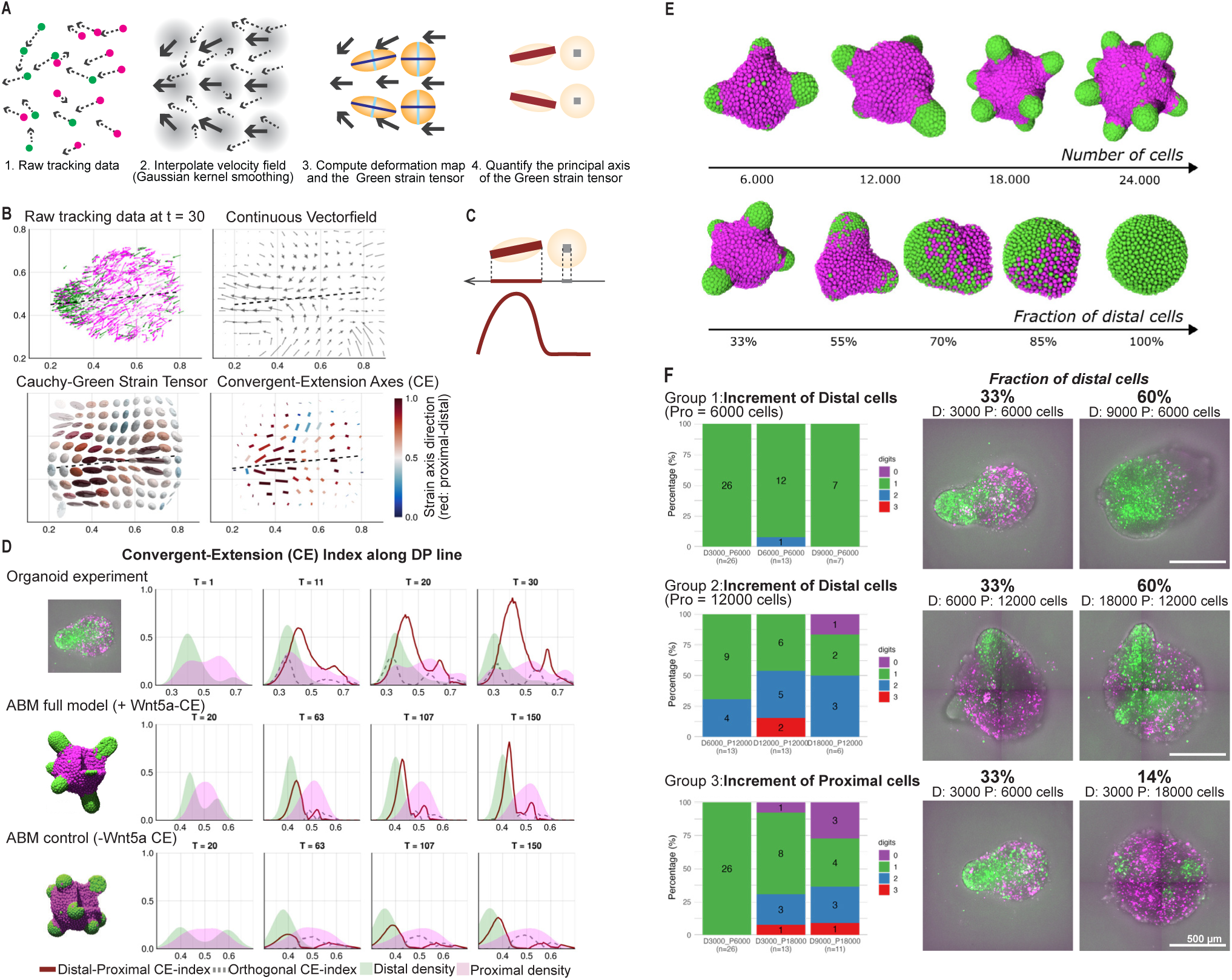
Comparisons of experimental observations and ABM. (A–C) Convergent-extension (CE) analysis. Schematics of workflow (A) and a representative image (B): tracked cell positions are smoothed into a velocity field and integrated to estimate tissue deformation; from this, the primary axis of extension and its strength are extracted and visualized as deformation of a reference spheres into ellipses (See Supplementary document). (C) CE vectors projected along the tissue-elongation axis are plotted versus positions on the tissue-elongation axis (red). (D) CE index projected onto the tissue-elongation axis (solid red curve) and the orthogonal component (gray dashed curve), overlaid with Distal (green) and Proximal (magenta) cell-density profiles, highlighting regions where CE occurs, on experimental data (top), ABM full model, implementing differential adhesion, chemotaxis toward Fgf8b, Distal Wnt5a production, and Wnt5a-driven CE (middle) and ABM without Wnt5a-driven CE implementing differential adhesion, chemotaxis toward Fgf8b only (bottom). (E) ABM parameters controlling protrusions. Predicted protrusion number varies with (top) total cell number (*N*) and (bottom) Distal cell fraction (*D/(D+P)*). The panel summarizes how changes in these parameters shift the distribution of protrusions. (F) Starting composition vs protrusions. Left: Stacked bars show the percentage of organoids with 0, 1, 2, or 3 protrusions for each starting condition (Distal cell count (D)_Proximal cell count(P)). Numbers inside segments = raw counts; x-axis labels include total n; y-axis = %. Right: Representative images at 72 h; each image is annotated with the starting condition and D:P ratio. (top) Group 1 (P fixed at 6,000). (middle) Group 2 (P fixed at 12,000). (bottom) Group 3 (P increased).

The ABM also recapitulated the increase in the number of protrusions as the initial cell number increased in the organoids (Figure 4E, top row). Furthermore, the ABM predicted that as the proximal cell ratio increases, the number of protrusions also increases (Figure 4E, bottom row). Experimental data showed that when the distal cell ratio is higher, distal cell clusters do not bifurcate by themselves (Figure 4F: top and middle panel). In contrast, when proximal cell ratios were higher, it increased the chances of distal cluster separation (Figure 4F: bottom panel), indicating that our ABM is capable of predicting accurately the morphogenetic events occurring in our limb-mesenchymal organoids.

### The minimal sufficient mechanisms for digit morphogenesis in limb-mesenchymal organoids

The final form of our ABM is a dynamical system defined by four constitutive equations describing the evolution of distal and proximal cell positions, and the concentrations of morphogens for convergent extension (Wnt5a, in this case), and for chemotaxis (Fgf8b, in this case). The first equation indicates that the time evolution of a distal cell position can be determined by the sum of two kinds of traction forces and one kind of repulsion forces (Figure 5A, B). The first kind of traction forces is derived from a sum of elementary forces between distal-distal cell pairs within the distance *R*. Each of these elementary forces is directed along the vector linking the two cells with a magnitude given by the strong adhesion (*ξ_DD_*: based on the doublet experiments) and traction bias in the direction orthogonal to Wnt5a gradient (for convergent extension). Likewise, distal-proximal traction forces can be determined by the sum of the vectors linking each distal cell with the proximal cells within the distance *R* of it, weighted by the moderate adhesion (*ξ_DP_*) and with traction bias along Fgf8b gradient (for chemotaxis). The cell-cell repulsive forces come from the requirement of volume exclusion within the distance *δ* (*R* > *δ*) (Figure 5A, B). The time evolution of the proximal cell position can be computed similarly, with the difference that the proximal-proximal traction forces are weighted by weak adhesion (*ξ_PP_*) and with influence from neither Fgf8b nor Wnt5 gradient (Figure 5C). The time evolution of the morphogen concentrations can be defined by a simple reaction-diffusion network. The evolution of Wnt5a concentration is determined by the decay rate and the production rate, which depends on the distal cell density, and the slow diffusion within the organoid (Figure 5D). The evolution of Fgf8b concentration is determined by the decay rate, the saturated diffusion outside the organoid, and the slow diffusion within the organoid (Figure 5E, F).

**Figure 5.**
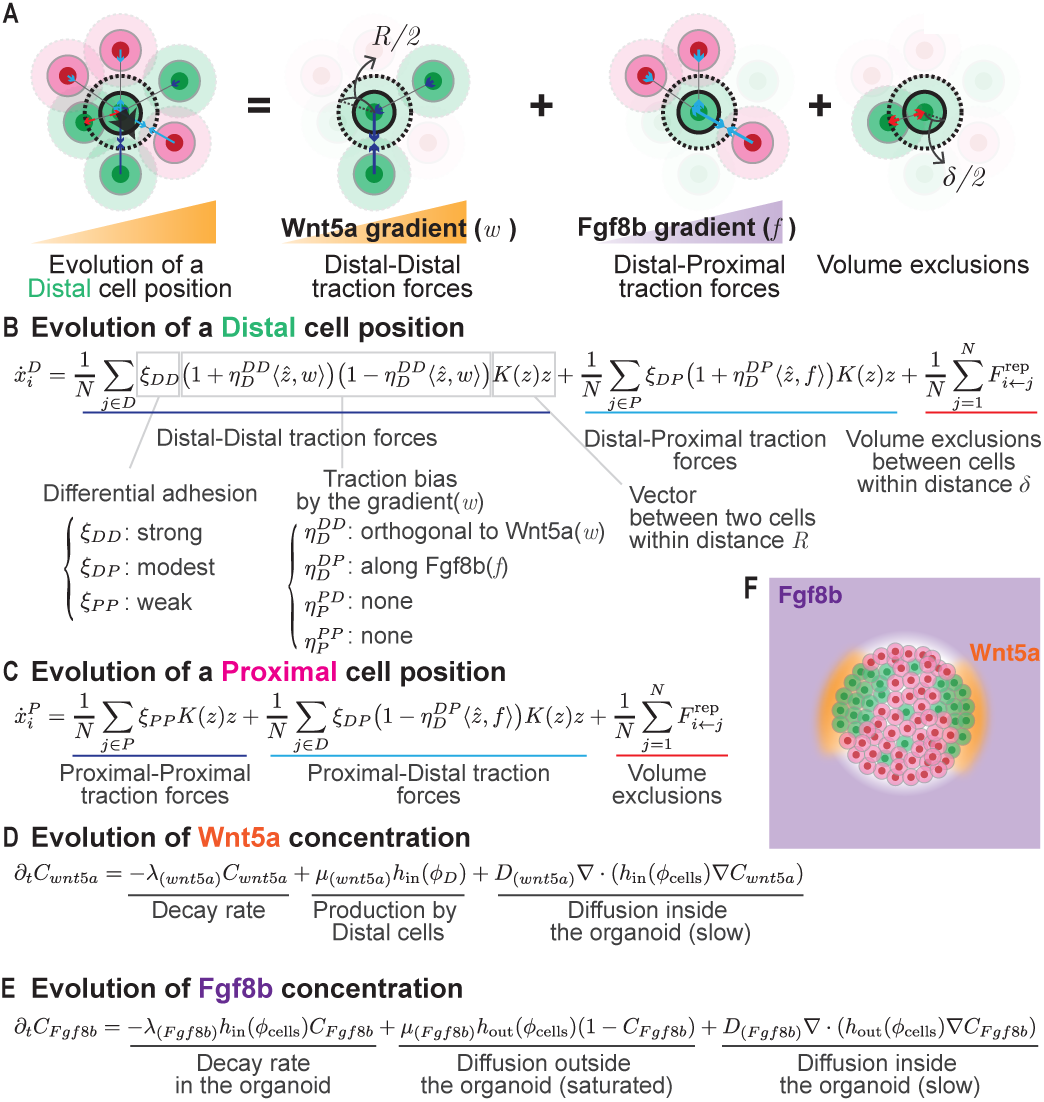
Equations defining the final form of ABM. (A) Schematic for the forces in the ABM: neighbors within diameter *R* exert identity-/gradient-dependent tractions; short-range repulsion prevents overlap; labels link terms to the equations. (B) Evolution of Distal cell position: sum of DD/DP tractions with the above angle biases plus repulsion within δ. (C) Evolution of Proximal cell position: sum of PP/DP tractions plus repulsion within δ. (D) Evolution of Wnt5a concentration: Distal production, slow diffusion, and decay. (E) Evolution of Fgf8b concentration: slow diffusion with decay inside organoids and a saturated external reservoir, forming an inner–outer gradient. (F) Overview of gradients in D–E.

### The derived PDE reveals a mechanochemical fingering instability

While our ABM, with minimal sufficient microscopic mechanisms, shows faithful recapitulation of the tissue-scale outcomes in the limb-mesenchymal organoids after running iterative simulations, it is still difficult to interpret how numerous individual cell behaviors collectively give rise to the emergence of digit morphogenesis. An established method to identify the fundamental mathematical mechanisms underlying macroscopic (tissue)-level emergence originates from statistical physics; it consists in transforming the ABM into a continuum description that is formally written as a system of Partial Differential Equations (PDEs) (Figure 6A). Such continuum equations provide principled mathematical building blocks amenable to analyzing and interpreting the resulting tissue-scale dynamics.

**Figure 6.**
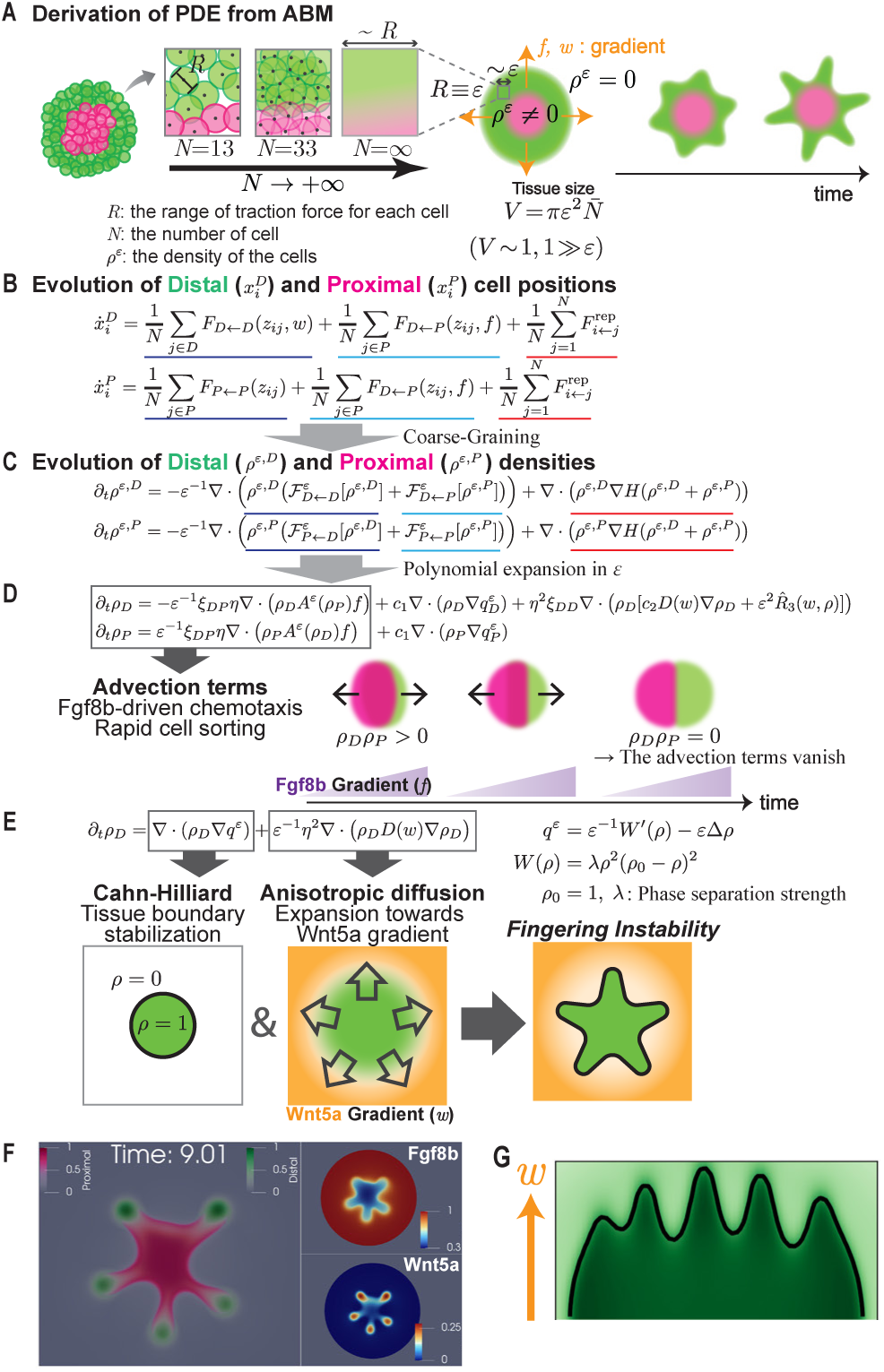
Fingering instability in PDE models. (A) Derivation scheme. The ABM is coarse-grained by letting the cell number *N→∞* while keeping the interaction range *R* constant. At the tissue scale, *R* ≡ *ε* is very small relative to tissue size *V. ρ^ε^*: Distal and Proximal cell densities. *f, w*: chemical gradient (Fgf8b and Wnt5a). (B) ABM: Distal/Proximal positions evolve by pairwise tractions plus short-range repulsion. (C) Distal *ρ^ε,D^* and Proximal *ρ^ε,P^* densities evolution with traction (dark/light blue) and repulsion (red) effects. (D) Advection terms dominate at order *ε^−1^*: Distal density drifts up , opposite to Proximal density (see cartoon). (E) At the next order, a Cahn–Hilliard (CH) equation describes the evolution of the Distal cell density: the CH term is coupled with a novel anisotropic diffusion operator arising from biased tractions in the direction *w*. (F) PDE simulation of limb-mesenchymal organoids: endpoint of the simulation of model in panel D with traction bias *η*=1 and differential adhesion *α*=4 (Left, see Supplementary and Figure S7A), coupled with morphogens evolution following the equations in Figure 5D and 5E (Right). (G) Same as (F) in an *in vivo*-like setup initiated from a hemispherical distal tissue and a constant unidirectional upward (see Figure S7C).

We started from the equations governing the microscopic dynamics of the distal and proximal cell positions (Figure 6B). From there, we derived a set of PDEs for the distal and proximal densities via a mean-field limit^43^ when the number of cells is taken to infinity (*N*→∞). This procedure is followed by an appropriate space-time rescaling that makes the interaction range (*R* ≡ *ε*) much smaller than the size of tissues (*V* = 1) in order to observe the consistent development of shapes at a macroscopic tissue scale. (Figure 6C; see Supplementary Text: Mathematical Theory for details). Eventually, the organoid is described solely by two fields, *ρ_D_* and *ρ_P_*, interpretable as the probabilities (densities) of finding distal and proximal cells at a given location. As the regions of the non-zero densities define the tissue shape, the resulting PDEs describe the evolution of tissue geometry. The force terms and parameters specified in the ABM carry through to the PDE system, while the morphogen concentrations can be described using the same kind of reaction-diffusion network as in the ABM.

The resulting system is composed of three fundamental components (Figure 6D, E). First, an advection term which acts on a fast timescale (the coefficient *ε^−1^*, which is large as *ε* is small). This term corresponds to Fgf8b-driven chemotaxis and produces rapid cell sorting, as distal and proximal cells migrate in opposite directions along the gradient (Figure 6D). Once segregation is achieved, this term vanishes (since *ρ_D_ρ_P_* = 0, corresponding to a zero probability of finding proximal and distal cells at the same location) and the remaining terms become predominant. These terms arise from differential adhesion and Wnt5a-directed traction. The first of these terms results in a Cahn–Hilliard equation^44^ (Figure 6E), which in a similar limit was also observed for other multicellular systems^45–48^. Here, we have extended this approach to two cell populations coupled with a reaction-diffusion network of morphogens. The Cahn–Hilliard term stabilizes the density towards 0 or 1, corresponding respectively to the outside region (medium) and the tissue. If there were only the Cahn–Hilliard term, the tissue would be stable as a sphere (Figure 6E: left panel). However, the traction bias introduces a novel anisotropic diffusion term, whose directionality is set by the Wnt5a gradient. This anisotropic diffusion term alone would uniformly expand the tissue and dilute its own density (Figure 6E: middle panel). Here, the combination of the Cahn–Hilliard term (as a stabilizing factor) and the anisotropic diffusion (as an expanding factor) makes the interface unstable and creates protrusions (Figure 6E: right panel, 6F, S7A, B), a process mathematically linked to the classical “fingering instability” in fluid dynamics (see Discussion)^44,49^.

As in the ABM, the number of protrusions is controlled by the strength of the traction bias (*η*), tissue size (*N*), and the ratio of distal cells (Figure S7B–E). In a setup mimicking limb development in vivo —where morphogenesis initiates from a hemispherical distal tissue with a unidirectional Fgf8b/Wnt5a gradient— digit-like protrusions emerge in the distal direction (Figure 6G, S7C). Thus, although the distal tissue is not a viscous fluid per se, its shape evolution is well described by the same class of PDEs, appearing as finger-like protrusions. Importantly, the PDE is not constructed independently to reproduce finger-like shapes; it was derived directly from, and biologically grounded in, the experimentally validated ABM. Despite its name, fingering instability has not been applied to-date as a mechanistic explanation for the formation of digits or “fingers”. Nevertheless, our work shows that fingering instability is sufficient to explain the emergence of digit-like structures.

## Discussion

To establish an explanatory framework for the emergence of digit tissue-like morphologies, we developed a novel limb mesenchymal organoid capable of forming cell aggregates that spontaneously break morphological symmetry to elongate into digit-like structures. Gene expression profiling and molecular characterization showed that our organoids recapitulate many of the key cellular and molecular features of digit formation during *in vivo* development. Based on our bottom-up approach from an ABM and biological validation, we demonstrate that the tissue-level symmetry-breaking and elongation can be achieved by a set of interacting mechanisms: 1) an inner-outer Fgf8b gradient, 2) differential adhesion, 3) a self-organizing Wnt5a gradient, 4) chemotaxis, and 5) convergent-extension. Coarse-graining the ABM further revealed that these interactions collectively give rise to a fingering instability, which appears to be a key mechanism driving the emergence of digit-like morphogenesis, and drawing an unexpected mathematical link between tissue morphogenesis and a broader class of interface-instability phenomena in fluid dynamics.

The organoid and theoretical model also recapitulated several features of known limb mutant phenotypes. By varying the total cell number and proportion of distal/proximal cells in our model, we were able to mimic phenotypes previously reported in *Hox* and *Gli3* mutants^5^. In *Gli3* mutants, limb buds enlarge and digit number increases, consistent with our observation that larger organoids produce more digit-like protrusions (Figure 1B, C, 4E, top row). While increasing the proportion of distal cells does not enhance the bifurcation of distal clusters, increasing proximal cells yields a higher number of digits in both our organoids and simulations (Figure 4E, bottom row, F). In mutant embryos with complete knock-out of distal *Hox* genes, digits are abolished, resembling the absence of distal cells in our organoids. In contrast, a partial knock-out of distal *Hox* genes increases the number of digits, consistent with our findings that organoids containing fewer distal cells generate more digits (Figure 4E, F). These parallels suggest that our simplified system captures not only generic tissue deformation, but it might also capture developmental relationships between tissue-level phenotypes of digit number and the genetically regulated cellular parameters such as cell number and distal/proximal composition.

The classical Turing model and our model provide explanations of two complementary aspects of digit development rather than opposing each other: the patterning and morphogenesis. During normal limb development *in vivo*, early digit patterning has been explained by Turing-type reaction–diffusion mechanisms operating across the anterior–posterior axis, with periodic molecular patterns appearing before overt morphological changes^5,6^. In this context, distal cells are typically not in direct contact with proximal cells, and previous 2D cultures of distal cells appear to capture this early phase of molecular pattern symmetry-breaking^5,6,30^ (Figure S4B, C). In contrast, our experimental approach —mixing distal and proximal cells in 2D/3D— allows us to capture morphological symmetry-breaking and subsequent digit elongation along the distal-proximal axis. Importantly, many of the features of our model are already supported by in vivo limb developmental studies. Although distal-proximal cell fate of the limb bud is specified by the biochemical gradients, including Fgf8 and Wnt3^19,20^, cell sorting also likely contributes to the establishment or maintenance of proximal-distal domains, during limb development *in vivo*, as observed in our model^32,50^. Likewise, Wnt5a-dependent convergent extension of the distal clusters in our organoids is also well aligned with known mechanisms of digit elongation *in vivo*^35–37^. Thus, the contribution of our study is not to identify these molecular and cellular mechanisms de novo, but to distill mechanisms underlying distal autopod tissue establishment and digit morphogenesis along the distal-proximal axis —mechanisms already implicated *in vivo*— into organoid and ABM frameworks (Figure S8). This reconstitution shows that these cell-level rules are sufficient to recapitulate the onset of digit-like tissue morphogenesis, and coarse-graining the experimentally grounded ABM supports the interpretation that their collective action gives rise to a tissue-scale mechanochemical fingering instability.

Depending on the context (e.g. stage and species), one mechanism may dominate over the other. In the 2D culture mixing distal and proximal cells, the progressive accumulation of distal cells coincided with cartilage differentiation in the 2D micromass culture (Figure S3D, E, S4A), although cell-cell biochemical interactions were not extensively analyzed. Together with the distal-only cultures, these observations suggest that patterning and morphogenesis are highly coordinated, likely utilizing a combination of both mechanical and biochemical interactions. In mouse embryos, the earliest stage of digit patterning involves a periodic arrangement of molecular signals, which is seen hours before any morphological changes are found^6^, suggesting that biochemical reaction–diffusion has a leading role at early stages. Amphibians, by contrast, form their morphological digits sequentially, a process where mechanical interactions may be especially prominent^51^. Our results suggest that these cases need not represent mutually exclusive mechanisms. Instead, species-specific developmental systems may occupy different regions of a shared mechanochemical landscape, in which biochemical patterning and tissue-level fingering contribute to different degrees.

The continuum limit of our ABM places digit-like morphogenesis within the broader mathematical class of fingering instabilities. Fingering instability is often discussed in the context of the Saffman–Taylor problem, where one viscous fluid invades into a less viscous one^44,49^. In general, this is a question of balance between stabilizing and expansive factors: the fluid interface tends to be round or flat due to the surface tension (stability), but when coupled to an expansive drive, the interface may develop protrusions (fingering). While we did not set out to equate digit morphogenesis with viscous-fluid fingering, the same mathematical framework arose unexpectedly from the continuum limit of our ABM. In this setting, the Cahn-Hilliard-based stabilization arises from the isotropic part of cell-cell adhesion and repulsion, whereas the expansive drive arises from anisotropic traction bias (in this case, driven by Wnt5a). The same general balance between stabilizing and destabilizing terms can therefore generate fingering morphology without requiring the tissue to behave as a classical fluid. Therefore, because convergent extension enters our continuum model as an effective anisotropic traction term, this framework is not restricted to digit-like morphogenesis. Rather, it should be, in principle, extendable to developmental contexts in which tissue-scale deformation is driven primarily by convergent extension or related polarized cell behaviors, including elongations of neural tube, notochord, renal tubule, mandibular arch, and cochlear duct, as well as gastruloid axial elongation^52–57^.

The parameters in our model should be interpreted as coarse-grained biological quantities rather than literal fluid-mechanical variables. Differential adhesion and traction bias may arise from various biochemical and physical origins, including anisotropic cell–cell or cell–matrix interactions, polarized adhesion molecules, spatial heterogeneity in extracellular matrix (ECM) distribution or stiffness and biased filopodial dynamics^33–37,48,58–60^. Especially, the involvement of ECM is suggested by related Cahn–Hilliard-type active-solid theory linking cell contractility and ECM elasticity to cartilage-condensation-like patterning^48^, as well as experimental evidence implicating reciprocal cell–ECM/supracellular interactions in limb cartilage patterning^59^. Future work will be needed to determine how these molecular and physical sources map onto the effective parameters of our model in vivo.

A key distinction of our framework from the previous works applying fingering instability in biological contexts is that the continuum description is derived from cell-level rules rather than assumed at the outset. Cahn-Hilliard equations have been used previously to explain biological phenomena such as tumour growth and expansion of epithelial colonies^61,62^, but these approaches generally begin from a fluid-like continuum tissue description. In our work, by contrast, the tissue-scale instability emerges from an experimentally validated ABM of limb mesenchymal cell behaviours. Although our derivation and simulations identified fingering instability as a general outcome of the derived PDEs, and revealed a parallel with classical viscous-fluid fingering at a fundamental mathematical level, the PDE introduced here has a distinct mathematical structure from those classical models. Thus, its mathematical framework introduced here suggests further analytical investigations to give a rigorous interpretation of the interplay between reaction-diffusion systems, geometry and mechanical constraints that leads to instability phenomena. Such a rigorous mathematical framework for the PDE model that we derived has recently been initiated in Fritz 2026a, b^63,64^.

More broadly, our study contributes to a growing effort to use multicellular in vitro systems to uncover principles of emergent tissue morphogenesis. In epithelial tissues, such as the optic cup and the intestinal crypt, organoid models have shown how tissue mechanics, geometry, and cell fate can interact to drive self-organized morphogenesis^11,65–68^. These insights have enabled robust and precise control in tissue engineering^69,70^. With mesenchymal tissues, “gastruloids” that model the development of growing axial tissues, show similar morphological characteristics as our model, including symmetry-breaking and elongation, that are based on differential adhesions and convergent extension^55,71–73^. Together with ours, these studies provide insights into the dynamic relationship between the genotypic variation controlling microscopic cellular behaviours and the macroscopic tissue-level morphological phenotypes.

Lastly, because the parameters of our PDE are directly connected to the underlying ABM, this framework may provide a route toward empirically estimating the developmental parameters that shape digit number and morphology. This may also reveal underlying mechanisms governing evolutionary conservation or variation of the digit number. For instance, despite the size of limb buds varying across animal species and number of digits being determined by the size of the limb bud, the five digit phenotype is robustly conserved among most extant tetrapods^74^. While some animals exhibit digit reduction via *ad hoc* developmental mechanisms^75,76^, virtually no extant species has more than five digits. Our model suggests that such robustness, often referred to as developmental constraints, may reflect not a single hard-wired genetic program, but a stable region of parameter space in which multiple mechanisms jointly regulate digit formation. Species differences may shift the balance among growth, cell composition, adhesion, ECM interactions, Wnt/Fgf signaling, and traction, thereby changing the prominence of each mechanism without requiring an entirely different theory of digit development. Overall, our work offers a conceptual and experimentally tractable framework for linking molecular and cellular variation to tissue-scale morphology, with implications for developmental biology, evolutionary biology, and precision tissue engineering.

## Method details

### Animals

All animal procedures comply with all relevant ethical regulations and guidelines for animal studies, were approved by Research Ethical Committee Kyoto University and licensed by the Japan Ministry of Education, Culture, Sports, Science and Technology and the Science Council of Japan. Mice were obtained from Japan SLC, Inc.

### Cell culture

Limb mesenchyme was isolated from ICR or B6Jx129svEX/ICR Sox9-KI-GFP mouse strains, following a protocol modified from previous reports^18,19^. Briefly, the anterior two-thirds of distal forelimb buds (E11.5) were dissected in 10% fetal bovine serum (FBS, 173012, Sigma-Aldrich) / DMEM (D5796, Sigma-Aldrich) / Penicillin-Streptomycin (15140-122, Gibco) (NC medium) and collected into a 15-ml tube. Samples were rinsed with 5 ml PBS, then incubated in 1% trypsin/PBS at 4 °C for 30 min. The reaction was quenched with 2 ml of NC medium, and buds were transferred to 6-cm Petri dishes containing 3 ml of the same medium. Under a stereomicroscope, the ectoderm was carefully removed, and the distal and proximal halves of limb mesenchymal tissue were cut into separate tubes. The tissues were dissociated by gentle pipetting in culture medium, and cell numbers were determined using a hemocytometer.

Electroporation was performed using a modified protocol of a previously described method^77^ using the SF Cell Line 4D-Nucleofector™ X Kit S (V4XC-2032, Lonza) and the 4D-Nucleofector™ Core Unit (AAF-1003B, Lonza) with X Unit (AAF-1003X, Lonza). Cell pellets were resuspended in 20 µl of SF solution plus Supplemental 1 solution. For labeling, suspensions were mixed with 62.4 ng/0.5 µl of plasmids encoding CMV-H2B-Venus, CAG-H2B-tdTomato (Plasmid #116870, Addgene), CAG-H2B-tagBFP, or 7Tcf-EGFP. For overexpression experiments, 1 µg/0.5 µl of plasmid DNA encoding CAG-MmBmp2-, MmBmp4-, MmHoxd13-, MmFgf10-, MmWnt5a-, or MmInhba-P2A-tagBFP was added. Electroporation was carried out using program DS113, after which 80 µl of NC medium was added to each suspension. Cells were allowed to recover at 37 °C for 10 min. Following recovery, cell counts were adjusted to conserve the original Distal: Proximal ratio (approximately 1:2) in culture.

The cells were pelleted at 230 g for 10 min, then resuspended at 2.0 × 10⁵ cells/10 µl for 2D culture (plated on 3.5-cm dishes) or at 9.0 × 10³ cells/100 µl for 3D culture in ultra-low adhesive 96-well plates (PrimeSurface® Plate 96U, MS-9096U, Sumitomo Bakelite) in NC medium or 10% FBS/DMEM/PS supplemented with 150 ng/ml recombinant human/murine FGF-8b (100-25-250 µg, Peprotech; stock 25 µg/ml) and 250 ng/ml recombinant murine Wnt-3a (315-20-10 µg, Peprotech; stock 40 µg/ml) (FW medium). For 3D conditions, after 24 h, 20 µl of 18% Geltrex™ (A1413302, Gibco) in NC or FW medium was added to each well (final concentration 3%).

### Live Imaging and Image Analysis

2D micromass culture Live imaging data were acquired using either an incubator confocal microscope CellVoyager CV1000 (Yokogawa), or an incubator two-photon microscope LCV-FVMPE-RS (Evident)^11^.

For quantification of Sox9 intensities in individual cells, counting the number of distal and proximal cells in k-neighbors, and calculating the nearest neighbor segregation (NNS; Figure S3C), images obtained with the CellVoyager were processed for creating a hyperstack and cell segmentation using ImageJ/FIJI and its plugin TrackMate ^41^. The ImageJ macros and Python codes used for this analysis are available on GitHub ^78^. Based on the spatial coordinates of cells in each channel, NNS was computed as follows

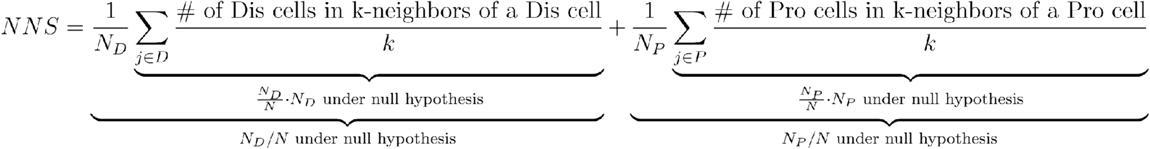

Of note, NNS was normalized to equal 1 under the null hypothesis (random distribution), independent of the overall distal: proximal ratio. NNS values greater than 1 indicate clustering of the same cell type.

For digit organoid cell tracking, images acquired with the LCV-FVMPE-RS microscope were segmented by the LoG detector with a diameter setting of 8 µm. Subsequently, ∼400 cells per image were manually tracked using TrackMate. These manually curated trajectories were then used as a training dataset for automated tracking with the deep learning–based 3D cell tracking systems Mastodon ^79^ and Elephant ^42^.

### PIC experiment

Five organoids, originating from four surrogates cultured in FW conditions, were snap-frozen in Tissue-Tek O.C.T. Compound (Sakura Finetek Japan) at Day 2. Cryosections prepared at a thickness of 10 µm were washed with PBS and fixed with 4% PFA/PBS for 10 min at room temperature. Specimens were permeabilized with 5% TritonX-100 in PBS-DEPC for 3 min and then with 0.1 N HCl for 5 min, followed by neutralization with 1 M Tris-HCl, pH 8.0. Permeabilized specimens were incubated in PBS-DEPC for 5 min at 65 °C and quickly cooled in ice-cold PBS-DEPC. In situ reverse transcription mix [0.5 ml of 500 ng/ml caged primer, 0.5 ml of 10 mM dNTPs (NEB), 5 ml of H_2_O, 2 ml of 5 × First-Strand Buffer, 1 ml of 0.1 M DTT, 0.5 ml of 40 U/ml RNase Out, and 0.5 ml of 200 U/ml Superscript II Reverse Transcriptase, all from Invitrogen] was applied to the specimens, which were then incubated at 42 °C for 60 min, followed by heat-inactivation in 70 °C PBS for 10 min.

Photo-irradiation for uncaging was performed with a multi-pattern LED illumination system (Opto-line; LEOPARD2) and an LED light source (Prizmatix; UHP-F-365 LED; 3 W). The light was illuminated to specimens through an objective lens [HC PL FLUOTAR 20x/0,55] at a wavelength of 365 nm for 3 min. Masking sheets loaded on the system were made by ImageJ/Fiji.

After photo-irradiation, specimens were lysed and cDNA:mRNA hybrids were purified with a Qiagen MinElute PCR Purification kit. The sequencing library was prepared according to a previous report^21^. The library was sequenced on the Illumina Novaseq 6000 platform. The sequence data described in this study have been deposited in Gene Expression Omnibus (GEO) under accession number GSE331258.

### scRNA-seq sample preparation

Micromass aggregates were enzymatically dissociated and barcoded prior to single-cell capture. Reagents were prepared as follows: Pronase (537088-10KUCN, Millipore) was reconstituted to 7,800 U/ml in PBS (1/100 stock; 10 µl aliquots). A 780 U/ml working solution was made immediately before use by mixing 900 µl DMEM with 100 µl pronase stock solution to a final volume of 1 ml, then filtered to remove debris.

A Day 0 samples (Distal/Proximal) were washed four times in NC medium with 1,000 rpm, 5 min spins. Organoids at Day 1 and Day 2 (FW/NC) were collected with a wide-bore pipette (24 organoids for each condition on Day1 and Day2), washed once in 1 ml PBS, and digested in 300 µl pronase working solution at 37 °C for 45 min, with vortexing every 15 min. Digestion was quenched with 500 µl NC medium, followed by centrifugation (1,000 rpm, 5 min).

Pellets were resuspended in 100 µl NC medium. After cell count, 100,000 cells each were taken, centrifuged (1,000 rpm, 5 min, 4°C), then resuspended in 100 µl of 15 µM DNA–lipid solution (×2 tubes) and incubated 30 min at 37 °C in CO₂. Cells were then diluted with 900 µl 1% BSA/PBS (total 1 ml) and centrifuged (1,000 rpm, 5 min, 4 °C). After resuspension in 1% BSA/PBS (1 ml), samples with the same barcode label were combined, centrifuged again (1,000 rpm, 5 min, 4 °C), resuspended in 1% BSA/PBS (60 µl), and counted. A final spin (1,000 rpm, 5 min, 4 °C) preceded adjustment to 700 cells/µl. DNA–lipid Barcode-1 and Barcode-2 labels were used. For sample loading, the cells were targeted at 5,000 cells per sample (10,000 cells total) for each of Days 0, 1, and 2.

### scRNA-seq analysis

Single-cell 3′ gene-expression libraries were prepared with the 10x Genomics Chromium platform according to the manufacturer’s instructions with modifications (Chromium Next GEM Single Cell 3’ Kit v3.1, 4 rxns, PN-1000269, User Guide CG000389, 10x Genomics). We used in-house lipid-DNA barcoding for cell multiplexing, as described above. CMO-based demultiplexing was performed in two steps. First, paired gene-expression and multiplexing-capture libraries for each day were processed with *cellranger multi* using a custom mouse reference (mm10-2020-A) extended with sequences for the H2B-Venus and H2B-tdTomato transgenes and a CMO reference to obtain initial sample labels. The initial per-day runs produced feature-barcoding and gene-expression count matrices. Sample identifiers were Day0_Distal and Day0_Proximal (Day 0), Day1_NC and Day1_FW (Day 1), and Day2_NC and Day2_FW (Day 2). We then imported CMO counts into Seurat and classified barcodes as singlet/doublet/negative. Final per-cell assignments were exported and supplied back to *cellranger multi* to regenerate per-sample count matrices and summaries, which were used for downstream integration and analysis.

Per-sample outputs were merged into a Seurat object (constructed with ≥200 genes per cell and genes present in ≥3 cells). For higher stringency, we restricted analyses to nFeature_RNA ≥ 2,500, then recomputed the embedding and graph on PCs 1–20 (UMAP from PCA; neighbors on the SCT assay) and performed Louvain clustering (algorithm = 1, resolution = 0.8). CMO assignments were imported from Seurat demultiplexing tables and standardized to six classes — Day0_Distal/Day0_Proximal, Day1_NC/Day1_FW, Day2_NC/Day2_FW— using day-prefixed barcodes; plots restricted to these labels where indicated.

Denoising was then applied using RECODE (Python screcode via reticulate)^80^. Before RECODE we computed QC metrics and retained cells with percent.mt ≤ 30% and hemoglobin ≤ 64 UMIs (summed over *Hba-a2, Hba-a1, Hbb-b2, Hba-x, Hbb-b1*), and excluded mitochondrial (mt-), ribosomal (Rps/Rpl), and hemoglobin genes from the feature set. We normalized/scaled this assay, ran PCA, and computed UMAP on PCs 1–30.

Analyses and figures for the FW-only subset were generated from the extracted Seurat object with the class Day1_FW and Day2_FW after the RECODE denoising. To evaluate the presence of epidermal/ectodermal vs. mesenchymal signatures, we generated DotPlots and summary tables using curated panels (e.g., Krt, Epcam, Cdh1 for ecto/AER; mesenchymal references) and reported (i) per-gene detection as the % of cells with ≥1 UMI and (ii) a per-tissue summary defined as the % of cells expressing ≥2 panel genes.

To quantify spatial programs, we imported spatial differential-expression results obtained from PIC and retained genes with padj < 0.05. We defined Distal and Proximal gene sets as the top 20 protein-coding genes by effect size (Distal: log2FC > 1; Proximal: log2FC < −1), and computed per-cell module scores (Seurat AddModuleScore) yielding Distal_Score and Proximal_Score. UMAP feature plots displayed the module scores with display ranges tuned per score (e.g., Distal: 1 to 3; Proximal: -3.5 to -0.5).

Velocity analyses were performed with scVelo on the RECODE-denoised FW dataset ^81^. Spliced/unspliced layers were supplied by concatenating loom files for Day1_FW and Day2_FW. The existing UMAP (from the RECODE assay) and Seurat clusters were used for visualization and grouping. Preprocessing followed scVelo defaults (pp.filter_and_normalize, then pp.moments), velocities were computed with *tl.velocity* (mode = “*stochastic*”). Velocity fields were plotted on the UMAP (stream).

Lineage inference was performed on the same UMAP using Slingshot ^82^; the root cluster was set to Day1_FW_Distal_1, and two endpoints were specified a priori: Day2_FW_Distal_2 (distal) and Day2_FW_Proximal (proximal), informed by the scVelo results. Gene expression dynamics along Slingshot pseudotime were modeled with *tradeSeq* (negative-binomial GAMs with spline smoothers), and *associationTest* was used to assess whether expression varied with pseudotime; genes with FDR < 0.05 were considered dynamic; selected developmental regulators (e.g., *Sox9, Wnt5a, Fgf10, Inhba, Bmp2, Bmp4*) were plotted as smoothed trends along inferred lineages.

For computing FW and NC signatures, Cells were grouped into six bins using CMO assignments with day labels: Day1-FW-Distal, Day2-FW-Distal, Day1-FW-Proximal, Day2-FW-Proximal, Day1-NC, Day2-NC. Signature derivation (Day 1): Differential expression (Wilcoxon) was run for Day1-FW (Proximal + Distal) vs Day1-NC with min.pct = 0.1 and logfc.threshold = 0. From differential expression results, we defined two 50-gene signatures at FDR < 0.05: a FW-up set (top 50 by descending avg_log2FC) and an NC-up set (top 50 by ascending avg_log2FC). Per-cell module scores were computed with *AddModuleScore* for the FW-up gene set (FW signature score) and the NC-up gene set (NC signature score). UMAP feature maps were drawn with percentile cutoffs: FW displayed from the 10th–90th percentiles of FW signature score; NC from the 20th–60th percentiles of NC signature score. Plots and display ranges: Violin/box plots showed FW signature scores across the six bins.

We analyzed an external mouse limb metacell atlas kindly provided by Elazer Zelzer’s laboratory at Weizzman Institute^26^. To compare our organoid signatures against atlas cell states, we computed FW and NC module scores at the metacell level. Scores were plotted on the atlas 2D map and in a bivariate scatter (FW score vs NC score), colored by atlas group labels inherited from Markman et al. 2023^26^. Developmental time was assigned to metacells by majority vote of their constituent single cells, and metacells were colored by this discrete time for visualization on the atlas layout.

To compare FW organoid cell states with the in vivo limb metacell atlas more directly, we constructed FW organoid metacells from the RECODE-denoised FW object. The FW object was reclustered on PCs 1–20 at high resolution (resolution = 5.0), and clusters containing fewer than 20 cells were removed before metacell-level analysis. For each FW metacell, the mean RECODE-normalized expression profile was calculated, and the original FW cluster label was assigned by majority vote. Markman metacell labels were recovered from the atlas color annotation defined in Markman et al. 2023^26^. Shared genes between the FW and Markman metacell matrices were z-scored within each dataset and compared by Pearson correlation. To visualize atlas cell states in the organoid embedding, each Markman metacell was projected onto the FW organoid UMAP as a weighted average of the coordinates of its top five correlated FW metacells, with negative correlations set to zero before weight normalization. The same correlation matrix was then used to generate the FW-versus-Markman metacell correlation heatmap.

### Hybridization-based in situ sequencing (HybISS)

HybISS was performed largely as described previously^27^, with minor modifications. Padlock probes were phosphorylated with T4 polynucleotide kinase (M0201S, NEB) in the presence of ATP (P0756S, NEB) at 37°C for 30 min, followed by heat inactivation at 65°C for 20 min. Organoids cultured in FW conditions were snap-frozen in Tissue-Tek O.C.T. Compound (Sakura Finetek Japan) at Day 2. Cryosections prepared at a thickness of 10 µm were washed in PBS, post-fixed with 3% formaldehyde in PBS for 5 min at room temperature, treated with 0.1 M HCl for 5 min, dehydrated sequentially in 70% and 100% ethanol, and air-dried. Secure-Seal hybridization chambers (621505, Grace bio-labs) were then attached to the sections.

For in situ reverse transcription, sections were incubated with random decamers at 65°C for 5 min and placed on ice for 5 min. Reverse transcription was carried out using SuperScript IV (18090010, Invitrogen) reverse transcriptase in 1× SSIV buffer containing DTT, dNTPs (18427013, Invitrogen), BSA (B9001S, NEB), random decamers (AM5722G, Invitrogen), and RNaseOUT (10777019, Invitrogen). Reactions were incubated for 10 min at room temperature and then overnight at 42°C in a humidified chamber. Sections were subsequently post-fixed in 3% formaldehyde in PBS for 40 min at room temperature.

Padlock probe hybridization and ligation were performed in ligation buffer containing KCl, formamide (064-0042, Fujifilm), BSA, phosphorylated padlock probes, Tth DNA ligase (EN13-250, QIAGEN), and RNase H (M0297S, NEB). Sections were incubated at 37°C for 30 min followed by 45°C for 90 min. Rolling-circle amplification was performed overnight at 30°C using Phi29 DNA polymerase (4001, Montserate) in Phi29 buffer containing glycerol, dNTPs, BSA, and Exonuclease I (M0293S, NEB).

For hybridization-based decoding, rolling-circle products were first hybridized with bridge probes in 2× hybridization buffer (4x SSC (UltraPure™ SSC, 20X, 15557-044, Invitrogen), 40% formamide) for 1 h at room temperature with gentle agitation. Sections were rinsed in 2× SSC and incubated with fluorescent detection oligonucleotides and DAPI in 2× hybridization buffer for 1 h at room temperature, protected from light. After washing in PBS, sections were mounted with SlowFade Gold Antifade Mounting Medium (S36937, Thermofisher) and imaged. For subsequent decoding cycles, coverslips were removed in PBS, and fluorescent probes were stripped using 65% formamide in 2× SSC at 30°C for 30 min. Sections were then washed in 2× SSC before the next hybridization cycle. Probe sequences are shown in Table S1.

HybISS images were processed in ImageJ/Fiji for visualization. Z-stack images from sequential hybridization cycles were aligned using DAPI signals as references using Register Virtual Stack Slices plugin^83^. HybISS spots were identified by particle detection in ImageJ/Fiji with a size threshold of 1 pixel to infinity and circularity of 0.00–1.00. For display in Figure 1K, the detected spot centroids were rendered as filled circles in the corresponding gene channels, while the DAPI channel was shown as the original projected image. The unrendered original HybISS images are shown in Figure S1I.

### Green strain tensor

We outline below the estimation of Green strain tensors for in vitro and in silico digit organoids (see Supplementary Text: Cauchy-Green Tensor Estimation for a detailed exposition). From the tracking data (in vitro) or full cell trajectories (in silico), we approximated a space–time continuous velocity field *v*(*x*, *t*). First, we scaled the datasets to fit into a unit cube. Then we performed Gaussian kernel smoothening in space using the formulas

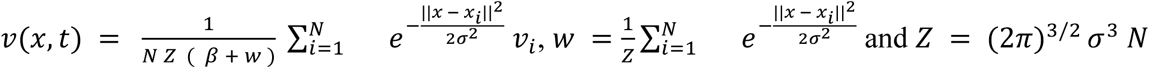

where *t* is the current frame, *N* is the number of tracks per frame, *x_i_*, *v_i_* are the spot locations and velocities and *σ* = 0.138, *β* = 0.5 are the smoothening width and bias parameter. Furthermore, we applied linear interpolation in time, yielding a space-time continuous field *v*(*x*, *t*). We then estimated the deformation map *φ*(*X*, *t*) of the organoid between a reference time *t*_0_ and a final time *t*_1_ by solving the differential equation *ẋ* = *v*(*x*, *t*) with initial condition *x*(*t*_0_) = *X*, which yielded the deformation as *φ*(*X*, *t*) = *x* = *x*(*t*_1_; *x*_0_ = *X*). The deformation gradient was then 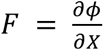 and to remove effect of rotational and translational motion, we computed the Green strain tensor as *C* = *F F^T^*. With the eigenvalues *λ*_1_ ≤ *λ*_2_ ≤ *λ*_3_of the Green strain tensor, we could represent the deformation as ellipses with corresponding axis lengths and we defined the convergent-extension (CE) axes by the eigenvector of the largest eigenvalue ( *V*_3_) scaled by a measure of elongation, namely 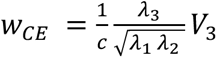 where c is a global normalization constant to constraint the values between 0 to 1. We finally projected these CE-axes *w_CE_* onto the anterior-posterior axis *w_CE_* ⋅ *n_AP_* to obtain a scalar measure for convergent extension.

### Doublet angle measurement

Cell aggregates composed of either distal or proximal cells were cultured under NC or FW conditions and collected into 1.5-ml tubes at Day 1 and Day 2. Aggregates were washed with 1 ml PBS to remove Geltrex, then resuspended in 50 µl of MemGlow-FW medium, prepared by supplementing NC or FW medium with 20 µM MemGlow 488 (MG01-02, Cytoskeleton, Inc.) or MemGlow 560 (MG02-02, Cytoskeleton, Inc.). Aggregates were combined in the following groups: Distal-488 (16 aggregates), Distal-560 (8 aggregates), Proximal-488 (8 aggregates), and Proximal-560 (16 aggregates). Cells were dissociated by vigorous pipetting (100 times) and incubated at room temperature for 10 min. One milliliter of NC medium was then added, and cells were centrifuged at 230 g for 5 min. The supernatant was discarded, replaced with 1 ml of fresh NC medium, and centrifuged again at 230 g for 5 min.

The resulting pellet was resuspended in NC or FW medium and mixed to generate the following combinations: Distal-488 (50 µl) + Distal-560 (50 µl), Distal-488 (50 µl) + Proximal-560 (50 µl), and Proximal-488 (50 µl) + Proximal-560 (50 µl) (Figure S5A). Suspensions were allowed to adhere for 15 min at room temperature and then centrifuged at 230 g for 5 min. After centrifugation, medium and bubbles were carefully removed, leaving 20 µl.

For imaging, 10 µl of the cell suspension was placed into a custom slide glass window. Sterilized microscope slides and 24 × 26 mm cover glasses were autoclaved, and a 5-mm square window was created on the slide glass using layered vinyl tape. Both the slide window and cover glasses were coated with 4 µl of 2-methacryloyloxyethyl phosphorylcholine solution. The cover glass was placed over the suspension, and the edges were sealed with nail polish.

Images were captured immediately using an Inverted Microscope DMi8 (Leica) equipped with an ORCA®-Flash4.0 LT3 Digital CMOS camera (C11440-42U40, Hamamatsu Photonics). From these images, doublets exhibiting both MemGlow 488 and 560 signals were selected, as these were presumed to be formed after dissociation. The angles between the two cells from two independent experiments were measured in ImageJ by technical staff blinded to the expected outcomes.

### Immunostaining

Organoids were fixed with 4%PFA/PBS for 2 hours, immersed in 20% Sucrose/PBS overnight, and embedded in Tissue-Tek O.C.T. Compound (Sakura Finetek Japan). Cryosections with 10 µm thickness were air-dried on a 37 °C plate for 15 min, then rinsed in PBS. For staining for anti-phosphorylated-ERK, TBS was used instead of PBS throughout. For staining for anti-Wnt5a, the cryosections were dehydrated in a graded ethanol (70%, 80%, 90% and immersed in 100% overnight). Subsequently, sections were hydrated in 90%, 80%, 70% Ethanol, followed by a brief wash, antigen retrieval at 95℃ for 20 min with Antigen Unmasking Solution, Citrate-Based (H3300-250, Vector Laboratories), and wash in PBS for 3 min.

The Sections were blocked in Blocking One (03953-95, Nacalai Tesque; also used as antibody diluent). Primary antibodies were applied at the following dilutions: rabbit anti-phosphorylated-ERK1/2 (4370S, Cell Signaling, 1:500), rat IgG2a anti-GFP (04404-84: Nacalai Tesque, Clone: GF090R, 1:1000), and mouse anti-Wnt5a (made in this study, 1:100) incubated overnight at 4 °C in a humidified chamber. Slides were then washed 3 × 5 min in 0.05% Tween-20 / PBS at RT. Secondary antibodies were prepared in Blocking One at the following dilutions: donkey Cy3 anti-rabbit IgG(H+L) (711-165-152, Jackson lab, 1:500), donkey 488 anti-rat IgG(H+L) 712-545-153, Jackson lab, 1:500), and donkey 488 anti-mouse IgG(H+L) (A21202, Thermo Fisher Scientific, 1:500) and incubated 1 h at RT in a humidified chamber, followed by 3 × 5 min washes in 0.05% Tween-20/PBS and a final PBS rinse. Slides were then coverslipped using SlowFade Gold Antifade Mountant (S36936, Invitrogen).

### Anti-Wnt5a antibody production

Balb/c mice were immunized with purified mouse Wnt5a protein (6His-tagged Wnt5a [residues 40-380]) expressed in E. coli BL21(DE3), using Freund’s complete adjuvant. Lymphocytes from the spleens of one of these immunized mice were fused with P3-X63-Ag8-U1 myeloma cells, and hybridomas producing anti-Wnt5a antibodies were screened. Screening was performed using ELISA, Western blot analysis, and immunohistochemistry on mouse tissue sections. As a result, three independent anti-Wnt5a antibody-producing hybridomas were obtained. In this study, the IgG1 antibody purified from the culture supernatant of one of these hybridomas (clone 5-5D) was used.

### Aggregate pairing experiment

Distal cells labeled with H2B-Venus and Proximal cells labeled with H2B-tdTomato were mixed at their original Distal:Proximal ratio, and 9.0 × 10³ cells per well were plated in ultra-low adhesive 96-well plates (PrimeSurface® Plate 96U, MS-9096U, Sumitomo Bakelite) in NC medium to form mixed aggregates. In parallel, Proximal cells expressing tagBFP, or Fgf8b-P2A-tagBFP, Wnt5a-P2A-tagBFP, or Inhba-P2A-tagBFP, were plated separately at 3.0 × 10³ cells per well in NC medium to generate signaling aggregates. After 24 hours, one Proximal signaling aggregate was transferred into each well containing a pre-formed mixed Distal+Proximal aggregate.

Live imaging was performed for 96 hours following aggregate pairing using an incubator confocal microscope (CellVoyager CV1000, Yokogawa). At 96 hours, a single straight line region of interest (ROI) was manually drawn on the Z-stack image in ImageJ/Fiji to traverse from the tagBFP-positive Proximal signaling region into the Distal+Proximal aggregate.

At each pixel position along the line, fluorescence intensities from tagBFP, H2B-Venus, and H2B-tdTomato channels were integrated across an orthogonal sampling line. For each aggregate and channel, the intensity-weighted median position was calculated to define a robust peak location. Distances from the tagBFP median peak to the Distal or Proximal median peaks were computed. Statistical comparisons were performed using a paired Wilcoxon signed-rank test with a one-sided alternative hypothesis (Distal closer than Proximal). Mean normalized intensity profiles with 95% confidence intervals and median peak positions were visualized using ggplot2 in R, with individual biological replicates and combined datasets displayed together.

### Aspect ratio measurement

Serial cryosection images were analyzed in ImageJ/Fiji. For each section image, organoids were segmented and their “Area”, “Feret”, and “MinFeret” were measured. We defined the Feret ratio as MinFeret ÷ Feret (minimum caliper diameter over maximum caliper diameter; range 0–1). For each organoid, we identified the single section with the largest segmented area (proxy for the organoid’s equatorial plane) and took that section’s Feret ratio.

### EdU assay

Cell proliferation was assessed using the Click-iT™ Plus EdU Alexa Fluor™ 647 Flow Cytometry Assay Kit (Thermo Fisher Scientific, Cat. no. C10635) following the manufacturer’s instructions. Briefly, after incubating with 10 µM EdU for 2 h in NC or FW medium, the organoids were digested with Pronase (Millipore, Cat. 537088-10KUCN; 780 U/ml, 10 min, vortex every 5 min). The dissociated cells were fixed, permeabilized, and subjected to the Click-iT™ reaction for fluorescent detection of incorporated EdU. Flow cytometric analysis was performed on a FACS Aria IIIu (BD Biosciences).

### RNA extraction and qPCR

Total RNA was extracted using the RNeasy Micro Kit (74004, QIAGEN) according to the manufacturer’s instructions. For 2D micromass cultures, cells were disrupted by directly applying Buffer RLT to the culture, followed by homogenization with a QIAshredder (79654, QIAGEN).

First-strand cDNA was synthesized from 30 ng total RNA using SuperScript™ II Reverse Transcriptase (Invitrogen, Cat. 18064-014) and random primers, following the manufacturer’s protocol.

Quantitative PCR (qPCR) was carried out using Power SYBR™ Green PCR Master Mix (4368702, Applied Biosystems) on QuantStudio 6 Flex (Thermofisher Scientific). The following primers were used: Mm Gapdh (Forward: 5′-CTCCCACTCTTCCACCTTCG-3′ Reverse: 5′-GCCTCTCTTGCTCAGTGTCC-3′)^84^, MmWnt5a (Forward: 5′-ATGCAGTACATTGGAGAAGGTG-3′ Reverse: 5′-CGTCTCTCGGCTGCCTATTT-3′)^85^. Relative expression was calculated using the ΔΔCt method, with Gapdh serving as the internal control.

### Preparation of DNA-lipid barcodes for scRNA-seq multiplexing

Single-stranded DNAs (ssDNAs), phosphorothioated at the 5′ end, were synthesized in accordance with the barcode flanking sequences used for TotalSeq-B anti-human hashtag antibody (BioLegend). Two barcode oligos were used (N denotes randomized bases; asterisks indicate phosphorothioate bonds in the capture tail):

1. GTGACTGGAGTTCAGACGTGTGCTCTTCCGATCTN₁₀-TGATGGCCTATTGGG-N₉GCTTTAAGGCCGGTCCTAGC*A*A

2. GTGACTGGAGTTCAGACGTGTGCTCTTCCGATCTN₁₀-TTCCGCCTCTCTTTG-N₉GCTTTAAGGCCGGTCCTAGC*A*A

(ssDNA layout: 5′ PCR handle – N₁₀ spacer – 15-nt barcode – N₉ spacer – capture sequence with terminal phosphorothioates.)

Dry oligos were reconstituted in 250 µl 1 M DTT/Tris-HCl and incubated at 37 °C, 400 rpm to deprotect thiols. The reaction was brought to 1 ml by adding 750 µl 1 mM EDTA/PBS, then applied to a NAP-10 columns (GE Healthcare) (equilibrated with 3 ml 1 mM EDTA/PBS, repeated 5×). Elution was performed with 200 µl fractions of 1 mM EDTA/PBS (8 sequential fractions). Each fraction was diluted 1:100 and A260 measured using NanoDrop One(Thermofisher); high-A260 fractions were pooled for subsequent conjugation. DNA concentration after purification/reaction was estimated as:

C_post (µg ml⁻¹) = C_pre × (A260_post / A260_pre),

Where C_pre and A260_pre are the concentration and absorbance before column purification, and A260_post is the absorbance of the pooled, purified material.

Maleimide–polyethyleneglycol(PEG)–1,2-dipalmitoyl-sn-glucero-3-phosphoethanolamine (DPPE) (prepared from maleimide–PEG–NHS and DPPE) was reconstituted to 5 mg/ml in 1 mM EDTA/PBS. The deprotected, purified DNA was mixed at equimolar ratio with maleimide–PEG–DPPE and incubated overnight at 25 °C, 400 rpm to yield DNA–PEG–DPPE.

### Computer simulations and visualization

All simulations are implemented in Python. The Ordinary Differential Equations that constitute the ABM are solved using a first-order explicit Euler scheme. The reaction-diffusion scheme uses a standard five-point stencil discretization of the diffusion operator on a square grid. The discretized equations are then solved using a first-order in time explicit scheme. To accelerate the computations, data is encoded as PyTorch tensors^86^ on the GPU. Cell-cell interactions are implemented using the KeOps framework^87^ and the SiSyPHE library^88^. The coarse-grained PDE models (Cahn-Hilliard equation) are solved using the Finite Element Method in FEniCS^89^. The 3D and 2D visualization rely on VTK and Paraview^90^.

The source code is freely available at https://github.com/antoinediez/PyLimbMorph

## Declarations

The authors used ChatGPT (OpenAI) for language-editing and formatting assistance. The authors reviewed and edited all AI-assisted text and take full responsibility for the manuscript.

## Data and code availability

Single-cell RNA-seq data, including raw FASTQ files, processed data files, and cell-level sample-assignment metadata, have been deposited in the Gene Expression Omnibus (GEO) under accession number GSE331455. PIC transcriptomic data, including raw FASTQ files and processed differential-expression tables, have been deposited in GEO under accession number GSE331258. Source-data tables underlying quantitative plots and custom scripts used for transcriptomic analyses and figure generation are available at Zenodo draft record under DOI: 10.5281/zenodo.20280291. Reviewer access is provided through https://zenodo.org/records/20280291?preview=1&token=eyJhbGciOiJIUzUxMiJ9.eyJpZCI6IjZjNmRjNGZlLWZjMzEtNGI4ZC1iNGJlLTRhODAwNzZiYjkzYyIsImRhdGEiOnt9LCJyYW5kb20iOiIyODdiMzdlYzFhYzE1M2E3NGJhN2I1ZDA5ZDNhYzZmMiJ9.vyU1iKIwMiKJT3R1K8xRkDa13PdPF3F1ZFHJq99wwhjsCga-e5LBneQLcoDWMg40SVHemWeSr_iKr_QM5RjaYw. The record will be made public upon publication.

Simulation code for the agent-based and continuum modeling is available at https://github.com/antoinediez/PyLimbMorph, which includes a minimal example for inspection. Raw microscopy images and replicate datasets used for image quantification are available from the corresponding authors upon reasonable request because of their large file size. Materials generated in this study are available from the corresponding authors upon reasonable request, subject to institutional and material transfer agreement requirements.

## Supporting information

Supplementary text for scRNA-seq

Supplementary Text on Mathematical Theory

Movie 1. Limb-mesenchymal organoid: culture conditions, Related to Figure 1B.

Movie 2. Limb-mesenchymal organoid: the distal:proximal labelling, Related to Figure 1E.

Movie 3. 2D micromass: conditions, Related to Figure S3B.

Movie 4. Simulation: differential adhesions, Related to Figure 2B.

Movie 5. Simulation: differential adhesions+Fgf8b-driven chemotaxis, Related to Figure 2H.

Movie 6. Limb-mesenchymal organoid: merging with chemoattractant aggregate, Related to Figure 2E and S5D.

Movie 7. Limb-mesenchymal organoid: overexpression of the distal morphogens, Related to Figure 3C.

Movie 8. Simulation: differential adhesions+Fgf8b-driven chemotaxis+Wnt5a-driven convergent extension, Related to Figure 3E.

Supplemental Table1: HyBISS probes

## Acknowledgments

We thank all members of the Eiraku Lab for helpful discussions; Dr. Hitomi Watanabe for generating Sox9–GFP mouse embryos; and Dr. Ko Sugawara for technical support with the ELEPHANT plug-in; and Dr. Elazer Zelzer for sharing scRNAseq data. We thank Single-cell Genome Information Analysis Core (SignAC) at WPI-ASHBi (Kyoto University) for their support. We also thank Dr. Spyros Goulas at WPI-ASHBi (Kyoto University), Dr. Alberto Roselló-Díez (University of Cambridge), and Dr. Kimberly L. Cooper (University of California, San Diego) for critical reading and constructive suggestions on our manuscript.

## Funding

This work is supported by Fusion Research Grant, ASHBi (FY2023-2024: to R.T, S.P., and A.D.), Grant-in-Aid for Research Activity Start-up, Japan Society for the Promotion of Science (JSPS) (JP20K22652 to R.T.), Grant-in-Aid for Early-Career Scientists (JP22K15122 to R.T., JP23K13015 to A.D., JP24K16962 to S.P.), Precursory Research for Embryonic Science and Technology (PRESTO), Japan Science and Technology Agency (JST) (JPMJPR2537 to R.T.), ISPF - International Collaboration Awards 2024 Round 2 (Japan), The Royal Society, UK (ICA\R2\242121 to R.T), Grant-in-Aid for Transformative Research Areas, Ministry of Education, Culture, Sports, Science, and Technology (MEXT), Japan (JP23H04933 to M.E., JP22H05110 supporting S.P.), and Core Research for Evolutional Science and Technology (CREST, JST) (JPMJCR12W2 to M.E.).

## Authors information

### Contributions

R.T. conceived, designed the study and performed experiments. A.D. and S.P. performed simulations and mathematical analyses. R.K. and S.O. performed PIC experiments. K.T. performed doublet analysis. M.H. and R.N. performed scRNA-seq multiplexing. H.A. provided the Sox9-GFP mouse line. R.Takada and S.T. produced the Wnt5a antibody. Cauchy–Green tensor analysis: S.P. and M.M., with initial analyses by M.M., supervised by J.S., who also provided critical scientific feedback and discussions. M.E. conceived, designed and supervised the study. R.T., A.D., and S.P. wrote the manuscript with input from all authors.

## Ethics declarations

### Competing interests

Authors declare no conflict of interests.

## Supplementary Materials

Supplementary Text: scRNA-seq

Supplementary Text: Cauchy-Green Tensor Estimation

Supplementary Text: Mathematical Theory

## Figure Legends

Table S1: HybISS probes Movies 1 to 8

**Movie 1. Limb-mesenchymal organoid: culture conditions, Related to Figure 1B**. Limb mesenchyme from Sox9–KI–EGFP E11.5 mouse embryos is aggregated and cultured for 193 h in NC or FW (Fgf8+Wnt3a) conditions, initiated with 9,000 or 27,000 cells; Sox9-GFP marks cartilage (green) on bright-field.

**Movie 2. Limb-mesenchymal organoid: the distal:proximal labelling, Related to Figure 1E**. Distal–proximal lineage labeling. The cells from the distal half of autopod are labeled H2B-Venus (green) and proximal cells H2B-tdTomato (magenta) before aggregation, and cultured in an organoid for 76 h in NC and FW, started from 9,000 or 27,000 cells, overlaid with bright-field. Tips were enriched in Distal cells.

**Movie 3. 2D micromass: conditions, Related to Figure S3B.** The cells from the distal and proximal half of autopod of Sox9–KI–EGFP (white) E11.5 mouse embryos are labeled before aggregation, and cultured as micromass culture. (B) Micromass culture of limb mesenchyme in NC and FW at 72 h; Sox9-GFP in white; Distal and Proximal cells labeled with H2BtdTomato (green) and H2BtagBFP (magenta).

**Movie 4. Simulation: differential adhesions, Related to Figure 2B**. ABM implementing differential adhesion only (DD>DP>PP). Right: Fgf8b and Wnt5a fields.

**Movie 5. Simulation: differential adhesions+Fgf8b-driven chemotaxis, Related to Figure 2H**. ABM implementing differential adhesion + traction bias toward the Fgf8b gradient. Right: Fgf8b and Wnt5a fields.

**Movie 6. Limb-mesenchymal organoid: merging with chemoattractant aggregate, Related to Figure 2E and S5D.** A proximal signaling aggregate (tagBFP or Fgf8b-P2A-tagBFP, blue) is assembled with a Distal (Green) + Proximal (Magenta) aggregate and cultured in NC condition for 96 h.

**Movie 7. Limb-mesenchymal organoid: overexpression of the distal morphogens, Related to Figure 3C**. Overexpression of tagBFP or Wnt5a-P2A-tagBFP (blue) in Proximal cells, cultured in FW condition for 96h.

**Movie 8. Simulation: differential adhesions+Fgf8b-driven chemotaxis+Wnt5a-driven convergent extension, Related to Figure 3E**. ABM combining differential adhesion (DD>DP>PP), DP bias toward Fgf8b, Distal Wnt5a production, and DD bias orthogonal to Wnt5a. Right: Fgf8b and Wnt5a fields.

---

**Figure S1.**
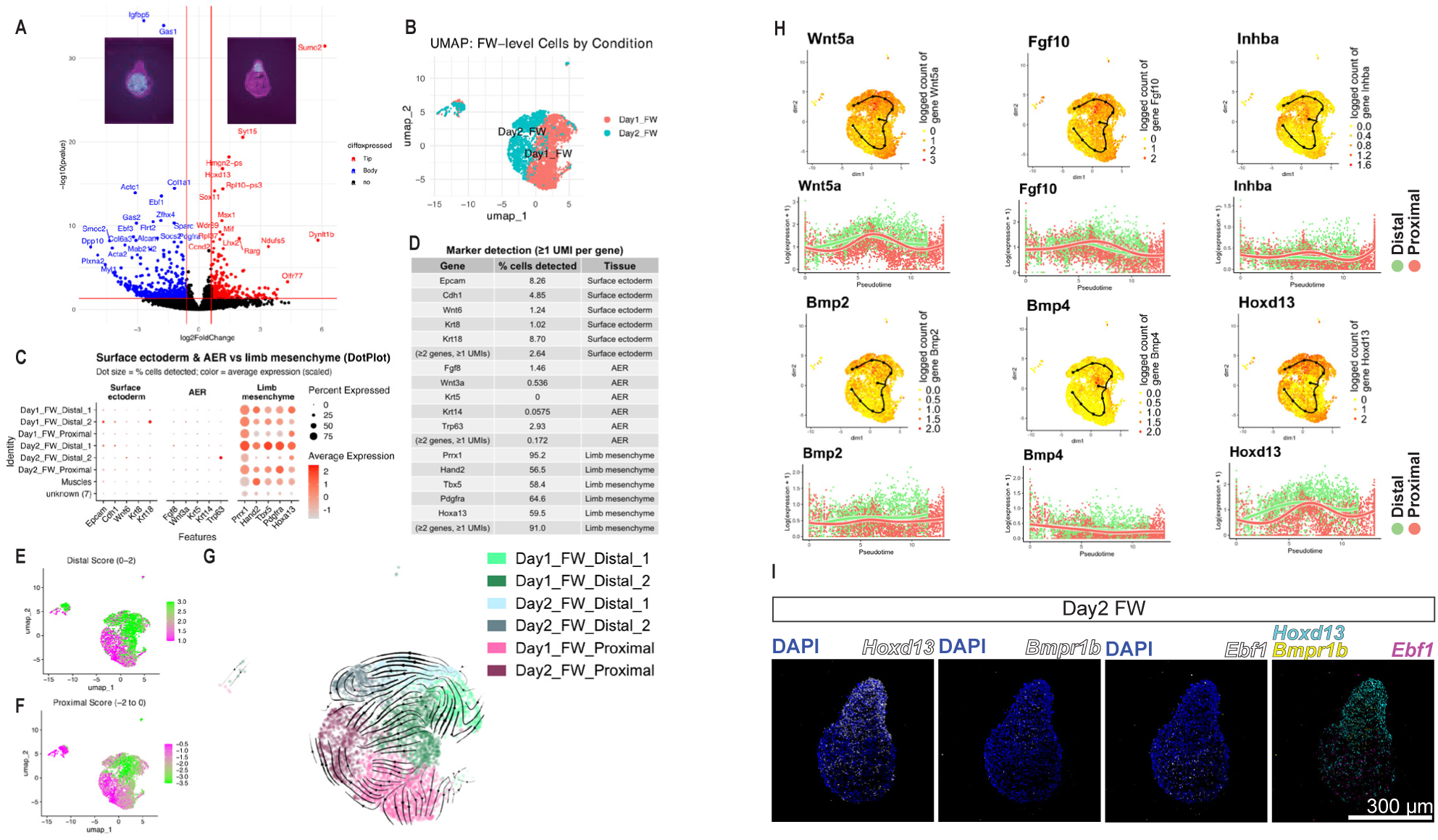
Distal–Proximal gene programs in limb-mesenchymal organoids (FW). (A) Volcano plot comparing tip (Distal) vs bottom (Proximal) PIC profiles; x = log₂FC, y = −log₁₀(p). Genes called Tip (red) if log₂FC > 0.6 & p < 0.05, Bottom (blue) if log₂FC < −0.6 & p < 0.05; others black. Vertical lines ±0.6 (∼1.5-fold); horizontal line p = 0.05; significant genes labeled. (B) UMAP of scRNA-seq: Day 1 (red), Day 2 (blue). (C) Marker table for surface ectoderm, AER, limb mesenchyme: % cells with ≥1 UMI; summary row shows % cells with ≥2 markers/tissue. (D) DotPlot of these markers across clusters (Day1/Day2 Distal/Proximal; muscles): dot size = %≥1 UMI; color = scaled mean. (E–F) UMAP FeaturePlots of Distal_Score (average of top 20 tip-enriched genes from A; magenta→green) and Proximal_Score (top 20 bottom-enriched; green→magenta). (G) RNA-velocity streamlines (scVelo). (I) (J) For selected genes (*Wnt5a*, *Fgf10*, *Inhba*, *Bmp2/4*, *Hoxd13*): top, UMAP colored by log(expression+1) with Slingshot curves, rooted as in G, with endpoints Day2_FW_Distal_2 (lineage 1: Distal) and Day2_FW_Proximal (lineage 2: Proximal); bottom, log(expression+1) vs Slingshot pseudotime per lineage (Distal: purple, Proximal: yellow) with tradeSeq NB-GAM smoothers. (I) Expressions of *Hoxd13*, *Bmpr1b*, and *Ebf1* in FW organoids (Raw images of Figure 1K).

**Figure S2.**
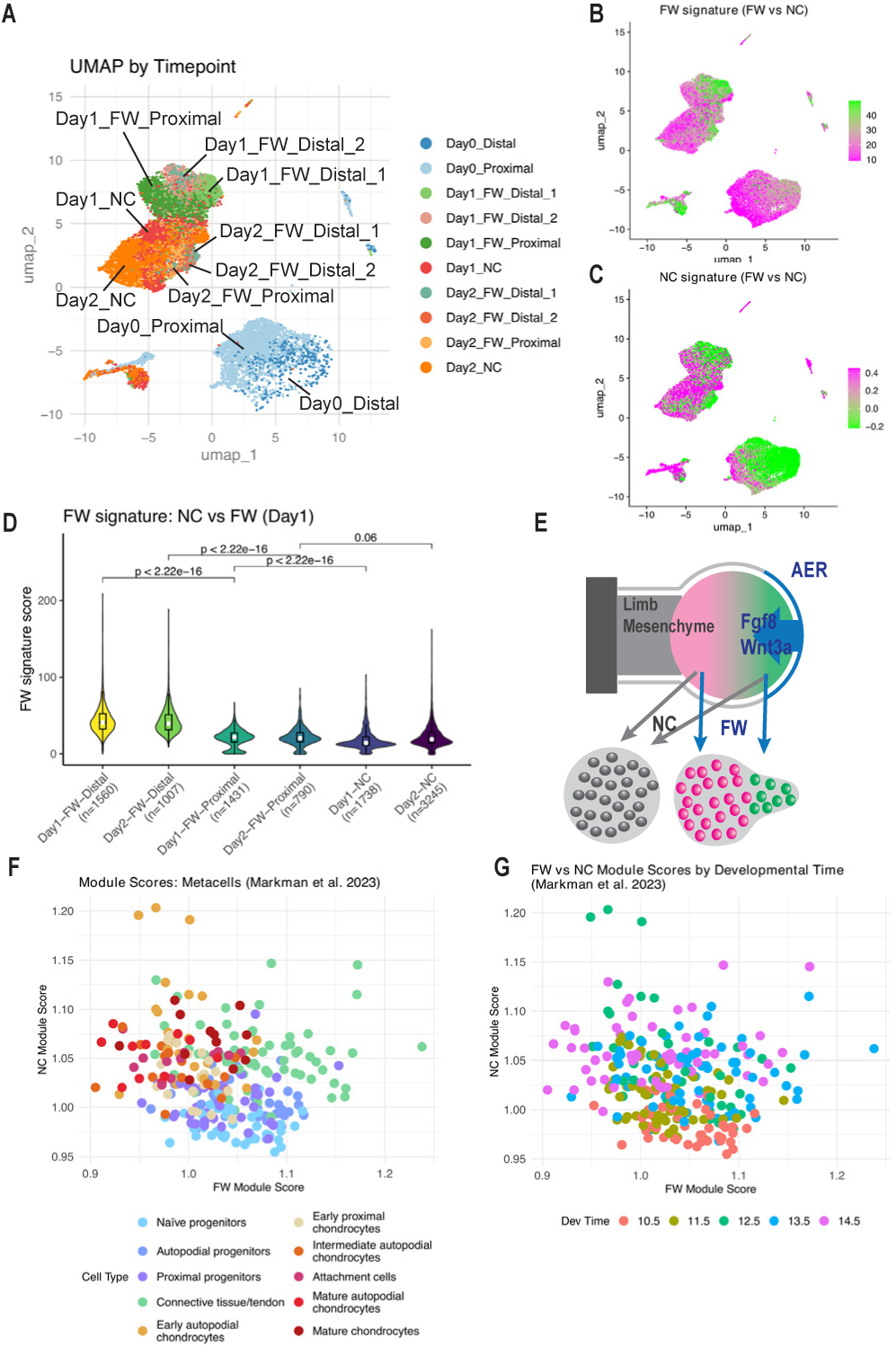
FW condition maintains Distal identity and an “active” state. (A) UMAP of limb-mesenchymal organoids (FW/NC; Day 1/Day 2) together with E11.5 Distal/Proximal cells (Day 0). (B–C) FeaturePlots of per-cell module scores on the UMAP. The FW signature is the average expression of the top 50 genes up-regulated in Day1_FW vs Day1_NC (Wilcoxon FindMarkers, padj < 0.05); the NC signature is the corresponding NC-up set. Scores are shown per cell with truncated scales (10th–90th percentiles): (B) FW signature (green→magenta), (C) NC signature (magenta→green). (D) Violin/box plots of the FW signature across binned clusters (Day1-/Day2-FW-Distal, Day1-/Day2-FW-Proximal, Day1-/Day2-NC); white dots mark medians. Pairwise two-sided Wilcoxon rank-sum tests; Benjamini–Hochberg–adjusted p-values are shown above brackets. x-axis labels include per-bin sample sizes (n); values are module-score units. (E) Schematic: FW sustains Distal identity and an “active” program. (F–G) Metacell scatter: x = FW module score, y = NC module score. (F) Colored by metacell class defined in Markman et al. 2023. (G) Same scatter colored by developmental time defined in Markman et al. 2023 (E10.5–E14.5; majority vote per metacell).

**Figure S3.**
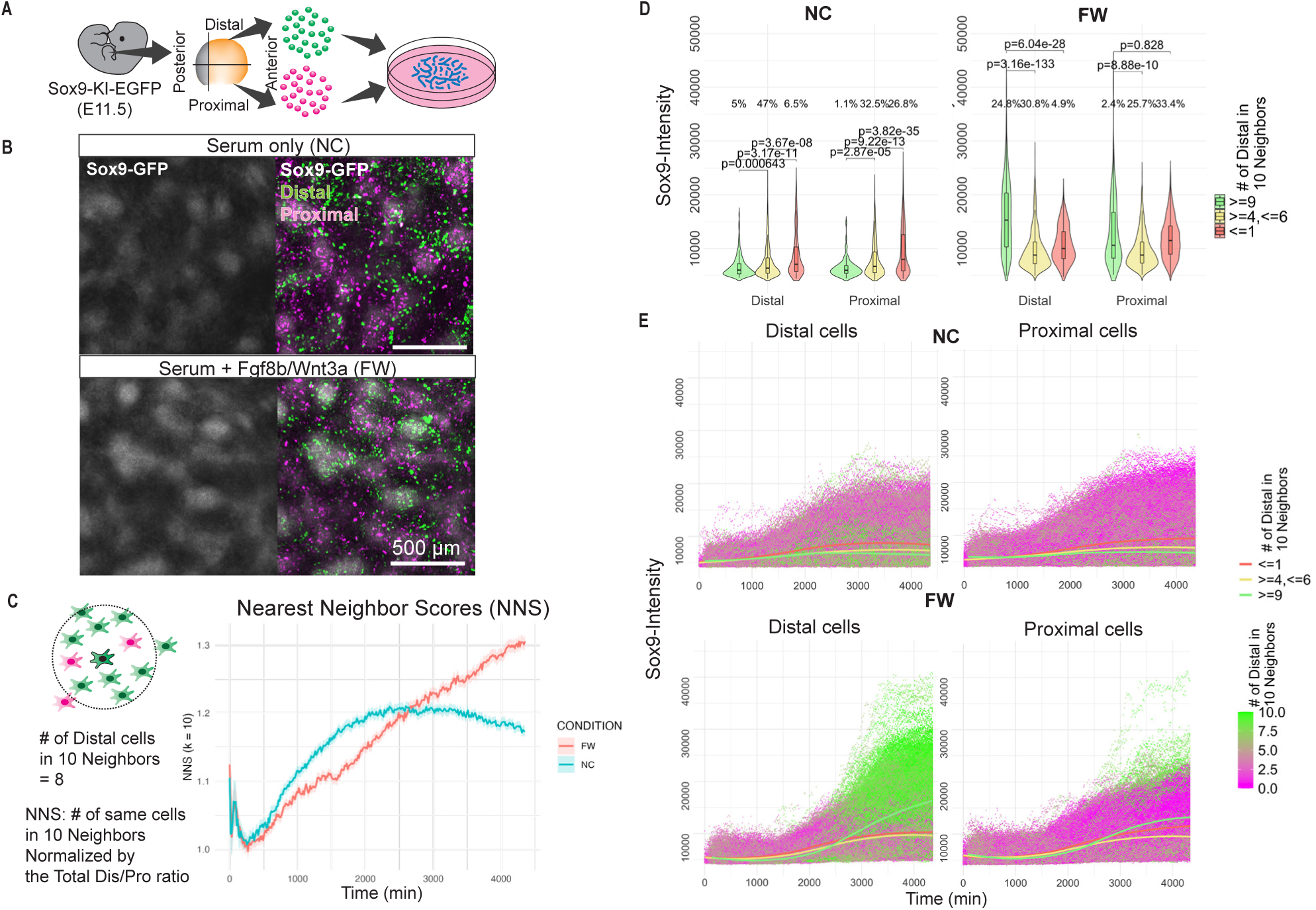
Variations of limb-mesenchymal organoid cultures. (A) Schematics: Distal– proximal lineage labeling. The cells from the distal and proximal half of autopod of Sox9–KI– EGFP (white) E11.5 mouse embryos are labeled before aggregation, and cultured as micromass culture. (B) Micromass culture of limb mesenchyme in NC and FW at 72 h; Sox9-GFP in white; Distal and Proximal cells labeled with H2BtdTomato (green) and H2BtagBFP (magenta). (C) Dot plots of Sox9-GFP intensity over time in Distal (left) and Proximal (right) cells; points colored by the number of Distal neighbors among the 10 nearest. (D) Violin/box plots at 72 h comparing neighbor groups (≥9 vs ≤1 Distal neighbors) within each cell type using two-tailed unpaired Student’s t-tests; significance shown as asterisks (*p*<0.05, \**p*<0.01, \*\**p*<0.001). Percentages per group are shown below; sample sizes (*n*) and means are annotated. (E) Average Nearest Neighbor Score (NNS, k=10) over time in NC and FW, defined as the proportion of same-type neighbors normalized by cell-type frequency; lines show means, ribbons show bootstrapped 95% CIs.

**Figure S4.**
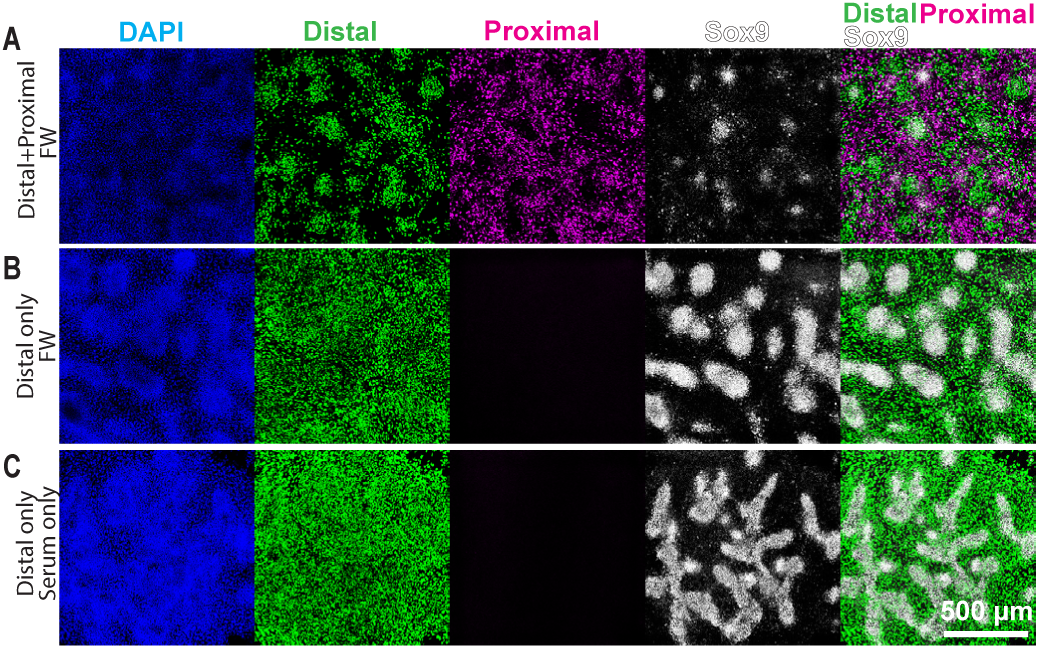
2D micromass culture in various cell types and culture conditions. (A) Distal + Proximal cells in FW, (B) Distal cells in FW, (C) Distal cells in NC at 72 h. DAPI (blue), Distal (green), Proximal (magenta), Sox9–GFP (white). Scale bar, 500 µm.

**Figure S5.**
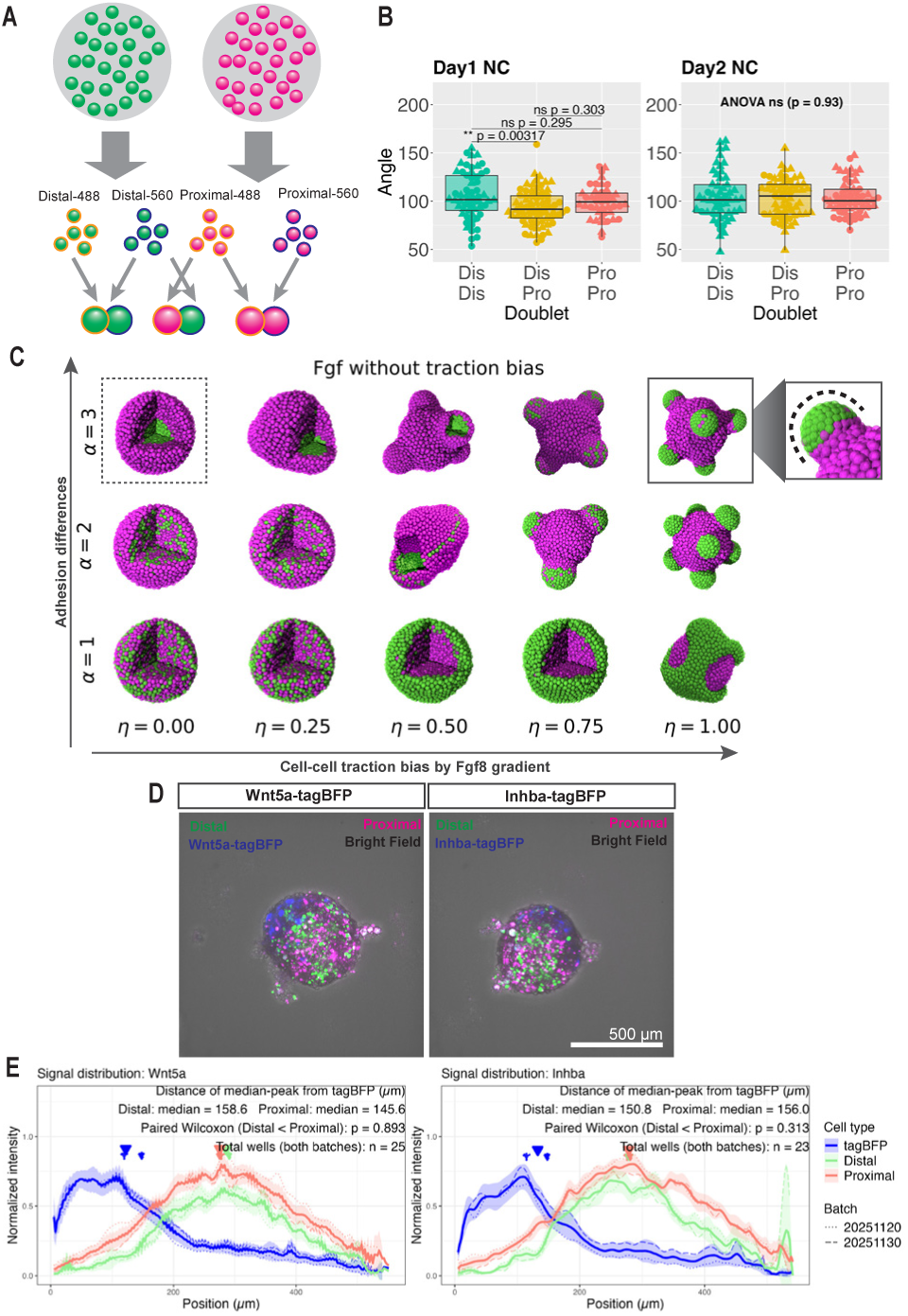
Details on the mechanisms of the morphological symmetry breaking. (A) Doublet assay workflow. Distal and Proximal cell aggregates were cultured in FW, dissociated on Day 1 or Day 2, and membrane-labeled with MemGlow dyes (488 or 560). Doublets were then assembled in predefined pairings: Distal_488–Distal_560 (DD), Distal_560–Proximal_488 (DP), and Proximal_488–Proximal_560 (PP). (B) Beeswarm+box plots of Angle by doublet class (DD, DP, PP) for Day1/Day2 NC. Dots = doublets; shape = batch. Stats: one-way ANOVA per experiment; if significant (α=0.05), Tukey HSD shown. (C) ABM with variable parameters of differential adhesion (DD > DP > PP) and a traction bias toward Fgf8b (outer→inner). Axes: α, adhesion contrast; η, traction bias. Dotted box: adhesion-only. Solid box: adhesion + traction with an inset showing outline of a protrusion (dotted). (D) Live imaging of assembly assay at 96h: a proximal signaling aggregate (Wnt5a-P2A-tagBFP or Inhba-P2A-tagBFP) assembled with a Distal+Proximal aggregate. (E) Signal distribution at 96 h. Intensities for tagBFP, H2B-Venus (Distal), and H2B-tdTomato (Proximal) were integrated along a line from the tagBFP region; intensity-weighted medians defined peaks. Distances from the tagBFP peak to Distal vs Proximal peaks were compared by paired one-sided Wilcoxon (hypothesis: Distal closer). Plots show mean normalized profiles ±95% CI with median markers.

**Figure S6.**
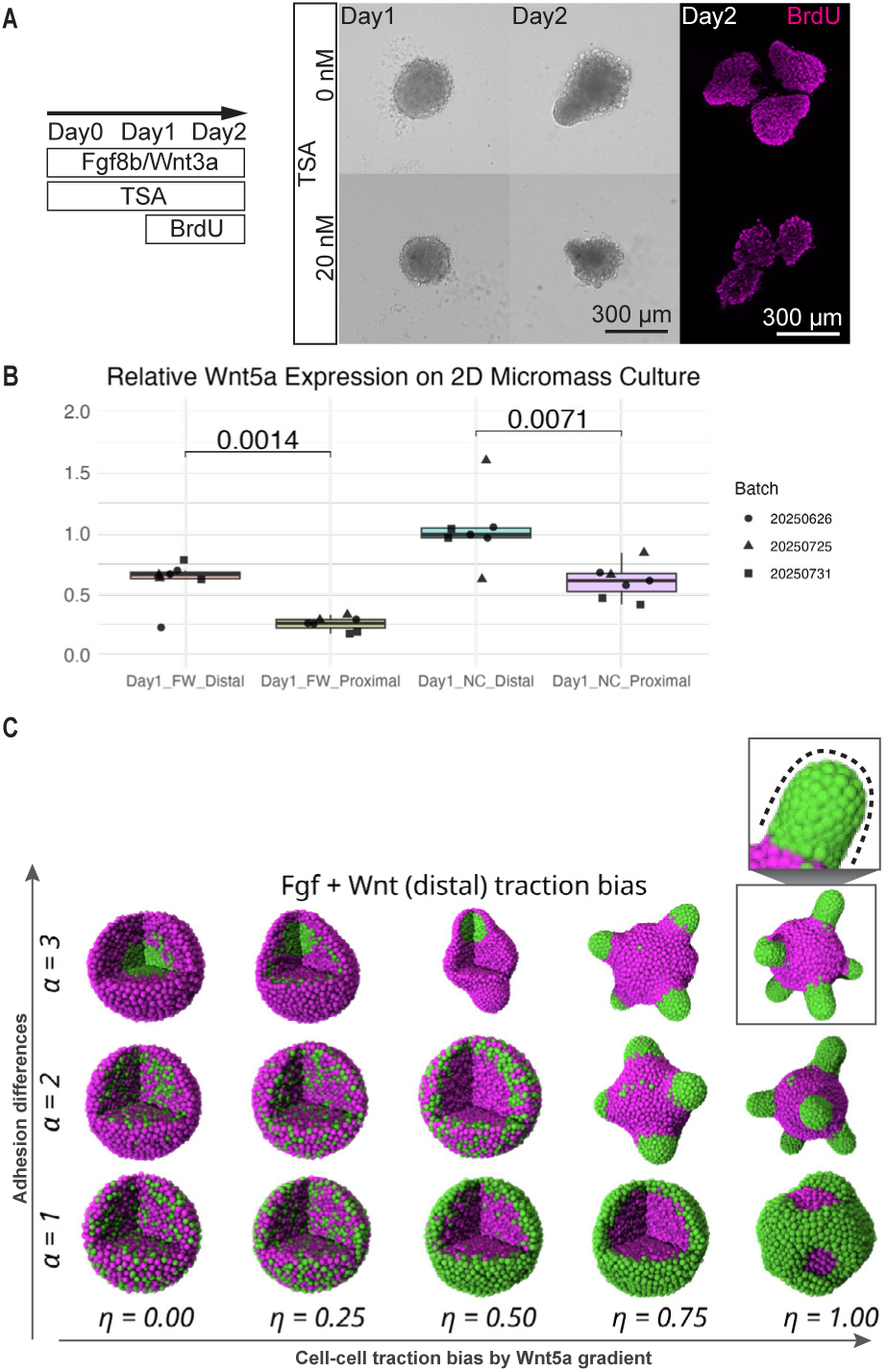
Tissue elongation is independent of biased proliferation, and Wnt5a is enriched in Distal cells. (A) Inhibitor treatment for cell proliferation. Experimental schedule (left): organoids were cultured in FW (Fgf8b/Wnt3a) from Day0–Day2; TSA (HDAC inhibitor) was added Day1–Day2; BrdU was pulsed Day1-Day2. Bright-field images at Day1 and Day2 are shown for 0 nM and 20 nM TSA, with BrdU immunostaining (magenta) on Day2 (right). scale bars, 300 µm. (B) Relative Wnt5a expression (qPCR) in 2D micromass, Day 1. Box–dot plots display Day1_FW_Distal, Day1_FW_Proximal, Day1_NC_Distal, and Day1_NC_Proximal. Each dot is one biological replicate; the y-axis shows expression normalized to Gapdh. Statistics: one-way ANOVA across the four groups with Tukey’s HSD for pairwise comparisons. Planned Distal vs Proximal contrasts within FW and within NC were additionally tested by two-sided t-tests; the unadjusted p values are printed on the plot. (C) ABM combining differential adhesion (DD>DP>PP), DP bias toward Fgf8b, Distal Wnt5a production, and DD bias orthogonal to Wnt5a, with variable parameters. Parameters: traction bias (*η)* vs adhesion differences (*α*) solid box marks protrusion shape (dotted line in inset).

**Figure S7.**
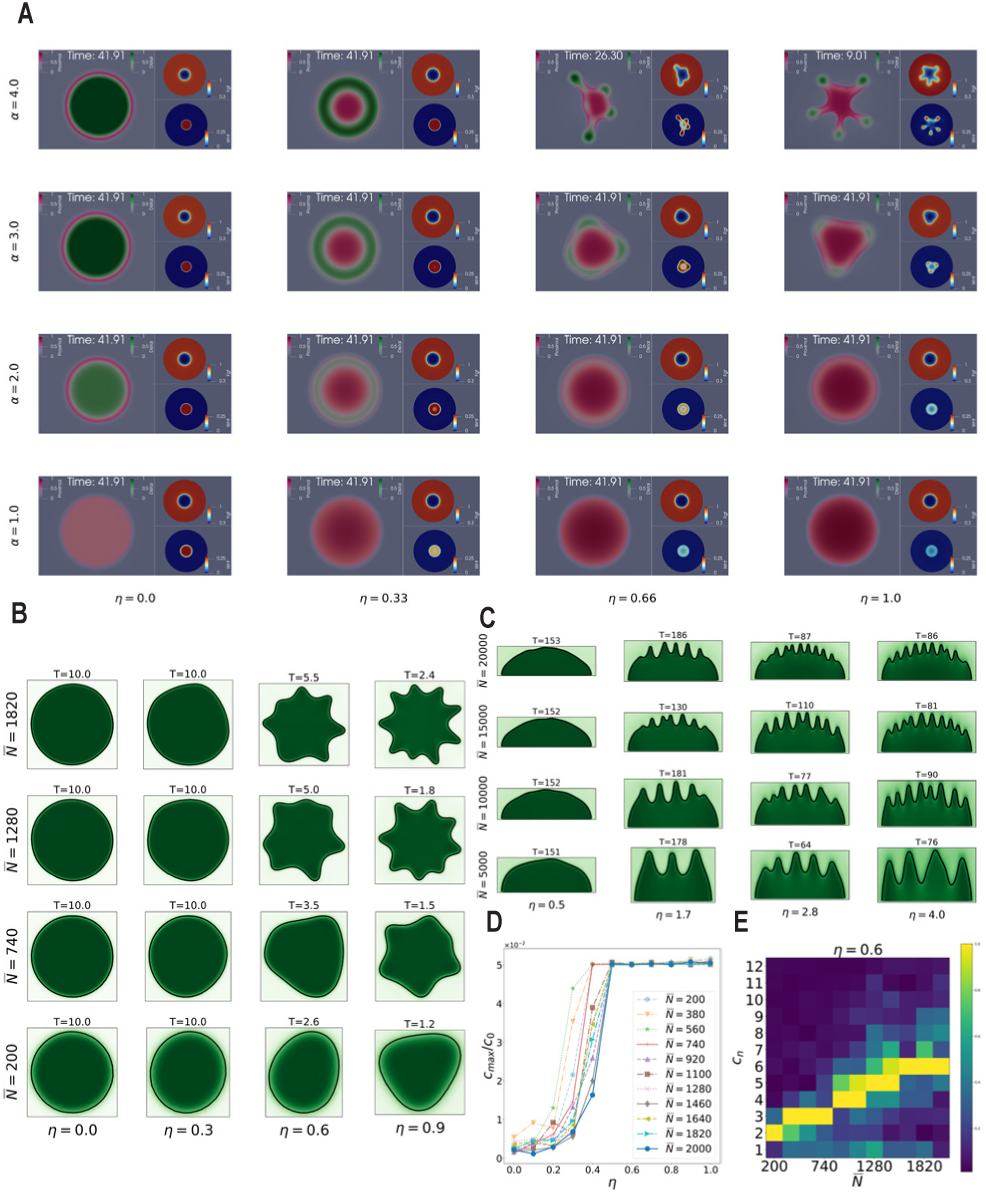
PDE simulations with variable parameters. (A) Simulations of equations from Figure 6D coupled with the model in Figure 5D and 5E with traction bias *η*∈{0, 0.33, 0.66, 1} and differential adhesion *α*∈{1, 2, 3, 4} (see Figure 6F). Setting identical to the ABM version (Figure S6C). Simulations are stopped when instabilities have developed or at the arbitrary endpoint *T*=42. (B) Simulations of the Distal tissue only as stated in Figure 6E initiated from a sphere, with constant radial and various traction bias values and tissue sizes *<u>N</u>*. Simulations are stopped when instabilities have developed or at the arbitrary endpoint *T*=10. (C) Same as Figure 6G for various traction bias values *η* and tissue sizes *<u>N</u>*. (D) Normalized value *c_k_*/*c_0_* of the largest Fourier coefficient of each contour as a function of *η* in the setting of Figure S7B. Instabilities develop if *c_k_*/*c_0_* reaches the arbitrary cap value 0.05. (E) In the setting of Figure S7B, heatmap of the values of the first 12 Fourier modes of the contours as a function of *<u>N</u>* for *η* = 0.6 at the endpoint of the simulation. The number of protrusions corresponds to the maximal Fourier mode, normalized to 1.

**Figure S8.**
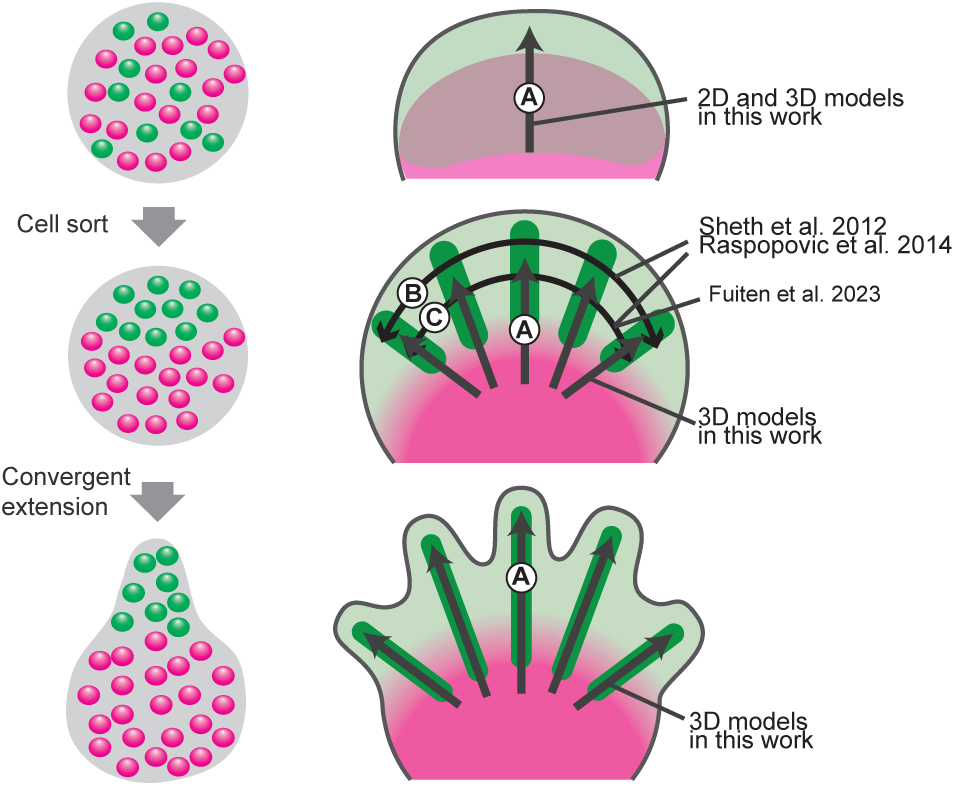
Biological interpretation of the existing and current model. 2D micromass culture of distal and proximal cells in FW condition in Figure S4A corresponds to establishment of the distal autopod domain and elongation of digits along the proximal–distal (PD) axis. 2D micromass culture of distal cell only in FW (Figure S4B) and NC (S4C) condition correspond to the periodic patterning of digit and interdigital tissues along the anterior–posterior axis at, respectively, the distal-most part (FW: longer wavelength) and a more proximal region (NC) of the distal autopod.

## Notes

https://github.com/antoinediez/PyLimbMorph

https://zenodo.org/records/20280291?preview=1&token=eyJhbGciOiJIUzUxMiJ9.eyJpZCI6IjZjNmRjNGZlLWZjMzEtNGI4ZC1iNGJlLTRhODAwNzZiYjkzYyIsImRhdGEiOnt9LCJyYW5kb20iOiIyODdiMzdlYzFhYzE1M2E3NGJhN2I1ZDA5ZDNhYzZmMiJ9.vyU1iKIwMiKJT3R1K8xRkDa13PdPF3F1ZFHJq99wwhjsCga-e5LBneQLcoDWMg40SVHemWeSr_iKr_QM5RjaYw

### Competing Interest Statement

The authors have declared no competing interest.

### Summary of Updates

Updated on the data, discussions, and supplementary movies

