## Supplementary text for scRNA-seq for "Self-organized mechanochemical instabilities drive the emergence of digit tissue morphogenesis"

### Supplementary Text: scRNA-seq

To analyze the expression dynamics of the secreted molecules from Distal cells, we first focused on scRNA-seq profiles from digit organoids grown in FW condition collected on Day 1 and Day 2 (Day1_FW and Day2_FW), which were segregated on UMAP space (Figure S1B). To confirm that epidermal contribution was effectively minimized, surface ectoderm and AER markers (*Epcam, Cdh1, Wnt6, Krt8/18, Fgf8, Wnt3a, Krt5/14, Trp63*) were detected in only a very small minority of cells, with only 2.64% and 0.172% of cells expressing more than 2 surface ectoderm or AER markers, respectively. In contrast, mesenchymal limb bud markers (*Prrx1, Hand2, Tbx5, Pdgfra, Hoxa13*) were robustly expressed in 55–95% of cells, and 91% of cells expressed more than two mesenchymal markers (Figure S1C, D). To characterize Distal and Proximal cells, we picked up top 20 differentially expressed genes (DEGs) between the tip and the bottom of the organoid from the PIC experiments (Figure S1A) and computed the normalized average expression levels of those genes as "Distal Score" and "Proximal Score", respectively (Figure S1E, F). As the results suggested that the cell clusters in the top half on the UMAP were Distal cells, we annotated Day1_FW_Distal, Day2_FW_Distal, Day1_FW_Proximal, and Day2_FW_Proximal clusters on the UMAP (Figure 1G). Based on this information, we generated the pseudo-trajectory using RNA velocity and slingshot (Figure S1G and 1G). Subsequently, we inferred the expression dynamics of the genes that are upregulated in the Distal lineage, such as *Bmp2, Bmp4, Hoxd13, Wnt5a, Fgf10*, and *Inhba* (Figure S1H).

Next, to get some insight into the role of FW in in vitro culture, we mapped the cell clusters annotated above onto a larger data set including Distal and Proximal limb autopod mesenchyme at E11.5 (Day0_Distal, Day0_Proximal), which are the source cells of the organoids, as well as the cells from organoids grown in NC condition collected on Day1 and 2 ( Day1_NC, Day2_NC) (Figure S2A). We picked up the top 50 DEGs between {Day1_FW and Day2_FW} vs {Day1_NC and Day2_NC} and computed the normalized average expression levels of those genes as "FW signature" and "NC signature". In both plots, we validated that the cells high in the FW signature and low in the NC signature (both colored in green) overlapped with the cells cultured in FW conditions (Figure S2B, C). Interestingly, Day0_Distal cells also turned on green on both plots, suggesting that the difference between the cells cultured in FW and NC conditions correlated with the distal-proximal difference in the limb bud (Figure S2B, C). The violin plot for the FW signature further showed that FW_Distal are highest in the FW signature, FW_Proximal cells come next, and NC cells are lowest (Figure S2D). These results implied that FW, namely AER signals maintain the cells as Distal and "naïve"state, while the cells deprived of AER signals become Proximal and differentiated state (Figure S2E).

Re-analysis of the scRNA-seq data from Markman et al. 2023 (Markman et al., 2023) further confirmed that the metacells assigned as "Naive progenitors," "Autopodial progenitors", were "Proximal progenitors" were high in FW signature and low in NC signature, while the cells assigned as "Mature chodrocytes", "Mature autopodial chodrocytes", and "Intermediate autopodial cells" were high in NC signature and low in FW signature (Figure S2F). The correlation of FW and NC signatures with "active" and "inactive" is also represented by the plot showing the metacells assigned as earlier stages were higher in FW signature and lower in NC signature (Figure S2G). Taken together, our scRNA-seq results suggested that FW conditions maintained the Distal and Proximal identities as undifferentiated mesenchyme while NC conditions drive both of the cells towards a more Proximal and differentiated state (Figure S2E).
