## Supplementary Text on Mathematical Theory for "Self-organized mechanochemical instabilities drive the emergence of digit tissue morphogenesis"

Supplementary Material

### Derivation of the mathematical models in *"Self-organized mechanochemical instabilities drive the emergence of digit tissue morphogenesis"*

R. Tsutsumi, A. Diez, S. Plunder et al.

#### 1 Summary of the models

This supplementary contains the mathematical definition and the derivation of the two main models, ABM and PDE, shown respectively in Figs.(M) 5 and 6 in the main text. From now on, the figures in the main text are referred using the notation (M) to distinguish them from the figures in this document. While the ABM is simply defined by a set of ordinary differential equations (shown in Section 2), it takes several steps to formally derive the final PDE model. This leads to several intermediate models that are summarized in Fig. 1. Our approach and the structure of this supplementary is explained below.

- The first step is the so-called mean-field limit that leads from the ABM to a first PDE model when the number of cells is taken to the infinity (Section 3.1).
- We consider a space-time rescaling corresponding to a large organoid observed during a long time scale (Section 3.2).
- At this point, the PDE is said to be non-local and is not prone to numerical simulations or theoretical analysis. The most technically demanding part is to formally compute a so-call localization limit that can be understood as taking the asymptotic limit of the previous space-time rescaling factor (Section 4).
- After the localization limit, we make a few simplifying assumptions to obtain the final PDE model (Section 5).

- The final model contains two major components that are analyzed independently for a better understanding: a two-population Cahn-Hilliard term leads to cell-sorting phenomena (Section 6); it is combined with a nonlinear anisotropic diffusion term that is responsible for the emergence of novel fingering patterns (Section 7). The combination of both elements leads to the full model that can be compared to the ABM simulations and the organoid experiments (Section 8).

In addition, a discussion of some open mathematical problems is presented in Section 9 and the detailed parameter values of the various models and numerical experiments can be found in Section 10.

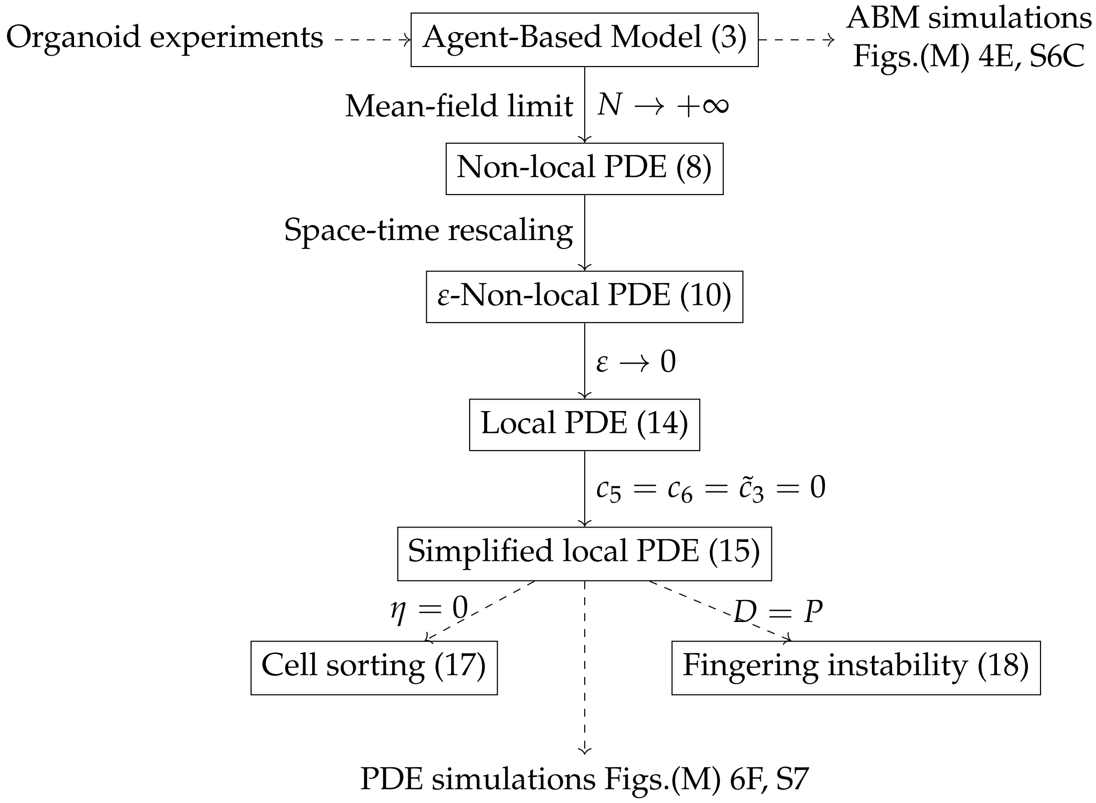


**Figure 1:** Summary of the models.

#### 2 Agent-Based Model

We consider $N = N_D + N_P$ cells where $N_D$ and $N_P$ respectively denote the number of distal and proximal cells. Cells are treated as punctual particles located at positions $x_1, \ldots, x_N$ in the space $\mathbb{R}^d$ of dimension $d$, with two interaction radii $\delta < R$, corresponding, respectively, to the repulsion and attraction ranges. The attraction and repulsion forces are binary forces along each of the pairwise connecting vectors $z_{ij} = x_j - x_i$ as shown in Fig. 2.

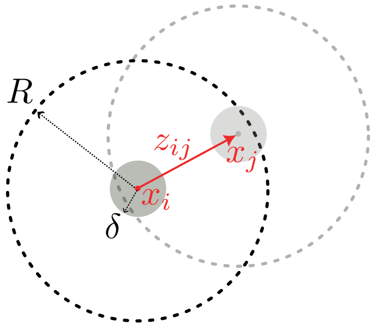


**Figure 2:** Two particles with repulsion radius $\delta$ and attraction radius $R$.

#### 2.1 Attraction forces

Given a connecting vector $z$ and a biased direction $w_{XY} \in \mathbb{R}^d$ with Euclidean norm $|w_{XY}| \leq 1$, we consider attraction forces of the form

$$F_{X \leftarrow Y}(z, w_{XY}) := \xi_{XY} \big(1 + \eta_X^{XY} \langle \hat{z}, w_{XY} \rangle\big) \big(1 - \eta_X^{XY} \langle \hat{z}, w_{XY} \rangle\big) K(z) z. \tag{1}$$

The meaning of each of the terms and symbols appearing in Eq. (1) is explained below.

- From now on, $X$ and $Y$, as index or exponent, are placeholders for the two cell types $D$ (distal) and $P$ (proximal).
- The force $F_{X \leftarrow Y}$ is the force exerted on a cell of type $X$ by a cell of type $Y$.
- The scalar $\xi_{XY} > 0$ is the base adhesion magnitude between cell types $X$ and $Y$.
- The vector $\hat{z} = z/|z|$ is the normalized connecting vector and $\langle \hat{z}, w \rangle$ denotes its Euclidean dot product with the direction $w$, which, in the following, will correspond to the (normalized) gradient of Wnt5a concentration.
- The scalar $\eta_X^{XY} \in [0, 1]$ is the traction bias magnitude of a cell of type $X$ when it interacts with a cell of type $Y$. Note that in most cases, $\eta_X^{XY}$ does not actually depend on $Y$ and will thus simply be denoted by $\eta_X$. In particular, under the Planar Cell Polarity (PCP) hypothesis, $\eta_X = 1$ corresponds to completely polarized cells of type $X$, with no adhesion receptors at their rear end, while $\eta_X = 0$ corresponds to non-polarized cells with evenly distributed adhesion receptors.
- The two product terms $1 + \eta_X^{XY} \langle \hat{z}, w \rangle$ and $1 - \eta_X^{XY} \langle \hat{z}, w \rangle$ are therefore, respectively, the adhesion potentials of the cells of type $X$ and the cells of type $Y$.

- The kernel $K(z)$ is a radial attraction kernel (meaning that it only depends on the norm of $z$) vanishing when $|z| > R$. For simplicity, in the numerical experiments, we take $K(z) = 1$ if $|z| \leq R$ and $K(z) = 0$ if $|z| > R$ but a smoother version should be considered for rigorous theoretical analysis.

#### 2.2 Repulsion forces

Let $K_\delta$ be an arbitrarily smooth radial kernel which vanishes at distance greater than $\delta > 0$. The repulsion exerted by a particle $j \in \{1, \ldots, N\}$ on a particle $i \in \{1, \ldots, N\}$ is defined by

$$F^{\mathrm{rep}}_{i \leftarrow j} := -h\big(K_\delta \star \hat{\mu}^N(x_i)\big) \nabla K_\delta(z_{ij}). \tag{2}$$

The meaning of each of the terms and symbols appearing in Eq. (2) is explained below.

- The empirical measure $\hat{\mu}^N := \frac{1}{N}\sum_{i=1}^N \delta_{x_i}$, also known as the counting measure, is an average of Dirac distributions at the cells' locations.

- The convolution product $\star$ is defined by

$$K_\delta \star \hat{\mu}^N(x) := \int_{\mathbb{R}^d} K_\delta(x-y)\hat{\mu}^N(\mathrm{d}y) = \frac{1}{N}\sum_{i=1}^N K_\delta(x - x_i).$$

  Note that if, let's say, $K_\delta$ is uniformly equal to 1 in a ball of radius $\delta$ and zero outside, then this quantity is simply the proportion of cells within the distance $\delta$ of the location $x$. We thus see this quantity as a measure of the local density of cells.

- The function $h : [0, +\infty) \to [0, +\infty)$ models the amount of repulsion depending on the local density, and can thus be understood as a pressure law. It is customary in the literature to consider power laws $h(\rho) = (\rho/\rho_0)^m$ for $m \geq 0$ or its degenerate version $h(\rho) = 0$ if $\rho \in [0, \rho_0)$ and $h(\rho) = \infty$ otherwise, where $\rho_0$ stands for the maximal acceptable local density [18].

- As the specific form of $K_\delta$ is not qualitatively important, in the simulation we consider a Morse-type repulsion kernel $K_\delta(r) = \exp(-5(r/\delta)^2)$. In the particle simulations, we consider a simple setting with $h$ constant equal to $F_0^{\mathrm{rep}} > 0$ so that in total

$$F^{\mathrm{rep}}_{i \leftarrow j} = -\frac{10F_0^{\mathrm{rep}}}{\delta^2}\exp\left(-5\left(\frac{|z_{ij}|}{\delta}\right)^2\right) z_{ij}.$$

  Keeping the full general form (2) will nevertheless be important for the coarse-graining procedure explained in the next section.

#### 2.3 Equations of motion

It is customary to describe the motion of biological cells at the microscopic scale in a high viscosity regime, leading to the overdamped Newton equations

$$\dot{x}_i = \frac{1}{N}\sum_{j\in X} F_{X\leftarrow X}(z_{ij}, w_{XX}) + \frac{1}{N}\sum_{j\in Y} F_{X\leftarrow Y}(z_{ij}, w_{XY}) + \frac{1}{N}\sum_{j=1}^{N} F^{\text{rep}}_{i\leftarrow j}, \tag{3}$$

for $i \in \{1, \dots, N\}$. This equation simply means that the time derivative of the position of the $i$-th cell, here assumed to be of type $X$, is the sum of all the binary interaction forces between pairs of cells. Note that, in accordance with the organoid experimental setting, we do not assume the existence of an Extra Cellular Matrix (ECM) that would give rise to active, non-binary, traction forces. We assume that the various quantities are properly adimensionalized and we renormalize each sum by $N$ to keep forces of order 1 regardless of the number of cells $N$. This latter scaling corresponds to a mean-field regime, as customary in statistical physics as it is the natural scaling to later take the large $N$ limit.

The system of $N$ equations (3) fully defines the cells' motion given a traction bias direction $w$. As explained in the main text, in the following the vector $w$ will be the gradient of the concentration of Wnt5a (up to some renormalization to make it of norm smaller than 1). The concentration of Wnt5a is itself coupled to the density of cells and to the concentration of Fgf8 through a reaction-diffusion network. To write this network, we first need to introduce a smoothen version of the cells's density, namely the functions

$$\phi_{\text{cells}}(x) := \frac{1}{N}\sum_{i=1}^{N} G_\sigma(x - x_i), \quad \phi_X(x) := \frac{1}{N_X}\sum_{i\in X} G_\sigma(x - x_i),$$

where $G_\sigma(z) = \frac{1}{(2\pi\sigma^2)^{d/2}} \exp\left(-\frac{|z|^2}{2\sigma^2}\right)$ is a Gaussian kernel with variance $\sigma^2$ (see Fig. 3). Then, the concentrations of Wnt5a and Fgf8, respectively denoted by $u$ and $v$ are given by the following reaction-diffusion network:

$$\partial_t u = -\lambda_u u + \mu_u h_{\text{in}}(\phi_D) + D_u \nabla \cdot (h_{\text{in}}(\phi_{\text{cells}})\nabla u), \tag{4a}$$

$$\partial_t v = -\lambda_v h_{\text{in}}(\phi_{\text{cells}}) v + \mu_v h_{\text{out}}(\phi_{\text{cells}})(1 - v) + D_v \nabla \cdot (h_{\text{out}}(\phi_{\text{cells}})\nabla v), \tag{4b}$$

supplemented by Dirichlet boundary conditions $u = 0$ and $v = 1$ at the boundary of the domain.

The meaning of each of the terms and symbols appearing in Eq. (4) is explained below.

- The nonnegative constants $\lambda_u, \lambda_v, \mu_u, \mu_v > 0$ are respectively the degradation rates and the production rates of Wnt5a and Fgf8.

- The nonnegative constants $D_u > 0$ and $D_v > 0$ are respectively the diffusion rates of Wnt5a and Fgf8.

- The functions $h_{\text{in}}$ and $h_{\text{out}}$ are convenience functions used to transform a field $\phi$ into smooth binary phase-fields bounded between two arbitrary values corresponding to $\phi$ small and large. Namely we consider

$$h_{\text{in}}(\phi) = h_- + (h_+ - h_-)\frac{(\phi/\phi_0)^2}{1 + (\phi/\phi_0)^2}$$

  and

$$h_{\text{out}}(\phi) = h_+ - (h_+ - h_-)\frac{(\phi/\varepsilon)^2}{1 + (\phi/\phi_0)^2},$$

  where $h_- < h_+$ are two arbitrary values, with typically $h_- = 0$ and $h_+ = 1$, and $\phi_0 > 0$ is a (small) thresholding parameter. To alleviate the notations here, we write only one set of parameters $(h_-, h_+, \phi_0)$ but we will consider different sets of parameter values for each of the terms where these functions appears (see Tables 1 and 2). An illustration of the purpose of these functions is shown (in dimension 1) in Fig. 3.

In summary, this reaction-diffusion models the following three rules:

1. Wnt5a is produced by distal cells.

2. Fgf8 is provided and saturated in the medium (with maximal concentration set to 1).

3. Wnt5a diffuses inside the cell aggregate while Fgf8 diffuses in the medium.

In practice we solve this reaction-diffusion system on a discrete uniform grid, with $M$ grid-cells per dimension.

Once $u$ and $v$ are defined, we will consider two polarization directions defined as

$$w_u := \frac{\nabla u}{g_0 + |\nabla u|}, \quad w_v = \frac{\nabla v}{g_0 + |\nabla v|} \tag{5}$$

where $g_0 > 0$ is another thresholding value.

### 3 Coarse-graining

The goal of this section is to mathematically derive a set of Partial Differential Equations (PDE) that describes the average behavior of the system in the limit of a large number of cells and on a large space-time scale.

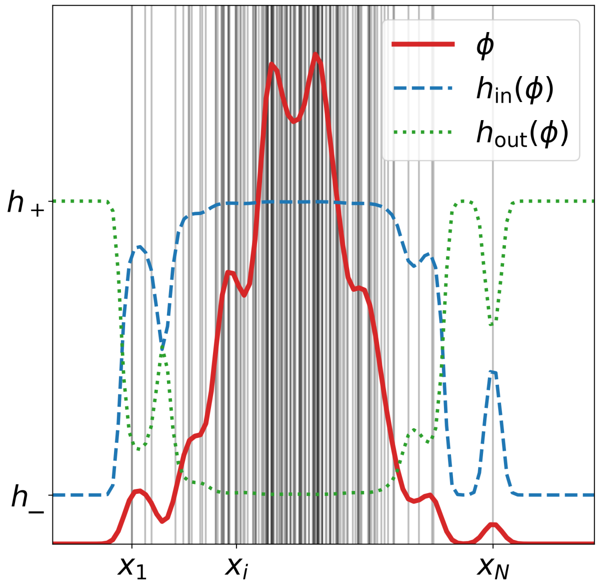


**Figure 3:** In dimension 1, the phase-field $\phi$ (red plain line) is obtained by convolving of a Gaussian kernel with Dirac masses at the positions $x_1, \ldots, x_N$ (gray vertical lines). This phase field is filtered by the convenience functions $h_{\text{in}}$ (blue dashed line) and $h_{\text{out}}$ (green dotted line) to bound it between two values $h_-$ and $h_+$.

#### 3.1 Mean-field limit

The first step is to consider the limit of a large number of cells $N \to +\infty$. In the equations of motion Eq. (3), the forces exerted on a particle can always be written as an average of binary elementary forces. This particular algebraic structure allows us to write an equivalent PDE system satisfied by the empirical measures

$$\hat{\mu}^D := \frac{1}{N}\sum_{i\in D}\delta_{x_i}, \quad \hat{\mu}^P := \frac{1}{N}\sum_{i\in P}\delta_{x_i}.$$

Namely, it can be checked that

$$\partial_t\hat{\mu}^D = -\nabla\cdot\Big(\hat{\mu}^D\big(\mathcal{F}_{D\leftarrow D}[\hat{\mu}^D] + \mathcal{F}_{D\leftarrow P}[\hat{\mu}^P] - h(K_\delta\star\hat{\mu}^N)\nabla K_\delta\star\hat{\mu}^N\big)\Big), \tag{6a}$$

$$\partial_t\hat{\mu}^P = -\nabla\cdot\Big(\hat{\mu}^P\big(\mathcal{F}_{P\leftarrow D}[\hat{\mu}^D] + \mathcal{F}_{P\leftarrow P}[\hat{\mu}^P] - h(K_\delta\star\hat{\mu}^N)\nabla K_\delta\star\hat{\mu}^N\big)\Big), \tag{6b}$$

where we recall that $\hat{\mu}^N = \frac{1}{N}\sum_{i=1}^N\delta_{x_i} = \hat{\mu}^D + \hat{\mu}^P$ and for an arbitrary measure $\rho$,

$$\mathcal{F}_{X\leftarrow Y}[\rho](x) := \xi_{XY}\int_{\mathbb{R}^d}\big(1+\eta_X^{XY}\langle\hat{z},w_{XY}\rangle\big)\big(1-\eta_Y^{XY}\langle\hat{z},w_{XY}\rangle\big)K(z)z\rho(x+z)\mathrm{d}z.$$

The system (6) is strictly equivalent to the equations of motion Eq. (3), it is often referred to as the Liouville equation. It should be understood in the so-called weak sense: by integration by parts on the right-hand side of the equations, it gives the time evolution of so-called observables of the form

$$\frac{1}{N}\sum_{i\in X}\varphi(x_i) = \int_{\mathbb{R}^d}\varphi(x)\hat{\mu}^X(x)\mathrm{d}x, \tag{7}$$

for any arbitrary (smooth) function $\varphi$.

The main advantage of the system (6) compared to the system of Eqs. (3) is that, despite the increased technical complexity, there are only two equations to consider instead of $N$, which makes it possible to compute the limit $N \to +\infty$. Namely, we want to prove the existence of a limit for the two empirical measures $\hat{\mu}^D$ and $\hat{\mu}^P$ and to be able to characterize it. Indeed, if such limiting measures did exist, it would allow us to compute the limit of any statistical quantity of the form (7). This problem is actually well understood and theorems can be readily applied to the present setting, even in a more general stochastic setting, see for example the review [6, 7]. Before writing this limit, we introduce another subtlety that will simplify the later analysis. We consider that as $N$ goes to infinity, the repulsion radius $\delta$ vanishes to zero (modeling the fact that repulsion interactions become purely localized in the large space-time regime considered). In that case and if $K_\delta$ is assumed to be smooth and of integral one, then for any $\rho$, $K_\delta \star \rho \to \rho$ and $\nabla K_\delta \star \rho \to \nabla\rho$ as $\delta \to 0$. By considering an appropriate scaling between $N$ and $\delta$ (often called moderate interaction in the literature [22, 20, 6, 7]), both limits can be taken at the same time. Finally, we obtain the following mean-field limit result: if the following limit is true at time $t = 0$ (which we assume),

$$(\hat{\mu}^D, \hat{\mu}^P) \underset{N\to+\infty}{\longrightarrow} (\rho^D, \rho^P),$$

then it remains valid at any further time $t > 0$ and the limiting functions are the solution of the system

$$\partial_t\rho^D = -\nabla\cdot\Big(\rho^D\big(\mathcal{F}_{D\leftarrow D}[\rho^D] + \mathcal{F}_{D\leftarrow P}[\rho^P] - \nabla H(\rho)\big)\Big), \tag{8a}$$

$$\partial_t\rho^P = -\nabla\cdot\Big(\rho^P\big(\mathcal{F}_{P\leftarrow D}[\rho^D] + \mathcal{F}_{P\leftarrow P}[\rho^P] - \nabla H(\rho)\big)\Big), \tag{8b}$$

where $H$ is an antiderivative of $h$ and $\rho = \rho^D + \rho^P$.

#### 3.2 Space-time rescaling

The kernel $K$ appearing in the attraction forces vanishes at a distance greater than the attraction radius of the cells, previously denoted by $R$. In the following we want to study cell aggregates that are much larger than this sensing radius (that is, we consider that the typical size of a cell is much smaller than the size of the aggregate). We follow the approach outline in [18]. Without loss of generality, in the following, we consider that $R = 1$ and we consider a large aggregate of total mass

$$\int_{\mathbb{R}^d}\rho(t,x)\mathrm{d}x = \varepsilon^{-d},$$

where the new variable $\varepsilon > 0$ plays the role of a small scaling parameter. We then define the new macrosocopic variables

$$\tilde{t} = \varepsilon^2 t, \quad \tilde{x} = \varepsilon x, \tag{9}$$

and the rescaled densities $\rho^{\varepsilon,D}(\tilde{t},\tilde{x}) = \rho^D(t,x)$, $\rho^{\varepsilon,P}(\tilde{t},\tilde{x}) = \rho^P(t,x)$ and $\rho^{\varepsilon}(\tilde{t},\tilde{x}) = \rho(t,x)$. After this change of variables, the total mass of $\rho^\varepsilon$ is normalized to 1 for any $\varepsilon > 0$ and the system (8) becomes, dropping the tildes in the space-time variables for simplicity,

$$\partial_t \rho^{\varepsilon,D} = -\varepsilon^{-1}\nabla\cdot\Big(\rho^{\varepsilon,D}\big(\mathcal{F}^{\varepsilon}_{D\leftarrow D}[\rho^{\varepsilon,D}] + \mathcal{F}^{\varepsilon}_{D\leftarrow P}[\rho^{\varepsilon,P}]\big)\Big) + \nabla\cdot\big(\rho^{\varepsilon,D}\nabla H(\rho^\varepsilon)\big), \tag{10a}$$

$$\partial_t \rho^{\varepsilon,P} = -\varepsilon^{-1}\nabla\cdot\Big(\rho^{\varepsilon,P}\big(\mathcal{F}^{\varepsilon}_{P\leftarrow D}[\rho^{\varepsilon,D}] + \mathcal{F}^{\varepsilon}_{P\leftarrow P}[\rho^{\varepsilon,P}]\big)\Big) + \nabla\cdot\big(\rho^{\varepsilon,P}\nabla H(\rho^\varepsilon)\big), \tag{10b}$$

where for an arbitrary function $\rho$,

$$\mathcal{F}^{\varepsilon}_{X\leftarrow Y}[\rho](x) := \xi_{XY}\int_{\mathbb{R}^d}\big(1+\eta_X^{XY}\langle\hat{z},w_{XY}\rangle\big)\big(1-\eta_Y^{XY}\langle\hat{z},w_{XY}\rangle\big)K(z)z\rho(x+\varepsilon z)\mathrm{d}z. \tag{11}$$

The system (8) is a nonlinear system of two coupled continuity equations on the densities of distal and proximal cells. The first term on the right-hand side of each equation accounts for the attraction forces between the cells. The terms of the form $\mathcal{F}^{\varepsilon}_{X\leftarrow Y}[\rho](x)$ are indeed an averaged version of the attraction forces (1) at a typical distance $\varepsilon$ of $x$ and due to cells distributed according to $\rho$, hence the denomination mean-field. The second term on the right-hand side of each equation accounts for the repulsion forces. Mathematically, it is often referred to as a nonlinear diffusion term, with an arbitrary pressure law $H$ that offers some modeling freedom. Choosing for instance $H(\rho) = \log\rho$ actually corresponds to a linear diffusion, as the corresponding term reduces to the Laplace operator $\Delta\rho$. The pressure law is typically rather a polynomial $H(\rho) = (\rho/\rho_0)^m$, with $m \geq 1$ and a critical density $\rho_0 > 0$. Such case corresponds to the so-called porous medium operator that has been largely studied in the past decades to model various advection and spreading phenomena such as gas flow, heat transfer or population dynamics, see for instance [29]. Compared to the linear diffusion operator, the case $m > 1$ often has more realistic properties, in particular propagation at finite time that naturally lead to the study of free boundary problems. It comes at the price of increased mathematical complexity, due to the nonlinear structure, that we will not investigate further here.

### 4 From nonlocal to local

Another major difficulty in the analysis of the system (10) is its nonlocal nature, meaning that the solutions $(\rho^{\varepsilon,D},\rho^{\varepsilon,P})$ appear in integrated form in the term (11). In the following we compute a formal approximation of this nonlocal term in the limit $\varepsilon \to 0$

by using a Taylor expansion of the solutions. Note that the procedure is somehow known for aggregation-diffusion equations that correspond to a simple version of our setting, without traction bias (that is when $\eta_{XX} = 0$) and mostly for one population. In this case, the localized version of the system (10) is the Cahn-Hilliard equation, see the following section as well as [3, 11, 18] and the references therein. Rigorous convergence results and analysis are shown for instance in [18, 10]. On a formal point of view, the system (10) that we derived from the ABM brings two novelties to this framework.

1. We consider two populations with differential adhesion. Hence our model contains a model for cell sorting which is an important topic in developmental biology and in mathematical modeling. Although it is not the main point of the present article, we introduce in the following a new two-population version of the Cahn-Hilliard equation that reproduces classical cell sorting experiments, and in particular that can be compared with the continuum models introduced in [5]. This intermediary step demonstrates the relevance of the approach before introducing the traction bias hypothesis and its consequences.

2. The traction bias hypothesis introduces a novel modification of the Cahn-Hilliard equation, which formally will take the form of an additional nonlinear anisotropic diffusion term (that is, diffusion biased in the direction denoted by $w$). Importantly, this additional term is sufficient to make a spherical aggregate unstable and to produce fingering like patterns.

#### 4.1 Taylor expansion of the mean-field force

There are four terms that appear in the system (10) and that share the same algebraic structure, we thus write a Taylor expansion in the abstract form (11) before assembling the various terms together. To alleviate the notations we consider the case $\eta_X^{XY} \equiv \eta_X$ in the following. Then Eq. (11) can be rewritten

$$\mathcal{F}^{\varepsilon}_{X\leftarrow Y}[\rho](x) = \xi_{XY} \int_{\mathbb{R}^d} \Big(1 - \eta_X\eta_Y \langle \hat{z}, w_{XY}\rangle^2 + (\eta_X - \eta_Y)\langle \hat{z}, w_{XY}\rangle\Big) K(z) z \rho(x+\varepsilon z)\mathrm{d}z,$$

and the Taylor expansion of $\rho$ around $x$ is

$$\rho(x+\varepsilon z) = \rho(x) + \varepsilon\langle \nabla\rho(x), z\rangle + \varepsilon^2 \langle H_\rho(x) z, z\rangle + \varepsilon^3 D^3\rho(x)(z,z,z) + \mathcal{O}(\varepsilon^4),$$

where $H_\rho(x)$ denotes the Hessian matrix of $\rho$ at $x$ and $D^3\rho(x)$ its third derivative, seen as a trilinear operator. Inserting this expansion in the expression above, we can write

$$\mathcal{F}^{\varepsilon}_{X\leftarrow Y} = \xi_{XY} \sum_{k=0}^{3} \varepsilon^k f^k_{X\leftarrow Y} + \mathcal{O}(\varepsilon^4),$$

where each of the terms $f^k_{X\leftarrow Y}$ involves the $k$-th derivative of $\rho$ at $x$ multiplied by a factor that is the integral of a polynomial in $z$ times $K(z)$. As the latter is a symmetric radial kernel, all odd monomials will automatically vanish (as it can be seen for instance with the change of variable $z \mapsto -z$). This leads to the following splitting for the first four orders in $\varepsilon$.

##### 4.1.1 Order 0

Since, due to the radial symmetry of $K$,

$$\int_{\mathbb{R}^d} (1 - \eta_X \eta_Y \langle \hat{z}, w_{XY} \rangle^2) K(z) z \mathrm{d}z = 0,$$

only the cross term between $X$ and $Y$ remains:

$$f^0_{X \leftarrow Y} = (\eta_X - \eta_Y) \left( \int_{\mathbb{R}^d} K(z) \langle \hat{z}, w_{XY} \rangle z \, \mathrm{d}z \right) \rho(x).$$

We can be more precise about the integral factor. Using the spherical change of variable $z = ru$ with $r \in [0, +\infty)$ and $u \in \mathbb{S}^{d-1}$ (the $(d-1)$-dimensional sphere), it holds that

$$\int_{\mathbb{R}^d} K(z) \langle \hat{z}, w_{XY} \rangle z \, \mathrm{d}z = m_d \left( \int_{\mathbb{S}^{d-1}} u \otimes u \, \mathrm{d}u \right) w_{XY},$$

where we introduce the moments

$$m_k := \int_0^{+\infty} K(r) r^k \, \mathrm{d}r,$$

and where $u \otimes u$ denotes the tensor product of two vectors, which, in components, is the matrix $(u_i u_j)_{i,j=1,\ldots,d}$ when $u = (u_1, \ldots, u_d)^{\mathrm{T}}$. Using the fact that $|u| = 1$ and the invariance by rotation, it can be shown that

$$\int_{\mathbb{S}^{d-1}} u \otimes u \, \mathrm{d}u = \frac{1}{d} \mathrm{Id},$$

and therefore

$$f^0_{X \leftarrow Y} = \frac{m_d}{d} (\eta_X - \eta_Y) \rho(x) w_{XY}.$$

##### 4.1.2 Order 1

Using the same arguments, namely the radial symmetry of $K$ and a spherical change of variable, we obtain

$$f^1_{X \leftarrow Y} = m_{d+1} \Big( d^{-1} \mathrm{Id} - \eta_X \eta_Y D(w_{XY}) \Big) \nabla \rho(x),$$

with the matrix

$$D(w_{XY}) := \int_{\mathbb{S}^{d-1}} \langle u, w_{XY} \rangle^2 u \otimes u \, \mathrm{d}u.$$

Again, this matrix can be written more explicitly using the integration formula for polynomials on the sphere that is shown for instance in [12]. As we will use this formula a lot in the following, we reproduce it below for convenience. Let $P(x) = x_1^{\alpha_1} \dots x_d^{\alpha_d}$ be a monomial in $d$ dimensions, with $x = (x_1, \dots, x_d)$ and $\alpha_1, \dots, \alpha_d \geq 0$. Then,

$$\int_{\mathbb{S}^{d-1}} P(x) \mathrm{d}\sigma(x) = \frac{2\Gamma(\beta_1)\Gamma(\beta_2)\dots\Gamma(\beta_d)}{\Gamma(\beta_1 + \beta_2 + \dots \beta_d)}, \tag{12}$$

if all the coefficients $\alpha_i$ are even and the integral is equal to zero otherwise. In this expression, $\mathrm{d}\sigma$ denotes the surface measure of the sphere, $\Gamma$ is the Gamma function and $\beta_i = \frac{1}{2}(\alpha_i + 1)$ for $i = 1, \dots, d$.

In the present case, we obtain that

$$D(w_{XY}) = \beta \left( \frac{2}{3} w_{XY} \otimes w_{XY} + \frac{1}{3} \mathrm{Id} \right),$$

where

$$\beta = \int_{\mathbb{S}^{d-1}} u_1^4 \, \mathrm{d}u = \frac{3}{d(d+2)}.$$

To prove this, let $w^\perp, \tilde{w}^\perp$ be unit vectors orthogonal to $w_{XY}$ and to each other when the dimension allows (that is when $d \geq 3$). Then

$$\begin{aligned}
\langle w_{XY}, D(w_{XY}) w_{XY} \rangle &= \beta \\
\langle w^\perp, D(w_{XY}), w_{XY} \rangle &= \langle w_{XY}, D(w_{XY}) w^\perp \rangle = \int_{\mathbb{S}^{d-1}} u_1^3 u_2 \, \mathrm{d}u = 0 \\
\langle w^\perp, D(w_{XY}) w^\perp \rangle &= \int_{\mathbb{S}^{d-1}} u_1^2 u_2^2 \, \mathrm{d}u =: \alpha \\
\langle \tilde{w}^\perp, D(w_{XY}) w^\perp \rangle &= \int_{\mathbb{S}^{d-1}} u_1^2 u_2 u_3 \, \mathrm{d}u = 0.
\end{aligned}$$

It implies that

$$D(w_{XY}) = \beta w_{XY} \otimes w_{XY} + \alpha(\mathrm{Id} - w_{XY} \otimes w_{XY}).$$

The conclusion follows by noting that $1 = d\beta + d(d-1)\alpha$ and using the relation $\beta = 3\alpha$ for any $d$ which can be proved by applying the formula 12.

###### 4.1.3 Order 2

Using the same arguments, we obtain

$$f^2_{X \leftarrow Y} = (\eta_X - \eta_Y) m^{d+2} \int_{\mathbb{S}^{d-1}} \langle u, w_{XY} \rangle \langle u, H_\rho(x) u \rangle u \, \mathrm{d}u.$$

As before, this leads to the computation of a polynomial on the sphere with linear coefficients in $w_{XY}$ and the $k$-th derivative of $\rho$. To write it more explicitly, it will be useful to write it in components. Writing $v_k$ the $k$-th component of a vector $v$ and

$\partial^2_{ij} \equiv \partial^2/\partial x_i \partial x_j$ the partial second derivative along the $i$ and $j$ components, the $\ell$-th component of the integral term is equal to

$$\int_{\mathbb{S}^{d-1}} \langle u, w_{XY}\rangle \langle u, H_\rho(x) u\rangle u_\ell \, \mathrm{d}u = \sum_{i,j,k=1}^{d} (w_{XY})_k \partial^2_{ij}\rho(x) \int_{\mathbb{S}^{d-1}} u_i u_j u_k u_\ell \, \mathrm{d}u.$$

By symmetry, the last integral on the right-hand side is equal to $\beta$ if all the indices are equal, to $\alpha$ if the indices are equal two by two (e.g. $i = j$ and $k = \ell$ but $i \neq k$) and zero otherwise, where we recall that $3\alpha = \beta = \frac{3}{d(d+2)}$. Thus, more generally, given arbitrary coefficients $(a_{ijk})_{i,j,k}$ it holds that

$$\begin{aligned}\sum_{i,j,k} a_{ijk} \int_{\mathbb{S}^{d-1}} u_i u_j u_k u_\ell \, \mathrm{d}u &= \alpha \left( \sum_{k\neq\ell} a_{\ell kk} + \sum_{k\neq\ell} a_{k\ell k} + \sum_{k\neq\ell} a_{kk\ell} \right) + \beta a_{\ell\ell\ell} \\ &= \alpha \left( \sum_{k} a_{\ell kk} + \sum_{k} a_{k\ell k} + \sum_{k} a_{kk\ell} \right) \qquad (13)\end{aligned}$$

where the last line comes from the fact that $\beta = 3\alpha$. Applying this formula to $a_{ijk} = (w_{XY})_k \partial^2_{ij}\rho(x)$, this leads after simplifications to

$$f^2_{X\leftarrow Y} = (\eta_X - \eta_Y) \frac{3m_{d+2}}{d(d+2)} \left( \frac{1}{3}\Delta\rho(x)\mathrm{Id} + \frac{2}{3} H_\rho(x) \right) w_{XY}.$$

###### 4.1.4 Order 3

Using the same arguments as before,

$$f^3_{X\leftarrow Y} = m_{d+3} \int_{\mathbb{S}^{d-1}} \left(1 - \eta_X \eta_Y \langle u, w_{XY}\rangle^2\right) D^3\rho(x)(u,u,u) u \, \mathrm{d}u$$

For the $\ell$-th component, using the formula (13) with $a_{ijk} = \frac{\partial^3 \rho}{\partial x_i \partial x_j \partial x_k}$, it holds that

$$\begin{aligned}\int_{\mathbb{S}^{d-1}} D^3\rho(x)(u,u,u) u_\ell \, \mathrm{d}z &= \sum_{i,j,k} \int_{\mathbb{S}^{d-1}} u_i u_j u_k u_\ell \, \mathrm{d}u \, \frac{\partial^3 \rho}{\partial x_i \partial x_j \partial x_k} \\ &= \beta \sum_k \frac{\partial^3 \rho}{\partial x_\ell \partial x_k^2} = \beta \frac{\partial}{\partial x_\ell} \Delta\rho(x),\end{aligned}$$

for the constant $\beta = \frac{3}{d(d+2)}$. The second term is more involved since we have to integrate a polynomial of order 6.

$$\int_{\mathbb{S}^{d-1}} \langle u, w_{XY}\rangle^2 D^3\rho(x)(u,u,u)u_\ell \,\mathrm{d}u = \sum_{i,j,k} \int_{\mathbb{S}^{d-1}} \langle u, w_{XY}\rangle^2 u_i u_j u_k u_\ell \,\mathrm{d}u \frac{\partial^3 \rho}{\partial x_i \partial x_j \partial x_k}$$
$$= 3\sum_j \int_{\mathbb{S}^{d-1}} \langle u, w_{XY}\rangle^2 u_\ell^2 u_j^2 \,\mathrm{d}u \,\frac{\partial^3 \rho}{\partial x_\ell \partial x_j^2}$$
$$= 3\sum_{k,j} (w_{XY})_k^2 \int_{\mathbb{S}^{d-1}} u_k^2 u_j^2 u_\ell^2 \,\mathrm{d}u \,\frac{\partial^3 \rho}{\partial x_\ell \partial x_j^2}$$
$$=: 3\sum_{j=1}^d M_{\ell,j}(w_{XY}) \frac{\partial^3 \rho}{\partial x_\ell \partial x_j^2}$$

where at the second line we have used the fact that one of the indices $i, j, k$ must be equal to $\ell$ and the two others must be equal together. At the third line, we expand the dot product and only retain the square monomials as all the other cross products lead to vanishing odd monomials. Finally, at the last line, we have introduced the matrix $M(w) := (M_{ij}(w))_{i,j}$ with components

$$M_{ij}(w) = \int_{\mathbb{S}^{d-1}} \langle u, w\rangle^2 u_i^2 u_j^2 \,\mathrm{d}u = \sum_k w_k^2 \int_{\mathbb{S}^{d-1}} u_i^2 u_j^2 u_k^2 \,\mathrm{d}u.$$

The diagonal terms are equal to

$$M_{ii}(w) = \gamma_2 |w|^2 + (\gamma_3 - \gamma_2) w_i^2$$

and the off-diagonal terms are equal to

$$M_{ij}(w) = \gamma_1 |w|^2 + (\gamma_2 - \gamma_1)(w_i^2 + w_j^2), \quad i \neq j$$

where we have introduced the coefficients

$$\gamma_1 = \int_{\mathbb{S}^{d-1}} u_1^2 u_2^2 u_3^2 \,\mathrm{d}u, \quad \gamma_2 = \int_{\mathbb{S}^{d-1}} u_1^4 u_2^2 \,\mathrm{d}u, \quad \gamma_3 = \int_{\mathbb{S}^{d-1}} u_1^6 \,\mathrm{d}u.$$

Note that using 12 again, one can compute $\gamma_2 = \frac{\beta}{d+4} = \frac{3}{d(d+2)(d+4)}$ and check that $\gamma_3 = 5\gamma_2 = 25\gamma_1$. However, note that $\gamma_1$ only exists when $d > 2$. Hence, for any $w \in \mathbb{R}^d$

$$\frac{1}{3}\int_{\mathbb{S}^{d-1}} \langle u, w_{XY}\rangle^2 D^3\rho(x)(u,u,u)u_\ell \,\mathrm{d}u$$
$$= \left(\gamma_2 |w_{XY}|^2 + (\gamma_3 - \gamma_2)(w_\ell^2)\frac{\partial^3 \rho}{\partial x_\ell^3} + \gamma_1 |w|^2 \sum_{j\neq \ell} \frac{\partial^3 \rho}{\partial x_\ell \partial x_j^2} + (\gamma_2 - \gamma_1)\sum_{j \neq \ell} (w_\ell^2 + w_j^2)\frac{\partial^3 \rho}{\partial x_\ell \partial x_j^2}\right.$$
$$= \gamma_2 |w|^2 \frac{\partial}{\partial x_\ell}\Delta\rho + (\gamma_3 - \gamma_2) w_\ell^2 \frac{\partial^3 \rho}{\partial x_\ell^3} + (\gamma_1 - \gamma_2)\sum_{j \neq \ell}\sum_{i \neq i,\ell} w_i^2 \frac{\partial^3 \rho}{\partial x_\ell \partial x_j^2}$$

Gathering everything finally leads to

$$f^3_{X\leftarrow Y} = \frac{3m_{d+3}}{d(d+2)}\left(\left(1-\eta_X\eta_Y\frac{3|w_{XY}|^2}{d+4}\right)\nabla\Delta\rho \right.$$
$$\left. -\eta_X\eta_Y\frac{12}{d+4}R_3(w_{XY},\rho)+\eta_X\eta_Y\frac{12}{5(d+4)}\mathbb{1}_{d>2}\tilde{R}_3(w_{XY},\rho)\right),$$

where for any $w \in \mathbb{R}^d$, $R_3(w,\rho)$ and $\tilde{R}_3(w,\rho)$ are $d$-dimensional vectors the $\ell$-th components of which are respectively equal to

$$R_3(w,\rho)_\ell = w_\ell^2\frac{\partial^3\rho}{\partial x_\ell^3},\quad \tilde{R}_3(w,\rho)_\ell = \sum_{j\neq\ell}\sum_{i\neq i,\ell} w_i^2\frac{\partial^3\rho}{\partial x_\ell \partial x_j^2}$$

#### 4.2 Summary and conclusion

We recall that our goal is to approximate at order $\varepsilon^3$ the system (10) that we recall below for convenience:

$$\partial_t\rho^{\varepsilon,D} = -\varepsilon^{-1}\nabla\cdot\left(\rho^{\varepsilon,D}\big(\mathcal{F}^\varepsilon_{D\leftarrow D}[\rho^{\varepsilon,D}]+\mathcal{F}^\varepsilon_{D\leftarrow P}[\rho^{\varepsilon,P}]\big)\right)+\nabla\cdot\big(\rho^{\varepsilon,D}\nabla H(\rho^\varepsilon)\big),$$
$$\partial_t\rho^{\varepsilon,P} = -\varepsilon^{-1}\nabla\cdot\left(\rho^{\varepsilon,P}\big(\mathcal{F}^\varepsilon_{P\leftarrow D}[\rho^{\varepsilon,D}]+\mathcal{F}^\varepsilon_{P\leftarrow P}[\rho^{\varepsilon,P}]\big)\right)+\nabla\cdot\big(\rho^{\varepsilon,P}\nabla H(\rho^\varepsilon)\big),$$

by inserting the the Taylor expansion

$$\mathcal{F}^\varepsilon_{X\leftarrow Y} = \xi_{XY}\sum_{k=0}^{3}\varepsilon^k f^k_{X\leftarrow Y}+\mathcal{O}(\varepsilon^4),$$

and we have computed the coefficients:

$$f^0_{X\leftarrow Y} = \frac{m_d}{d}(\eta_X-\eta_Y)\rho(x)w_{XY},$$
$$f^1_{X\leftarrow Y} = m_{d+1}\Big(d^{-1}\mathrm{Id}-\eta_X\eta_Y D(w_{XY})\Big)\nabla\rho(x),$$
$$f^2_{X\leftarrow Y} = (\eta_X-\eta_Y)\frac{3m_{d+2}}{d(d+2)}\left(\frac{1}{3}\Delta\rho(x)\mathrm{Id}+\frac{2}{3}H_\rho(x)\right)w_{XY},$$
$$f^3_{X\leftarrow Y} = \frac{3m_{d+3}}{d(d+2)}\left(\left(1-\eta_X\eta_Y\frac{3|w_{XY}|^2}{d+4}\right)\nabla\Delta\rho \right.$$
$$\left. -\eta_X\eta_Y\frac{12}{d+4}R_3(w_{XY},\rho)+\eta_X\eta_Y\frac{12}{5(d+4)}\mathbb{1}_{d>2}\tilde{R}_3(w_{XY},\rho)\right).$$

In the following we will consider the case

$$\eta\equiv\eta_D>0,\quad \eta_P=0,\quad w_{DD}=w_u,\quad w_{DP}=w_v.$$

In plain words, this modeling choice assumes that only the distal cells are subject to a traction bias which follows the gradient of Wnt5a in the case of distal-distal interactions and the gradient of Fgf8 in the case of distal-proximal interactions. For notational convenience, in the following we write $w_u \equiv w$ and $w_v \equiv f$. The morphogens concentrations $u$ and $v$ are given by a reaction-diffusion network analogous to (4) that will be explained below. Reporting this leads to the final system:

$$\partial_t \rho_D + \varepsilon^{-1} \xi_{DP} \eta \nabla \cdot \big(\rho_D A^\varepsilon(\rho_P) f\big) = c_1 \nabla \cdot (\rho_D \nabla q_D^\varepsilon) + \eta^2 \xi_{DD} \nabla \cdot \big(\rho_D [c_2 D(w) \nabla \rho_D + \varepsilon^2 \hat{R}_3(w,\rho)]\big) \tag{14a}$$

$$\partial_t \rho_P - \varepsilon^{-1} \xi_{DP} \eta \nabla \cdot \big(\rho_P A^\varepsilon(\rho_D) f\big) = c_1 \nabla \cdot (\rho_P \nabla q_P^\varepsilon) \tag{14b}$$

where we recall that $\rho = \rho_D + \rho_P$ and we have defined the potential functions

$$q_D^\varepsilon := W'(\rho) - \varepsilon^2 \xi_{DP} c_3 \Delta\rho - (\xi_{DD} - \xi_{DP})(c_4 \rho_D + \varepsilon^2 c_3 \Delta \rho_D) + \varepsilon^2 \eta^2 \xi_{DD} \tilde{c}_3 \Delta \rho_D$$
$$q_P^\varepsilon := W'(\rho) - \varepsilon^2 \xi_{DP} c_3 \Delta\rho - (\xi_{PP} - \xi_{DP})(c_4 \rho_P + \varepsilon^2 c_3 \Delta \rho_P)$$

and the following quantities

**(Advection matrix)** $A^\varepsilon(\rho) := c_0 \rho \mathrm{Id} + \varepsilon^2 c_5 (\Delta \rho \mathrm{Id} + 2 H_\rho)$,

**(Anisotropic diffusion matrix)** $D(w) := \frac{2}{3} w \otimes w + \frac{1}{3} \mathrm{Id}$,

**(Interaction potential)** $W'(\rho) := H(\rho) - \xi_{DP} \frac{m_{d+1}}{d} \rho$,

**(Third-order correction)** $\hat{R}_3(w,\rho) := c_6 \left( R_3(w,\rho) - \frac{1}{5} \mathbb{1}_{d>2} \tilde{R}_3(w,\rho) \right)$.

All the coefficients $c_0, \dots, c_6 > 0$ have an explicit expression, they only depend on the dimension and the moments of $K$ (that can be defined arbitrarily), namely

$$c_0 = \frac{m_d}{d}, \quad c_1 = 1, \quad c_2 = \frac{3 m_{d+1}}{d(d+2)}, \quad c_3 = \frac{3 m_{d+3}}{d(d+2)}, \quad c_4 = \frac{m_{d+1}}{d}$$

$$\tilde{c}_3 = \frac{9 m_{d+3} |w|^2}{d(d+2)(d+4)}, \quad c_5 = \frac{m_{d+2}}{d(d+2)}, \quad c_6 = \frac{36 m_{d+3}}{d(d+2)(d+4)}$$

### 5 Discussion and simplified system

Despite its slightly convoluted form, the full system (14) is nothing but a system of continuity equations (that is a system of equations of the form $\partial_t \rho + \nabla \cdot (\rho J) = 0$) that reflects the conservation of the number of cells. The important quantity to characterize in this situation is the so-called flux $J$. In the present case, it can be split into three terms.

1. The dominating term of order $\varepsilon^{-1}$ is a pure advection term in the direction $A^\varepsilon(\rho_P)w$ for the distal cells and in the direction $-A^\varepsilon(\rho_D)w$ for the proximal cells. Note that neglecting the $\varepsilon^2$ correction in the definition of $A^\varepsilon$, we simply obtain that the two cell populations move in the opposite directions $w$ for the distal cells and $-w$ for the proximal cells, as long as $\rho_D\rho_P > 0$ and $\eta > 0$ do not vanish. This induces a segregation phenomenon along the direction $w$ under the traction bias hypothesis.

2. The first term on the right-hand side of each equation encodes the attraction-repulsion interactions into the potentials $q_D^\varepsilon$ and $q_P^\varepsilon$. These potentials generalize the well-known Cahn-Hilliard potential to the case of two populations. The Cahn-Hilliard potential is classically written

$$q^\varepsilon = W'(\rho) - \varepsilon^2\Delta\rho$$

in the case of a single population with density $\rho$. As explained in [18], when the repulsion term $H$ is a power law, the interaction potential $W$ is typically a bistable potential with two stable wells, here by convention, at 0 and $\rho_0 > 0$ corresponding respectively to empty zones and to zones with a maximal density above which repulsion will dominate. In the single population case, this term is responsible for the phase separation between two domains $\{\rho = 0\}$ and $\{\rho = \rho_0\}$. In the present case where two populations are considered, the correction terms with prefactor $c_4$ reflect the potentially different adhesion strengths between each pair of cells which might modify the maximal density for each cell type. Naturally, this term vanishes if $\xi_{DP} = \xi_{DD} = \xi_{PP}$ as the two-population model reduces to a one-population model in that case. It is important to note that, due to the sign of $W'$, removing the $\varepsilon^2$ term in $q_D^\varepsilon, q_P^\varepsilon$ would lead to an ill-posed forward-backward diffusion term, which is the reason why an expansion up to the order 3 was needed. With this term, the interface between the phases is typically of size $\varepsilon^2$. The term $\tilde{c}_3$ (for the distal cells only) has a different nature and comes from the traction bias.

3. Finally, the main novelty in this system is the last term on the right-hand side of the equation on $\rho_D$. This term is a consequence of the traction bias hypothesis (in the sense that it vanishes when $\eta = 0$). The present analysis reveals that at order 1, the main consequence of this hypothesis is an additional anisotropic diffusion term, with diffusion matrix $D(w)$. Additional third-order correction terms should be considered at order $\varepsilon^2$ but their convoluted form makes them difficult to interpret directly. We will neglect them in the following as they do not seem to be involved in the fingering phenomenon that we are interested in.

In the following, we will consider a simplified system that preserves the structural properties of the full system (14) but which will be simpler to simulate numerically while retaining a similar expected behavior. First of all we will neglect the $\varepsilon^2$ term in the advection matrix as well as the third order correction terms. These small order terms are unlikely to change drastically the advection-diffusion directions and do not seem to play a role for the well-posedness of the system. On the contrary, as explained before, we do need to keep the $\varepsilon^2$ terms in the Cahn-Hilliard potentials, except the one corresponding to the term $\tilde{c}_3$ that is rather similar to the third-order corrections. Moreover, although we have shown that all the terms can be directly related to the parameters of the agent-based model, we rather seek a qualitative agreement here so for convenience and to keep more modeling freedom, we will consider the coefficients $c_0, c_1, c_2, c_3, c_4 > 0$ as free parameters (and $c_5 = c_6 = \tilde{c}_3 = 0$). Each one has a specific physical meaning that is summarized in the Table 3 below. Finally, as a scaling choice, we also fix the thresholding density $\rho_0 = 1$ by considering a function $H$ such that

$$W(\rho) = \lambda\rho^2(\rho_0 - \rho)^2,$$

for some arbitrary $\lambda > 0$ that account for the strength of the phase separation (see [18] for how to do this). Finally, we add a (small) linear diffusion term to each equation, here mostly for numerical stability reasons but note that linear diffusion can also be rigorously derived from the agent-based model with the classical assumption that Brownian noise is added to the position of the particles. These simplifications lead to the following system that will be considered in the remaining of the article

$$\partial_t\rho_D + \varepsilon^{-1}\eta\xi_{DP}c_0\nabla\cdot(\rho_D\rho_P f) = c_1\nabla\cdot(\rho_D\nabla q_D^\varepsilon) + \eta^2\xi_{DD}c_2\nabla\cdot\big(\rho_D D(w)\nabla\rho_D\big), \quad (15a)$$

$$\partial_t\rho_P - \varepsilon^{-1}\eta\xi_{DP}c_0\nabla\cdot(\rho_P\rho_D f) = c_1\nabla\cdot(\rho_P\nabla q_P^\varepsilon), \quad (15b)$$

$$q_D^\varepsilon = W'(\rho) - \varepsilon^2\xi_{DP}c_3\Delta\rho - (\xi_{DD} - \xi_{DP})(c_4\rho_D + \varepsilon^2 c_3\Delta\rho_D) + \nu\log\rho_D, \quad (15c)$$

$$q_P^\varepsilon = W'(\rho) - \varepsilon^2\xi_{DP}c_3\Delta\rho - (\xi_{PP} - \xi_{DP})(c_4\rho_P + \varepsilon^2 c_3\Delta\rho_P) + \nu\log\rho_P. \quad (15d)$$

where we recall that $\rho = \rho_D + \rho_P$ and we also added linear diffusion with coefficient $\nu$ to the system as an additional modelling tool. Finally, the traction bias directions are given, as in the agent-based model, by

$$w = \frac{\nabla u}{g_0 + |\nabla u|}, \quad f = \frac{\nabla v}{g_0 + |\nabla v|},$$

where $u$ is the Wnt5a concentration that is coupled to the Fgf8 concentration $v$ and the cell density through the following reaction-diffusion network

$$\partial_t u = -\lambda_u u + \mu_u\rho_D + D_u\nabla\cdot(\rho\nabla u), \quad (16a)$$

$$\partial_t v = -\lambda_v\rho v + \mu_v(1-\rho)(1-v) + D_v\nabla\cdot((1-\rho)\nabla v). \quad (16b)$$

This latter system is identical to the previous one (4) up to replacing $\phi_D$ and $\phi_{\text{cells}}$ by $\rho_D$ and $\rho$ and to using the simplest functions $h_{\text{in}}(\rho) = \rho$ and $h_{\text{out}}(\rho) = 1 - \rho$ since the potential $W$ has two stable wells in 0 and 1.

The parameters, their meaning and numerical values are summarized in the Table 3.

### 6 Two populations without traction bias: cell-sorting experiments

In order to link the continuum model (15) with the existing models in the literature, we first consider the case $\eta = 0$. In that situation, the model reduces to the following two-population aggregation-diffusion model where the main parameters are the (relative) adhesion strength $\xi_{DD}$, $\xi_{PP}$ and $\xi_{DP}$.

$$\partial_t \rho_D = c_1 \nabla \cdot (\rho_D \nabla q_D^\varepsilon), \tag{17a}$$

$$\partial_t \rho_P = c_1 \nabla \cdot (\rho_P \nabla q_P^\varepsilon), \tag{17b}$$

$$q_D^\varepsilon = W'(\rho) - \varepsilon^2 \xi_{DP} c_3 \Delta\rho - (\xi_{DD} - \xi_{DP})(c_4 \rho_D + \varepsilon^2 c_3 \Delta \rho_D) + \nu \log \rho_D, \tag{17c}$$

$$q_P^\varepsilon = W'(\rho) - \varepsilon^2 \xi_{DP} c_3 \Delta\rho - (\xi_{PP} - \xi_{DP})(c_4 \rho_P + \varepsilon^2 c_3 \Delta \rho_P) + \nu \log \rho_P. \tag{17d}$$

As already mentioned earlier, this model is similar to other models in the literature that rather focus on cell sorting phenomena. Although it is not the main goal of the present article, as a sanity check and for the sake of completeness we briefly review a few milestones below and show that the reduced model (17) generalizes some of them and can overall achieve comparable results.

Since the seminal article [2], several continuum (PDE) models have been proposed to model the cell-cell adhesion phenomena that lead to striking cell sorting patterns, as originally shown experimentally by Steinberg [26, 26, 27]. As in the present article, such models are often non-local models since they provide a lot of modeling freedom to tune attraction-repulsion interactions (with e.g. population pressure and density saturation) and they naturally appear as the scaling limit of agent-based models [2, 5]. The derivation of local continuum model through a procedure similar to the one presented above has been carried out in [2, 3] for one population and leads to Cahn-Hilliard type equations. In the mathematical physics literature, these models are also often related to the family of thin-film equations. Although the extension to two populations does not represent a major difficulty, a local two-population model has been introduced only recently in [11]. This latter model corresponds to a special case of our model (17) with a quadratic kernel such that $W'(\rho) \propto \rho$.

A major question for all the aforementioned models is the stability of the constant solution (that is of the uniform and stationary state) that can be studied using a so-called linear stability analysis. More precisely, the goal is to consider a small perturbation of a constant solution $(\rho_D^*, \rho_P^*)$:

$$\rho_D^\delta = \rho_D^* + \delta \rho_D \mathrm{e}^{ik\cdot x + \sigma(k)t}, \quad \rho_P^\delta = \rho_P^* + \delta \rho_D \mathrm{e}^{ik\cdot x + \sigma(k)t},$$

for $\delta > 0$ small and where $k \in \mathbb{R}^d$ is a wave number which is unstable as soon as the coefficient $\sigma(k)$ as a nonnegative real part. To assess whether such unstable mode can exist, we first insert this expression into the equation (17), then an expansion at order 1 when $\delta \to 0$ classically leads to a dispersion relation linking $\sigma(k)$ and $k$. In the case of a two-population model, $\sigma(k)$ is equal to the eigenvalues of the $2 \times 2$ matrix corresponding to the linearization of the right-hand side of (17) evaluated at $(\rho_D^*, \rho_P^*)$. After classical and elementary computations, this leads to the following sufficient condition for a constant solution $\rho_D^* = \rho_P^*$ to be unstable is

$$\frac{\xi_{DD} + \xi_{PP}}{2}(\varepsilon|k|^2 - 1) + \xi_{DP} + W''(\rho^*) + \frac{4\nu}{\rho^*} < 0.$$

Thus, regardless of the choice of $W$, for $\varepsilon$ small enough and $\xi_{DP}$ given, unstable modes exist as soon as the average $\frac{\xi_{DD}+\xi_{PP}}{2}$ is large enough, that is, as soon as the adhesion strength among one population is strong enough. In the case $\xi_{DD} = \xi_{PP} = \xi_{DP} \equiv \xi$ and $\nu = 0$, the model reduces to a single population and we recover a classical instability condition for the Cahn-Hilliard equation

$$\xi\varepsilon|k|^2 + W''(\rho^*) < 0,$$

in particular unstable modes can exist only if $W''(\rho^*) < 0$ corresponding to the purely attractive case considered e.g. in [11].

The analysis predicts that it is possible to make a stationary state unstable by an appropriate tuning of the adhesion coefficients $\xi_{DD}, \xi_{DP}, \xi_{PP}$. We confirm this finding in Fig. 4 by reproducing the three main sorting configurations of the Steinberg experiments that emerge from an initially perturbed stationary state, as in [2, 18, 11] among others.

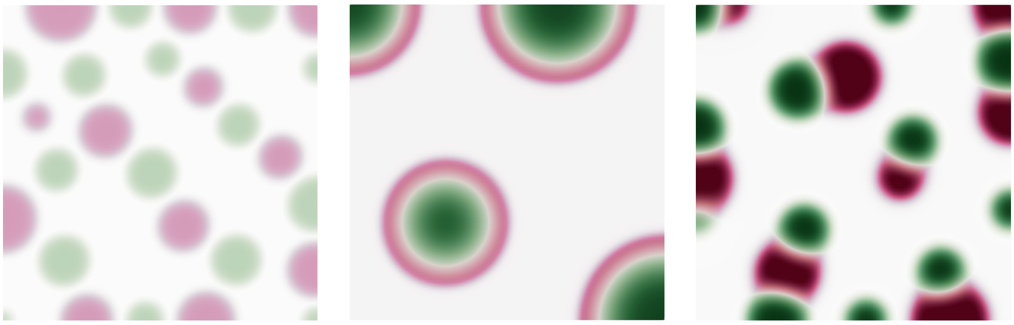

**(a)** Segregation **(b)** Engulfment **(c)** Partial sorting

**Figure 4: Sorting experiments.** Three sorting patterns that emerge from a random perturbation of a uniform state. (a) Segregation with $\xi_{DD} = \xi_{PP}$, $\xi_{DP} = 0$ and a purely repulsive potential $W(\rho) \propto \rho^4$. (b) Engulfment when $\xi_{PP} < \xi_{DP} < \xi_{DD}$ (c) Partial sorting when $\xi_{DP} < \xi_{PP} < \xi_{DD}$.

### 7 One-population with anisotropic diffusion: fingering instabilities

#### 7.1 Setting

In this section we consider the reverse problem with only one population but $\eta > 0$. In this section only, we rescale the time by $\varepsilon$ to obtain the following equation (with all the other coefficients equal to 1).

$$\partial_t \rho = \nabla \cdot (\rho \nabla q^\varepsilon) + \varepsilon^{-1}\eta^2 \nabla \cdot \big(\rho D(w) \nabla \rho\big), \tag{18a}$$

$$q^\varepsilon = \varepsilon^{-1} W'(\rho) - \varepsilon \Delta \rho \tag{18b}$$

With $\eta = 0$, we recover the standard Cahn-Hilliard equation for which we expect the existence of smooth radially symmetric solutions (centered at zero by convention) that partition the space between two domains $\{\rho = 0\}$ and $\{\rho = 1\}$ (corresponding to the stable wells of $W$) with an interface of typical size $\varepsilon$. The goal is to investigate numerically the stability of these solutions depending on $\eta$ and the total size of the aggregate, defined by $\overline{N} = \pi^{-1}\varepsilon^{-2}\int \rho$. This latter definition is taken by analogy with the agent-based model, here $\rho$ should be regarded as a phase-field corresponding to an aggregate of $\overline{N}$ spherical cells each with radius $\varepsilon$.

The initial configuration is a small random perturbation of the following custom smooth radially symmetric phase-field:

$$\rho(t=0,x) = \frac{\exp(-\frac{1}{R-r})\mathbb{1}_{r<R}}{\exp(-\frac{1}{R-r})\mathbb{1}_{r<R} + \exp(-\frac{1}{\varepsilon-(R-r)})\mathbb{1}_{r>R-\varepsilon}}, \quad r = |x|, \tag{19}$$

where $R = \varepsilon\sqrt{\overline{N}}$ is the radius of the disk of prescribed surface $\pi\overline{N}\varepsilon^2$.

In this toy setting, since only one population is considered, we do not implement any reaction-diffusion mechanism and simply assume that $w = x/|x|$ is constant in time and given. Note however, that since the aggregate is centered at zero, the anisotropic diffusion matrix $D(w)$ still preserves the spherical symmetry.

Then, for various values of $\eta$ and $\overline{N}$, we fix a (long) interval of time $T = 10$ and run the simulation until $T$ or until some instabilities have developed (see below for more details on how we choose the actual endpoint of the simulation).

#### 7.2 Results and discussion

An excerpt of representative results are shown in Fig.(M) S7B. When $\eta = 0$, the spherical aggregate remains stable with no visible significant deformation. However, when $\eta > 0$ is large enough, the anisotropic diffusion term is responsible for the symmetry breaking that takes the form of fingering-like instability.

Fingering instabilities are actually a well-known feature of systems involving Cahn-Hilliard type equations. The most classical and important example is the so-called Saffman-Taylor instability that describes the interface between fluids with different viscosity. In that context, instabilities typically develop after the injection of one of the fluids and/or due to fundamental principles such as gravity or Darcy law, that together induce the displacement of the interface. Such phenomena are often experimentally studied using so-called Hele-Shaw cells and modeled using phase-fields, and specifically Cahn-Hilliard equations, coupled to fluid mechanics equations, see for instance [25, 8, 23, 17] for some historical examples. Recently, the same heuristic has been applied to biological systems leading to novel models of tumor and tissue growth [21, 16, 9, 30]. Our framework and modeling assumptions may appear simpler (there is no coupling with fluid mechanics, mass is conserved etc) but all subsequent models are nevertheless derived from fundamental principles. We have already shown that this framework is sufficient to reproduce the formation of finger-like structures in the agent-based model and the present section shows that the same hold for the continuum model (18). This equation essentially reduces all the previous derivation to a Cahn-Hilliard term plus a novel anisotropic diffusion operator which models the motion of polarized cells. This latter term is the main destabilizing factor as shown below. In the following section we will simulate the complete system (15)-(16) and will observe a similar fingering behavior, thus suggesting that the reduced model (18) already captures the main driver of the emergence of fingers.

In conclusion, by analogy to the aforementioned literature on fluid instability, all this eventually points towards the conclusion that the formation of biological fingers is, ironically, a form of fingering instability.

##### 7.3 Fourier analysis

As in the biological experiments shown in the main text, the number of fingers is linked to the size of the aggregate $\overline{N}$. To evaluate this phenomenon more quantitatively and in order to automatically assess whether some instability has developed, we use the automatic contouring tool provided by [28] and we compute the Fourier decomposition of the contour seen as a periodic function. More precisely, for any integer $k$, we compute the $k$-th Fourier coefficient

$$c_k = \int_0^1 r(\theta)\exp(-2i\pi k\theta)\mathrm{d}\theta,$$

where $\theta$ is the polar angle measured from the center of the aggregate and $r(\theta)$ is the distance between the contour point and this center. This way, a non-zero $k$-th Fourier mode exactly corresponds to the emergence of $k$ fingers. The zero-th Fourier coefficient $c_0$ is equal to the radius of the aggregate. We stop the simulation when one Fourier coefficients becomes larger than 5% of $c_0$. The two figures Figs.(M) S7D and S7E show which Fourier mode becomes unstable for a given $\eta = 0.6$ and various $\overline{N}$ and the value of the largest Fourier coefficient as a function of $\eta$ for various $\overline{N}$.

From Fig.(M) S7D we conclude that one Fourier mode seems to always be dominating. This unstable mode grows more or less linearly as a function of $\overline{N}$. This observation is consistent with the experimental results that show that the number of emerging fingers is a growing function of the size of the organoid. The next figure Fig.(M) S7E indicates the value of $\eta$ required to break symmetry. Interestingly, this value does not seem to clearly depend on $\overline{N}$. Instead, symmetry is always broken for $\eta > 0.4$ and unstable modes develop when $\eta > 0.2$. This may point towards a phase transition phenomenon triggered solely by the polarization parameter $\eta$. This preliminary numerical analysis could serve as a starting point for a more thorough and mathematical analysis of the simplified model (18) that is left for future work.

### 8 Two populations with traction bias: organoid experiments

Finally we simulate the full system (15) with Distal and Proximal cells coupled to the reaction-diffusion network (16). We initialize the simulation with a uniformly mixed randomly perturbed spherical aggregate. As in the particle simulations, we compare the influence of the traction bias $\eta$ versus the differential adhesion ratio $\alpha = \xi_{DD}/\xi_{PP}$. We assume $\alpha \geq 1$ and set $\xi_{DP} = \xi_{PP} + \frac{1}{3}(\xi_{DD} - \xi_{PP})$. We show the results after an arbitrary time $T = 42$ where equilibrium is reached or until instabilities have developed. Both the ABM and PDE models show that instabilities develop when both the traction bias and the differential adhesion are strong enough. When one of these two conditions fail, we observe different types of sorting patterns. In particular Distal cells tend to be internalized due to differential adhesion and, on the contrary, to cluster at the periphery of the aggregate due to the traction bias (because the morphogen gradient is pointing outward the aggregate). The combination of these two antagonistic effects produces in both models the elongation of finger-like patterns starting from buds of Distal cells. We have also checked that the number of fingers increases with the size of the aggregate and that Distal only aggregates do not produce fingers as in the ABM and the organoid experiments. The qualitative agreement between the ABM and the PDE simulations also legitimate the PDE derivation. It points towards the claim that the main driver of finger formation can indeed be encoded in the combination of a Cahn-Hilliard operator with a destabilizing term, here modeled by anisotropic diffusion. Yet, we mention and comment below a few noticeable differences between the ABM and the PDE models that we have observed in the simulations.

- In the ABM simulations, the adhesion forces are strong enough to maintain the coherence of the aggregate. In the PDE simulations, the fingers keep elongating and may eventually dissociate from the main aggregate. Although this long-time behavior is not what would be expected biologically, we note that the focus of this model is rather the onset of the instabilities than their later development.

- On a similar note, when the adhesion is too weak, the diffusion terms make the cell density diffuse in the medium rather than maintain a dense aggregate shape. On the contrary, particle simulations do not suffer from this non-compact support issue, as already noticed in similar models of cell aggregation phenomena [21, 18].

- The range of admissible parameters in the PDE system still remains to be explored. The ABM is defined by a set of ordinary differential equations with regular terms and thus remains well-defined for all the physically meaningful parameters. On the contrary, PDE systems are typically not expected to be well-posed for any parameter set. While simulations show that the PDE is apparently well-posed in a region of interest of the parameter space, issues may happen outside. These issues may be both of theoretical or numerical nature.

### 9 Discussion and open mathematical problems

The derivation of the various PDE models from an ABM proceeds with several steps that are potentially mathematically challenging to write in a fully rigorous manner. On the one hand, the derivation of mean-field models is by now a well-established approach with many rigorous convergence results available [6, 7]. On the other hand, the localization limit that leads to the main PDE model seems more challenging and an ad hoc approach would be desirable to make rigorous the formal Taylor expansion presented here. Well-posedness conditions and an appropriate notion of solution for the equation (14) are likely to play an important role in this task. While similar results have been rigorously established for related models [10], such mathematical analysis remains to be adapted to the present setting. Along with this theoretical analysis, another important challenge is to design custom numerical schemes for the equation (15) and to establish the stability and convergence results that would properly justify the comparison with the ABM and the organoid experiments. These questions have recently been addressed in [13, 14] following the pre-publication of a first version of our model.

Here, we have numerically explored the instability properties of the novel PDE model (15). Although it clearly appears that the additional anisotropic diffusion term is the main factor that explains the formation of finger-like patters, such observation remains to be proved rigorously, analogously to what is known on diffusion-driven instability in Turing systems for instance. However, a major difficulty is that we would have to carry a (linear) perturbation analysis around a non-constant solution that is itself not clearly characterized. In the present case, the existence of radially symmetric solutions would need to be established first. We could also think of different geometrical configurations to reduce the complexity. As shown earlier, the emergence of finger-like patterns does not seem to require two populations, since similar patterns can emerge in a one-population Cahn-Hilliard model supplemented with a diffusion term. This simpler model could serve as a starting point for the instability analysis. It is also much more related to well-known models of viscous fluids, with a direct analogy to fingering instabilities in the Hele-Shaw cell discussed earlier.

In the present work, we have stopped our investigations to the PDE model (15). A natural follow-up question in this context is to investigate the so-called sharp interface limit $\varepsilon \to 0$. It is well-known that for the Cahn-Hilliard equation, this procedure leads to the study of free-boundary problems, in particular here, of the so-called mean-curvature flow [24, 1, 19, 4]. Such models consider only the dynamic of the interface (in our case, between the cells and the medium or between the two cell populations) and thus provides a much more geometrical description of the dynamics, potentially also better prone to an instability analysis. To the best of our knowledge, the derivation of sharp interface limits with an additional anisotropic diffusion term is largely open, with the exception of [15] that proves such a result but when the Cahn-Hilliard operator is replaced by the simpler Allen-Cahn operator. The case of two populations has not been explored.

### 10 Parameter values

#### 10.1 Summary table of the ABM parameters

The Tables 1 and 2 summarizes all the parameters of the ABM, their meanings and numerical values. When a value is indicated, it is kept constant in all the simulations presented. The values indicated as *variable* depend on the numerical experiment and are thus given in the corresponding part of the text. The considered choices are shown with default values in bold. The code repository `https://github.com/antoinediez/PyLimbMorph` contains scripts showing parameters scans with defined bounds.

| Symbol | Meaning | Value |
|---|---|---|
| $N$ | Total number of cells | *variable* $\{6, \mathbf{10}, 12, 18, 24\} \times 10^3$ |
| $\overline{N}_P$ | Ratio of proximal cells | *variable* $\{0, 0.25, 0.5, \mathbf{0.6\overline{6}}, 0.75, 1\}$ |
| $N_P = \overline{N}_P N$, $N_D = N - N_P$ | Cells per cell type | (see above) |
| $T$ | Terminal time | *variable* $\{\mathbf{600}, 1200, 1800\}$ |
| $\alpha$ | Relative attraction strength | *variable* $\{1, 2, \mathbf{3}\}$ |
| $\xi_{DD}, \xi_{DP}, \xi_{PP}$ | Attraction strength | $3$, $3 \cdot \frac{1+\alpha}{2}$, $3\alpha$ |
| $\eta_D^{DD}, \eta_D^{DP}$, $\eta_P^{PP}, \eta_P^{DP}$ | Traction bias | *variable* $\{0, 0.25, 0.5, \mathbf{0.75}, 1\}$ |
| $R$ | Attraction radius | $3\delta$ |
| $\delta$ | Repulsion radius | $0.3125/N_0^{1/3}$ with $N_0 = 10^4$ |
| $F_0^{\text{rep}}$ | Repulsion strength | 10 |
| $\lambda_u, \lambda_v$ | Decay rates | 35, 10 |
| $\mu_u, \mu_v$ | Production rates | 5, 0 |
| $D_u, D_v$ | Diffusion rates | 0.35, 1 |

**Table 1:** Modeling parameters of the ABM (with physical meaning).

| Symbol | Meaning | Value |
|---|---|---|
| $M$ | Number of discrete grid-cells per dimension | 64 |
| $\sigma$ | Smoothening parameter | $1/M$ |
| $h_-, h_+, \phi_0$ ($h_{\text{in}}(\phi_D)$ Wnt5a) | Minimal, maximal and thresholding values for the phase-fields | $(0, 1, 0.3)$ |
| $h_-, h_+, \phi_0$ ($h_{\text{in}}(\phi_{\text{cells}})$ Wnt5a) | Minimal, maximal and thresholding values for the phase-fields | $(0, 1, 1)$ |
| $h_-, h_+, \phi_0$ ($h_{\text{in}}(\phi_{\text{cells}})$ Fgf8) | Minimal, maximal and thresholding values for the phase-fields | $(1, 1, 1)$ |
| $h_-, h_+, \phi_0$ ($h_{\text{out}}(\phi_{\text{cells}})$ Fgf8) | Minimal, maximal and thresholding values for the phase-fields | $(0.01, 1, 0.5)$ |
| $g_0$ | Thresholding value for the gradient of Wnt5a | 0.1 |

**Table 2:** Technical parameters of the ABM.

##### 10.2 Summary table of the PDE parameters

| Symbol | Meaning |
|---|---|
| $\xi_{DD}, \xi_{DP}, \xi_{PP}$ | Attraction strength |
| $\eta$ | traction bias |
| $\lambda$ | Phase separation strength |
| $\varepsilon$ | Scaling size |
| $\rho_0$ | Thresholding density |
| $c_0$ | Advection speed |
| $c_1$ | Cell-cell attraction-repulsion strength |
| $c_2$ | anisotropic diffusion strength |
| $c_3$ | Interface smoothening |
| $c_4$ | Correction strength due to the differential adhesion |
| $\nu$ | Linear diffusion coefficient of the cells |
| $\lambda_u, \lambda_v$ | Decay rates |
| $\mu_u, \mu_v$ | Production rates |
| $D_u, D_v$ | Diffusion rates of the morphogens |
| $g_0$ | Thresholding value for the gradient of Wnt5a |
| $L$ | Size of the domain (radius in a disk and edge length in a square) |
| $\overline{N}$ | Size of the aggregate |
| $\bar{N}_D$ | Ratio of Distal versus Proximal cells |
| $\Delta t$ | Time step |
| $\Delta x$ | Mesh size |

**Table 3:** Parameters for the continuum model (15)-(16)

##### 10.3 Sorting experiments

Initial condition (up to some random perturbation):

$$\text{(a)} \quad \rho_D(t=0) = \rho_P(t=0) = 0.1$$

$$\text{(b,c)} \quad \rho_D(t=0) = \rho_P(t=0) = 0.2$$

| Symbol | Meaning | Value |
|---|---|---|
| $\xi_{DD}, \xi_{DP}, \xi_{PP}$ | Attraction strength | (a) 20, 0, 20<br>(b) 10, 1, 0.5<br>(c) 12, 1, 6 |
| $\eta$ | traction bias | 0 |
| $\lambda$ | Phase separation strength | 10 |
| $\varepsilon$ | Scaling size | (a) 0.02 (b,c) 0.1 |
| $\rho_0$ | Thresholding density | $\xi_{DP}$ |
| $c_0$ | Advection speed | 0 |
| $c_1$ | Cell-cell attraction-repulsion strength | 1 |
| $c_2$ | anisotropic diffusion strength | 0 |
| $c_3$ | Interface smoothening | 1 |
| $c_4$ | Correction strength due to the differential adhesion | 1 |
| $\nu$ | Linear diffusion coefficient of the cells | 1 |
| $L$ | Size of the domain $[0, L]^2$ | (a,b) 2 (c) 4 |
| $\bar{N}_D$ | Ratio of Distal versus Proximal cells | 0.5 |
| $\Delta t$ | Time step | $5 \cdot 10^{-4}$ |
| $\Delta x$ | Mesh size | $L/100$ |

**Table 4:** Parameters for Eq. 17 in Fig. 4

#### 10.4 Fingering instability

Initial condition: (19).

| Symbol | Meaning | Value |
|---|---|---|
| $\eta$ | traction bias | In the range $(0, 1)$ |
| $\lambda$ | Phase separation strength | $1/\varepsilon$ |
| $\varepsilon$ | Scaling size | 0.1 |
| $\rho_0$ | Thresholding density | 1 |
| $L$ | Radius of the disk domain | $100R$ |
| $\overline{N}$ | Size of the aggregate | In the range $(200, 2000)$ |
| $\bar{N}_D$ | Ratio of Distal versus Proximal cells | 1 |
| $\Delta t$ | Time step | $5 \cdot 10^{-4}$ |
| $\Delta x$ | Mesh size | $1.5\varepsilon$ |

**Table 5:** Parameters for Eq. (18) in Figs.(M) S7B, S7D, S7E.

#### 10.5 Two-population

Initial condition: (19) multiplied respectively by $\bar{N}_D$ and $1 - \bar{N}_D$ for Distal and Proximal cells.

| Symbol | Meaning | Value |
|---|---|---|
| $\xi_{DD}, \xi_{DP}, \xi_{PP}$ | Attraction strength | $\alpha$,$(2+\alpha)/3$,1 |
| $\eta$ | traction bias | In the range $(0,1)$ |
| $\lambda$ | Phase separation strength | 3 |
| $\varepsilon$ | Scaling size | 0.1 |
| $\rho_0$ | Thresholding density | 1 |
| $c_0$ | Advection speed | 0.1 |
| $c_1$ | Cell-cell attraction-repulsion strength | 1.5 |
| $c_2$ | anisotropic diffusion strength | 1 |
| $c_3$ | Interface smoothening | 1.5 |
| $c_4$ | Correction strength due to the differential adhesion | 1 |
| $\nu$ | Linear diffusion coefficient of the cells | 0.2 |
| $\lambda_u$, $\lambda_v$ | Decay rates | 0.8,1 |
| $\mu_u$, $\mu_v$ | Production rates | 1.2,1.2 |
| $D_u$, $D_v$ | Diffusion rates of the morphogens | 0.3,1.2 |
| $g_0$ | Thresholding value for the gradient of Wnt5a | 0.01 |
| $L$ | Radius of the disk domain | $20R$ |
| $\overline{N}$ | Size of the aggregate | 800 |
| $\bar{N}_D$ | Ratio of Distal versus Proximal cells | 0.35 |
| $\Delta t$ | Time step | $10^{-2}$ |
| $\Delta x$ | Mesh size | 0.05 |

**Table 6:** Parameters for Eqs. 15-16 in Fig.(M) S7A
