## Supplemental Data 1 for "Self-organized mechanochemical instabilities drive the emergence of digit tissue morphogenesis"

### Estimation of the Cauchy-Green strain tensor from cell tracks

#### 1 Cauchy-Green strain tensor analysis

Let us denote the tracking data as $(\boldsymbol{X}_i(t), \boldsymbol{Y}_i(t+1)) \in \mathbb{R}^3 \times \mathbb{R}^3$ where $t = 0, \ldots, M-1$ denotes the current frame, $i = 1, \ldots, N(t)$ is the label for a spot pair for which $\boldsymbol{X}_i(t)$ is the cell position at frame $t$ and $\boldsymbol{Y}_i(t+1)$ the position of the same cell at the next frame.[1] Forward velocities are defined as

$$\boldsymbol{V}_i(t) := \boldsymbol{Y}_i(t+1) - \boldsymbol{X}_i(t).$$

We denote the coordinates $x, y, z$ by $x(\boldsymbol{X}_i(t)), y(\boldsymbol{X}_i(t)), z(\boldsymbol{X}_i(t))$ to avoid adding another index.

##### 1.1 Choice of a coordinate system

The spot dataset shows only a part of the upper half of the roughly spherical organoid. We manually rotated the positions with a rotation around the $z$ -axis, so that the $x$ -axis aligns with the elongation direction. We translate and scale the coordinates so that

1. $\sum_i x(\boldsymbol{X}_i(t)) = \sum_i y(\boldsymbol{X}_i(t)) = 0 \quad \forall t,$
2. $\min_{i,t} z(\boldsymbol{X}_i(t)) = 0.5$
3. $\max_{i,t} \|\boldsymbol{X}_i(t) - (0.5, 0.5, 0.5)^T\|_\infty = 0.5.$

[1]Note that this setup is required as infered cell tracking data has gaps and the number of cells can vary between frames.

#### 1.2 Removal of outside tracks

The tracking datasets contain tracks which belong to spots outside of the core organoid. We remove most of these tracks as follows:

1. We have manually annotated the axis-aligned bounding box $\Omega(t_k)$ of the organoid for certain frames $t_k$; for in-between frames we use linear interpolation to obtain an approximate bounding box $\Omega(t)$.

2. Then, we remove all spots where $\boldsymbol{X}_i(t)$ or $\boldsymbol{Y}_i(t+1)$ are outside of the bounding box.

#### 1.3 Interpolated vector field via Gaussian kernel mollifier

We define a smooth continuous vector field at frame $t = 0, \ldots, M-1$ via

$$\begin{aligned} w(t,\boldsymbol{x}) &:= \frac{1}{(2\pi)^{3/2}\sigma^3 N(t)} \sum_{i=1}^{N(t)} \mathrm{e}^{-\frac{\|\boldsymbol{x}-\boldsymbol{X}_i(t)\|^2}{2\sigma^2}}, \\ \boldsymbol{v}(t,\boldsymbol{x}) &:= \frac{1}{(2\pi)^{3/2}\sigma^3 N(t)(\beta + w(t,\boldsymbol{x}))} \sum_{i=1}^{N(t)} \boldsymbol{V}_i \mathrm{e}^{-\frac{\|\boldsymbol{x}-\boldsymbol{X}_i(t)\|^2}{2\sigma^2}} \end{aligned}$$

where $\sigma$ denotes a smoothing radius, which should be on a scale comparable to the typical distance between cell tracks, $\beta$ is a bias parameter to ensure that, in the absence of data when $w \ll 1$, the velocity is zero.

For $t \in (0, M-1)$ we define the weights and the vector field via linear interpolation

$$\begin{aligned} w(t,\boldsymbol{x}) &= (\lceil t \rceil - t) w(\lfloor t \rfloor, \boldsymbol{x}) + (t - \lfloor t \rfloor) w(\lceil t \rceil, \boldsymbol{x}), \\ \boldsymbol{v}(t,\boldsymbol{x}) &= (\lceil t \rceil - t) \boldsymbol{v}(\lfloor t \rfloor, \boldsymbol{x}) + (t - \lfloor t \rfloor) \boldsymbol{v}(\lceil t \rceil, \boldsymbol{x}). \end{aligned}$$

#### 1.4 Deformation map and Cauchy-Green strain tensor

Let us denote the deformation map $\varphi_t : \mathbb{R}^3_{t_0} \to \mathbb{R}^3_t$ as the flow of the differential equation $\dot{\boldsymbol{x}} = \boldsymbol{v}(\boldsymbol{x})$, which means that it satisfies for all $\boldsymbol{x}_0 \in \mathbb{R}^3_{t_0}$ at initial time $t_0$. We write the indices to make the time explicit to differentiate the reference space from the dynamic coordinates. The differential equation is given by

$$\begin{aligned} \partial_t \varphi_t(\boldsymbol{x}_0) &= \boldsymbol{v}(\varphi_t(\boldsymbol{x}_\mathbf{0})) \quad \forall t \in (0, M-1), \\ \varphi_0(\boldsymbol{x}_0) &= \boldsymbol{x}_\mathbf{0}. \end{aligned}$$

The deformation gradient is defined as

$$F(\boldsymbol{x}) := \partial_{\boldsymbol{x}} \varphi_t(\boldsymbol{x}) \in \mathbb{R}^{3\times 3}$$

which maps from $T\mathbb{R}^3_{t_0} \to T\mathbb{R}^3_t$, e.g. from a tangent vector with respect to the reference coordinates to a tangent in the world coordinates. As these two coordinates systems are in general unrelated, it is useful to instead consider the Cauchy-Green tensor which defines a metric on one tangent space (either reference or spatial coordinates), we hence introduce

$$C(\boldsymbol{x}) := F^T(\boldsymbol{x})F(\boldsymbol{x}) \in T^2\mathbb{R}^3_{t_0}, \quad (1)$$

$$B(\boldsymbol{x}) := F(\boldsymbol{x})F^T(\boldsymbol{x}) \in T^2\mathbb{R}^3_t \quad (2)$$

which are the Cauchy-Green strain tensor and the Green strain tensor (also called left Cauchy-Green strain tensor). The square root of these tensors define metrics on $T\mathbb{R}^3_{t_0}$ and $T\mathbb{R}^3_t$ respectively.

#### 1.5 Measure of reliability and visualizations

Now, for a given point $\boldsymbol{x} \in \mathbb{R}^3_{t_0}$ we can visualize

$$\mathbb{R}^3_{t_0} \to T^2\mathbb{R}^3_{t_0} : \boldsymbol{x} \mapsto \sqrt{C(\boldsymbol{x})}$$

by plotting

$$\mathbb{R}^3_{t_0} \to T^2\mathbb{R}^3_{t_0} : X \mapsto \sqrt{C(\boldsymbol{x})}.$$

Alternatively, we can visualize in spatial coordinates as

$$\mathbb{R}^3_t \to T^2\mathbb{R}^3_t : \boldsymbol{X} \mapsto \sqrt{B(\varphi_t^{-1}(\boldsymbol{X}))}.$$

*Currently, we visualize the deformation tensors in spatial coordinates (B), since they align visually better with the cell tracks.*

#### 1.6 Axes and index of convergent-extension

Visualizing tensors is a bit tedious; therefore, we would like to represent the major convergent-extension direction and strength as a nematic vector.

Let us fix a location in spatial coordinates $\boldsymbol{x} \in \mathbb{R}^3_t$ and a time $t \in (0, M-1)$. We consider the eigenvalues $\lambda_1 \leq \lambda_2 \leq \lambda_3$ of $B(t, \boldsymbol{x})$ and denote the principal eigenvector (corresponding to $\lambda_3$) as $\boldsymbol{u}$. Note that the eigenvalues of $B$ and $C$ coincide, but the eigenvectors of $B$ live in spatial coordinates $T\mathbb{R}^3_t$, which is where we perform the visualization.

We define the extension ratio

$$\alpha := \frac{\lambda_3}{\sqrt{\lambda_1 \lambda_2}}$$

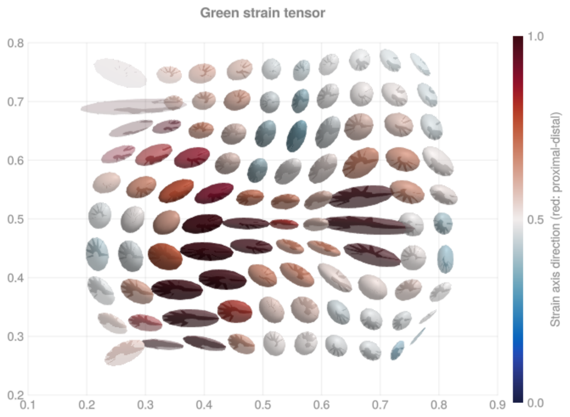


**Figure 1:** Green strain tensor $\sqrt{B}$.

which equals 1 for isotropic deformation and grows for anisotropic deformation along the principal axis. In other terms, $\alpha$ is large for convergent-extension like deformations.

One remaining issue is that we cannot be certain of how reliable the information is at each given point, since there might be only very few cell tracks close to the location $\boldsymbol{x}$. Here we can use the previously defined weights $w(t, \boldsymbol{x})$.

Combined, we obtain the following nematic vector

$$\boldsymbol{p}(t, \boldsymbol{x}) := \frac{1}{c} \frac{\lambda_3(\boldsymbol{x})}{\sqrt{\lambda_1(\boldsymbol{x})\lambda_2(\boldsymbol{x})}} w(t, \boldsymbol{x})\, \boldsymbol{u}(\boldsymbol{x}).$$

Here, $w(t, \boldsymbol{x})$ is the instantaneous weight from the kernel density estimate, which is large in regions with many nearby cell tracks and small in sparsely tracked regions. The normalization constant $c > 0$ is chosen as a global constant across all datasets so that the index is comparable between experiments. The vector $\boldsymbol{p}(t, \boldsymbol{x}) \in T\mathbb{R}^3_t$ is large if there are many cell tracks which together show convergent-extension like dynamics.

To finally answer if this convergent-extension like deformation happens along the proximaldistal axis (or orthogonal), we define the proximaldistal and orthogonal convergent-extension indices as

$$\mathcal{C}(t, \boldsymbol{x}) = |\langle \boldsymbol{p}(t, \boldsymbol{x}), \boldsymbol{e}_x \rangle|, \quad (3)$$

$$\mathcal{C}^{\perp}(t, \boldsymbol{x}) = \|\boldsymbol{p}(t, \boldsymbol{x})\| - |\langle \boldsymbol{p}(t, \boldsymbol{x}), \boldsymbol{e}_x \rangle|. \quad (4)$$

#### 1.7 CE index and cell density along proximaldistal center line

We manually annotate a line $L : [0, 1] \to \mathbb{R}^3_{t_0}$ which denotes a central line passing in the proximal-distal direction through the center of elongation, see Fig. 2.

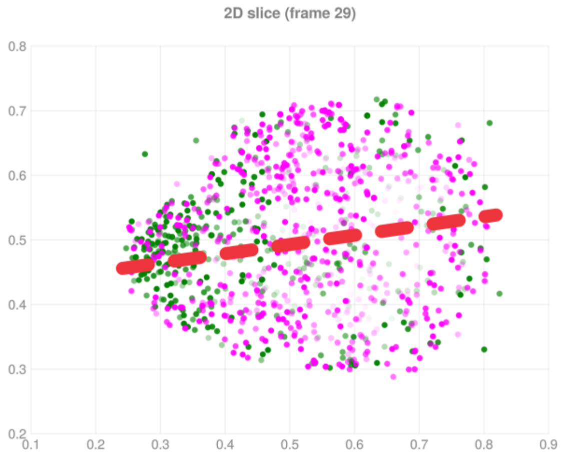


**Figure 2:** The line $L$ shows the apical-proximal axis, green and magenta spots show the tracking data.

Along this line, we plot

$$t \mapsto \mathcal{C}(t, L(t)) \quad \text{and} \quad t \mapsto \mathcal{C}^{\perp}(t, L(t)).$$

Moreover, we define the kernel density estimations of distal and proximal cells along the line as

$$\rho_D(\boldsymbol{x}) = \frac{1}{(2\pi)^{3/2}\sigma^3 N_D} \sum_{i=1}^{N_D} \mathrm{e}^{-\frac{\|\boldsymbol{x}-\boldsymbol{X}_i\|^2}{2\sigma^2}}, \tag{5}$$

$$\rho_P(\boldsymbol{x}) = \frac{1}{(2\pi)^{3/2}\sigma^3 N_P} \sum_{i=N_D+1}^{N_P} \mathrm{e}^{-\frac{\|\boldsymbol{x}-\boldsymbol{X}_i\|^2}{2\sigma^2}}. \tag{6}$$

Finally, this yields Fig. 3, which shows in particular the the convergence extension index peaks at the interface between distal and proximal cells.

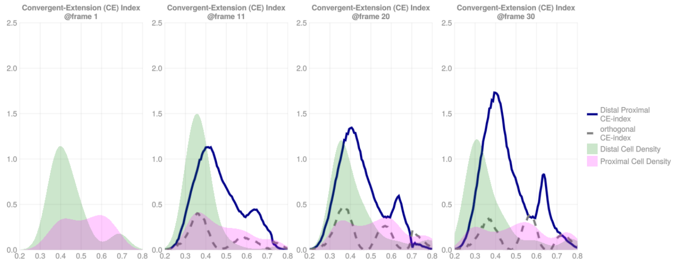


**Figure 3:** Plot of the convergent-extension index $\mathcal{C}$ and the kernel density estimates of the cell types along the line.
